# Identification of 38 novel loci for systemic lupus erythematosus and genetic heterogeneity that may underly population disparities in this disease

**DOI:** 10.1101/2020.04.12.037622

**Authors:** Yong-Fei Wang, Yan Zhang, Zhiming Lin, Huoru Zhang, Ting-You Wang, Yujie Cao, David L. Morris, Yujun Sheng, Xianyong Yin, Shi-Long Zhong, Xiaoqiong Gu, Yao Lei, Jing He, Qi Wu, Jiangshan Jane Shen, Jing Yang, Tai-Hing Lam, Jia-Huang Lin, Zhi-Ming Mai, Mengbiao Guo, Yuanjia Tang, Yanhui Chen, Qin Song, Bo Ban, Chi Chiu Mok, Yong Cui, Liangjing Lu, Nan Shen, Pak C. Sham, Chak Sing Lau, David K. Smith, Timothy J. Vyse, Xuejun Zhang, Yu Lung Lau, Wanling Yang

**Affiliations:** Department of Paediatrics and Adolescent Medicine, The University of Hong Kong; Department of Pediatric Surgery, Guangzhou Institute of Pediatrics, Guangdong Provincial Key Laboratory of Research in Structural Birth Defect Disease, Guangzhou Women and Children’s Medical Center, Guangzhou Medical University; Department of Rheumatology, The Third Affiliated Hospital of Sun Yat-Sen University; The Hormel Institute, University of Minnesota; Division of Genetics and Molecular Medicine, King’s College London; Department of Dermatology, No.1 Hospital, Anhui Medical University; Guangdong Provincial Key Laboratory of Coronary Heart Disease Prevention, Guangdong Provincial People’s Hospital; Department of Clinical Biological Resource Bank, Guangzhou Institute of Pediatrics, Guangzhou Women and Children’s Medical Center; School of Public Health, The University of Hong Kong; Radiation Epidemiology Branch, Division of Epidemiology and Genetics, National Cancer Institute, National Institutes of Health; Shanghai Institute of Rheumatology, Renji Hospital, Shanghai Jiao Tong University School of Medicine; Department of Pediatrics, Union Hospital Affiliated to Fujian Medical University; Department of Rheumatology, Affiliated Hospital of Jining Medical University; Department of Endocrinology, Affiliated Hospital of Jining Medical University; Department of Medicine, Tuen Mun Hospital; Department of Dermatology, China-Japan Friendship Hospital; Department of Psychiatry, The University of Hong Kong; Department of Medicine, The University of Hong Kong

**Author notes:** These authors contributed equally (Yong-Fei Wang, Yan Zhang). Correspondence to: WY (W.Y.) or YLL (Y.L.L.).

## Abstract

Systemic lupus erythematosus (SLE), a worldwide autoimmune disease with high heritability, shows differences in prevalence, severity and age of onset among different ancestral groups. Previous genetic studies have focused more on European populations, which appear to be the least affected. Consequently, the genetic variations that underly the commonalities, differences and treatment options in SLE among ancestral groups have not been well elucidated. To address this, we undertook a genome-wide association study, increasing the sample size of Chinese populations to the level of existing European studies. Thirty-eight novel SLE-associated loci and incomplete sharing of genetic architecture were identified. Nine disease loci showed clear ancestral group heterogeneity and implicated antibody production as a potential mechanism for differences in disease manifestation. Polygenic risk scores performed significantly better when trained on matched ancestral data sets. These analyses help to reveal the genetic bases for disparities in SLE among ancestral groups.

## Introduction

Systemic lupus erythematosus (SLE; OMIM 152700) is an autoimmune disease characterized by production of autoantibodies and multiple organ damage. Genetic factors play a key role in the disease, with estimates of its heritability ranging from 43% to 66% across populations^1–3^. Differences in the expression of the disease across ancestral groups have been reported with non-European populations showing an earlier age of onset, 2-4 fold higher prevalence and 2-8 fold higher risk of developing end-stage renal disease than European populations^4–7^. Responses to treatment of SLE with the novel monoclonal antibody against B-cell activating factor (BAFF), Belimumab, also show variation across ancestral groups^8,9^. These findings highlight the heterogeneous nature of the disease, so closer examination of ancestral group differences is likely to improve disease risk prediction and lead to more precise treatment options.

More than 90 loci have been shown to be associated with SLE through genome-wide association studies (GWAS)^10–12^. Trans-ancestral group studies conducted previously were primarily designed to increase power and to identify SLE susceptibility loci shared across ancestries^13,14^. However, due to inadequate power in studies involving non-Europeans, current findings are biased towards loci associated with SLE in European populations. Some risk alleles reported from studies on European populations, such as those in or near *PTPN22, NCF2, SH2B3* and *TNFSF13B*, are absent in East Asian populations^15^ while a missense variant (rs2304256) in *TYK2* points to a European-specific disease association^16–18^.

The basis for ancestral group differences in the manifestation of SLE at the genome level remains poorly understood. Further studies on non-European populations will help define the genetic architecture underlying SLE and the consequences of patients’ ancestral backgrounds. To this end, we genotyped 8,252 participants of Han Chinese descent recruited from Hong Kong (HK), Guangzhou (GZ) and Central China (CC), and combined these data with previous datasets to give a total of ten SLE genetic cohorts consisting of 11,283 cases and 24,086 controls. The increased sample size, particularly for those of Chinese ancestry, allowed identification of novel disease loci and comparative analyses of the genetic architectures of SLE between major ancestral groups. Functional annotation of these findings provided insights into ancestral group differences in SLE.

## Results

### Data set preparation

Han Chinese data: After removing individuals with a low genotyping rate or hidden relatedness, the 7,596 subjects of Han Chinese descent from HK, GZ and CC genotyped in this study and the 5,057 subjects from the existing Chinese GWAS^13^ gave a Chinese ancestry data set of 4,222 SLE cases and 8,431 controls (**Extended Data Table 2**). Ethnic European Data: Existing GWAS data from European populations^19^ were reanalyzed, based on principal components (PC) matching those for subjects from the 1,000 Genomes Project to minimize the potential influence of population substructures^20^ (**Methods**) and grouped into three cohorts, EUR GWAS 1-3, (**Extended Data Fig. 2**). The recent SP GWAS^21^ data was included. After quality control, the European data included 4,576 cases and 8,039 controls. A further 2,485 SLE cases and 7,616 controls were included as summary statistics from an Immunochip study of East Asians^22^ (**Extended Data Table 2**).

### Ancestral correlation of SLE

Genotype imputation and association analysis were performed independently for each GWAS cohort and as meta-analyses of each ancestral group (**Methods**). The trans-ancestral genetic-effect correlation, r_ge_, between the Chinese and European GWAS was estimated to be 0.64 with a 95% confidence interval (CI) of 0.46 to 0.81 by Popcorn^23^ (**Methods**), indicating a significant, but incomplete, correlation of the genetic factors for SLE between the two ancestries.

### Novel SLE susceptibility loci

Meta-analyses, involving a total of 35,369 participants (**Methods**; **Extended Data Table 2**) were conducted. Of the 94 previously reported SLE associated variants (**Extended Data Table 3**), 59 (62.8%) surpassed a genome-wide significance *P-*value threshold (5.0E-08) and 84 (89.4%) exceeded the threshold of 5E-05 in our study. Thirty-four novel variants reached genome-wide significance and four variants had *P-* values approaching this threshold based on either ancestry-dependent or trans-ancestral meta-analyses. The newly identified loci included the immune checkpoint receptor *CTLA4*, the TNF receptor-associated factor *TRAF3* and the type I interferon gene cluster on 9p21 (**Table 1** and **Extended Data Table 4**). The new loci bring the total of SLE-associated loci to 132 and produce a 23.5% and 16.5% increase in heritability explained for East Asians and Europeans, respectively (**Methods**).

**Table 1.**
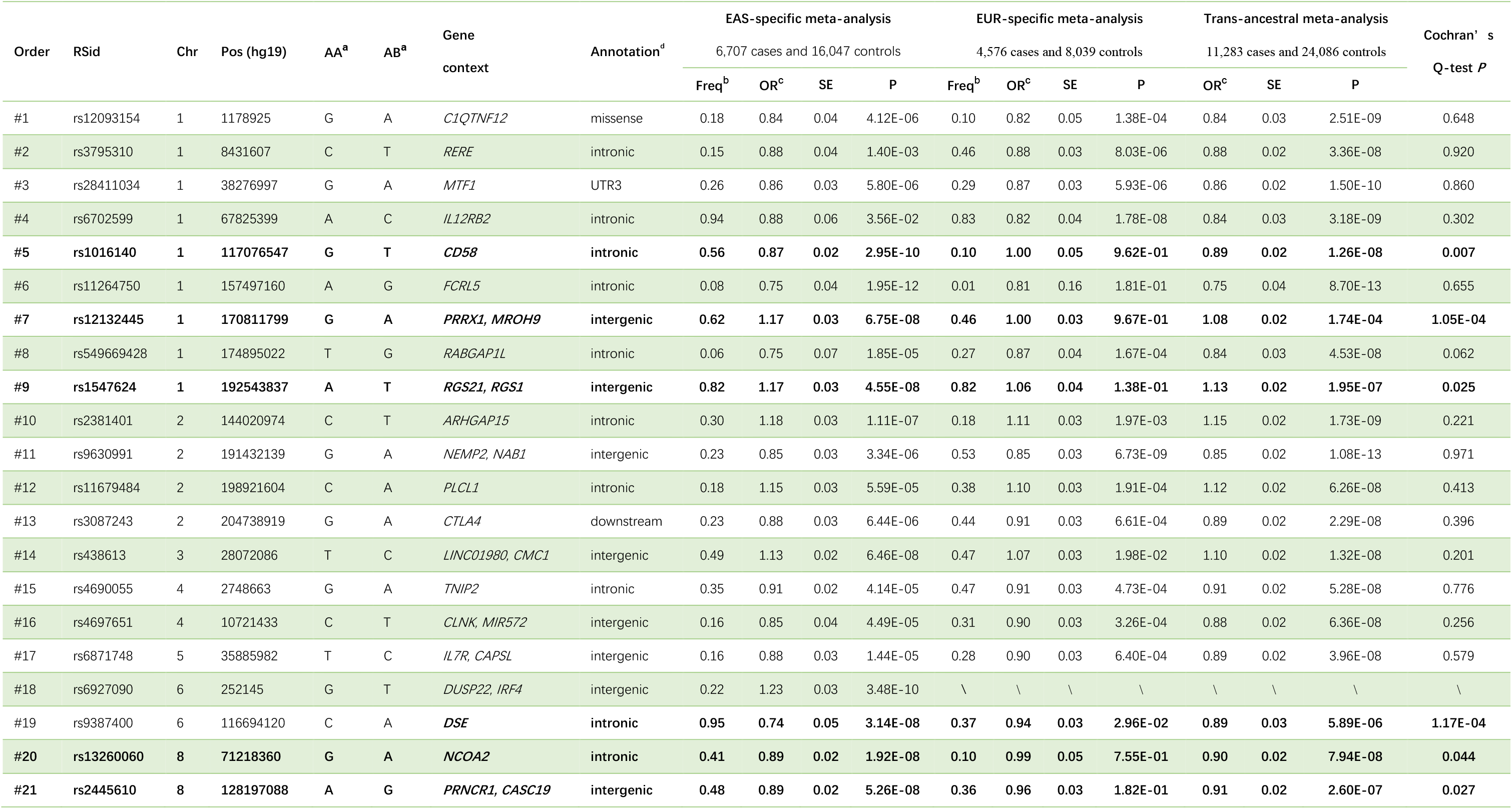
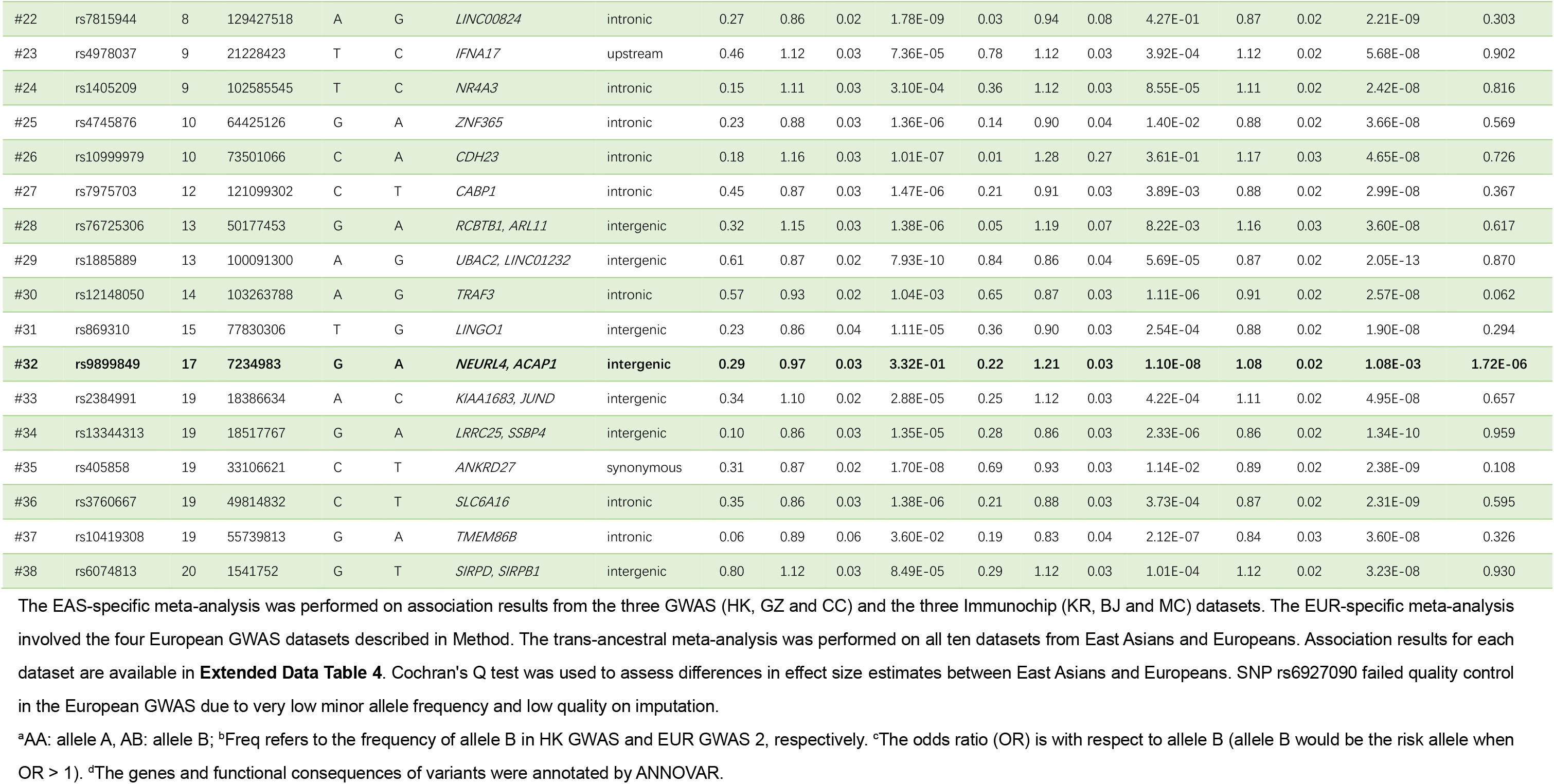
Summary association statistics of newly identified SLE-associated variants

### Annotation of SLE susceptibility loci

Functional annotations that might be enriched with SLE susceptibility loci were evaluated by the stratified LD score regression method^24^ (**Methods**). For non-cell type-specific annotations, heritability was significantly enriched in transcription start sites (*P* = 2.47E-05), regions that are conserved in mammals (*P* = 8.22E-03) and ubiquitous enhancers that are marked with H3K27ac or H3K4me1 modifications (*P* = 4.43E-02, *P* = 5.03E-02, respectively; **Extended Data Fig. 5**). Based on H3K4me1 modifications (associated with active enhancers) across 127 cell types, enrichment of specific cell types was investigated (**Methods**). Cells that surpassed the false discovery threshold rate (FDR < 0.05) were mostly hematological cells, with B and T lymphocytes the most prominent cell types associated with SLE (**Extended Data Fig. 6a**). Similar results were observed based on H3K4me3 modifications (associated with promoters of active genes) (**Extended Data Fig. 6b**).

### Identification of putative disease genes and pathways

Excluding the human leukocyte antigen (HLA) region, 179 putative disease genes were identified across the disease-associated loci reported before and those newly identified in this study (**Extended Data Table 5**; DEPICT^25^, **Methods**). A significant level of protein-protein connectivity corresponding to genes found at the novel loci and known SLE-associated loci was observed (*P* < 1E-16; **Extended Data Fig. 7**). Forty-five pathways were significantly enriched with these putative SLE susceptibility genes (ToppGene^26^, FDR < 0.05; **Extended Data Table 6**). The pathways of cytokine signaling, IFN-α/β signaling, Toll-like receptor (TLR) signaling, and B and T cell receptor signaling showed greatest enrichment. The RIG-I-like receptor signaling (*P* = 5.83E-10) and TRAF6-mediated IRF7 activation (*P* = 6.41E-10) pathways were designated as SLE associated pathways primarily based on genes newly identified in this study.

### Trans-ancestral fine-mapping of disease-associated loci

One hundred and eight SLE-associated loci tagged by SNPs having a minor allele frequency (MAF) greater than 1% in both Chinese and European populations were examined by PAINTOR^27^, making use of the differences in LD between ancestries. The median number of putative causal variants in the 95% credible sets reduced from 57 per locus when using only the European GWAS to 16 per locus when using data from both ancestries (one-sided paired t-test *P* = 9.79E-07, **Fig. 2**). The number of disease-associated loci with five or fewer putative causal variants increased from four when using the European GWAS alone to 15 (**Extended Data Table 7**). A single putative causal variant was identified for the *WDFY4* and *TNFSF4* loci, the latter of which was functionally validated in a previous study^28^ (**Extended Data Fig. 8**).

**Fig. 1.**
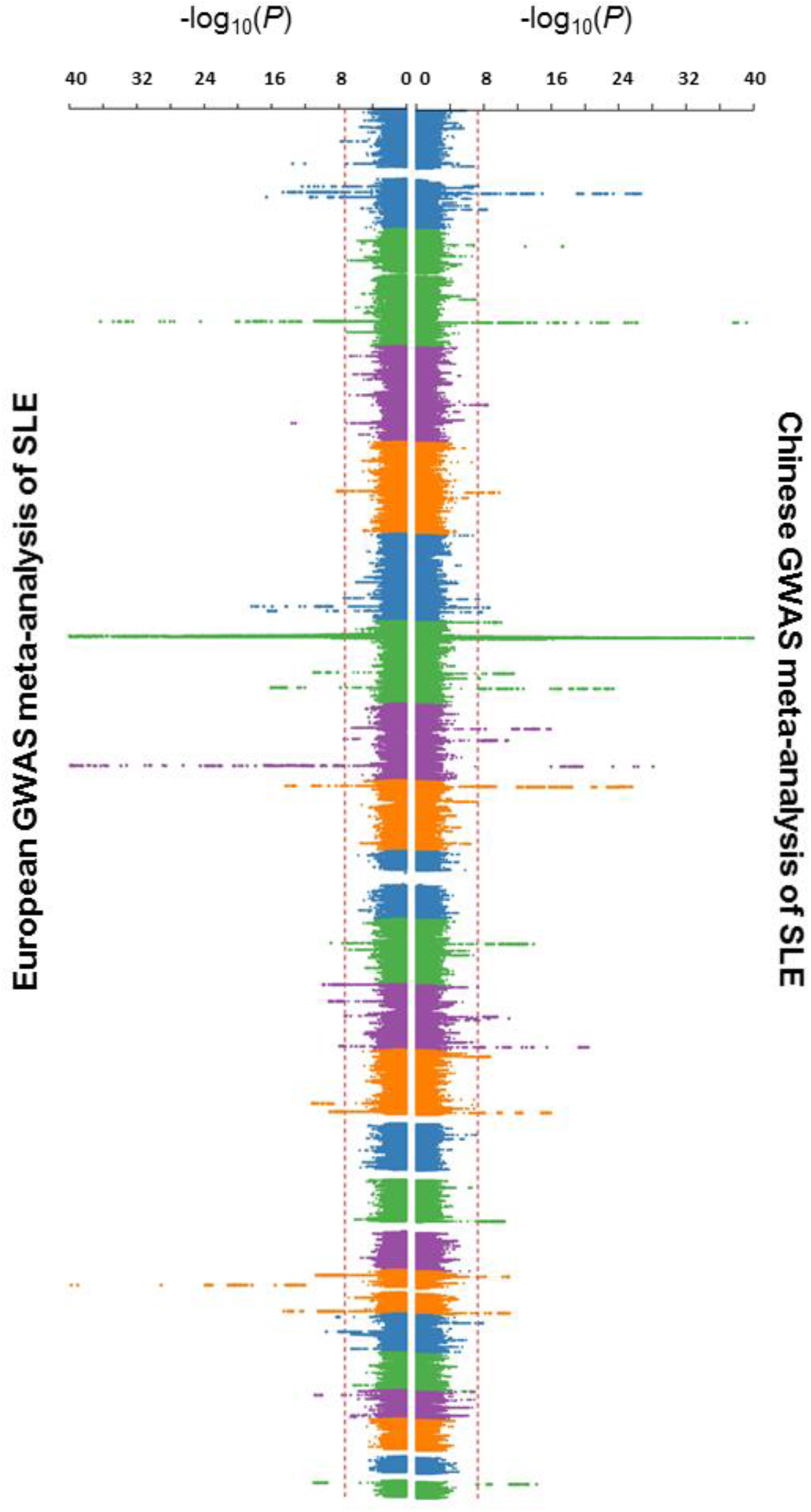
Manhattan plots for the Chinese and European SLE GWAS meta-analyses results. The Chinese SLE GWAS comprised 4,222 cases and 8,431 controls and the European GWAS comprised 4,576 cases and 8,039 controls. The X-axis is the *P*-value of association, as −log_10_ (*P*), for the meta-analyses of the Chinese (**right**) and European (**left**) ancestries. Red dashed lines indicate the threshold of genome-wide statistical significance (*P* = 5E-08). SNPs with *P* < 1E-40 in an associated locus are not shown from the plot.

**Fig. 2.**
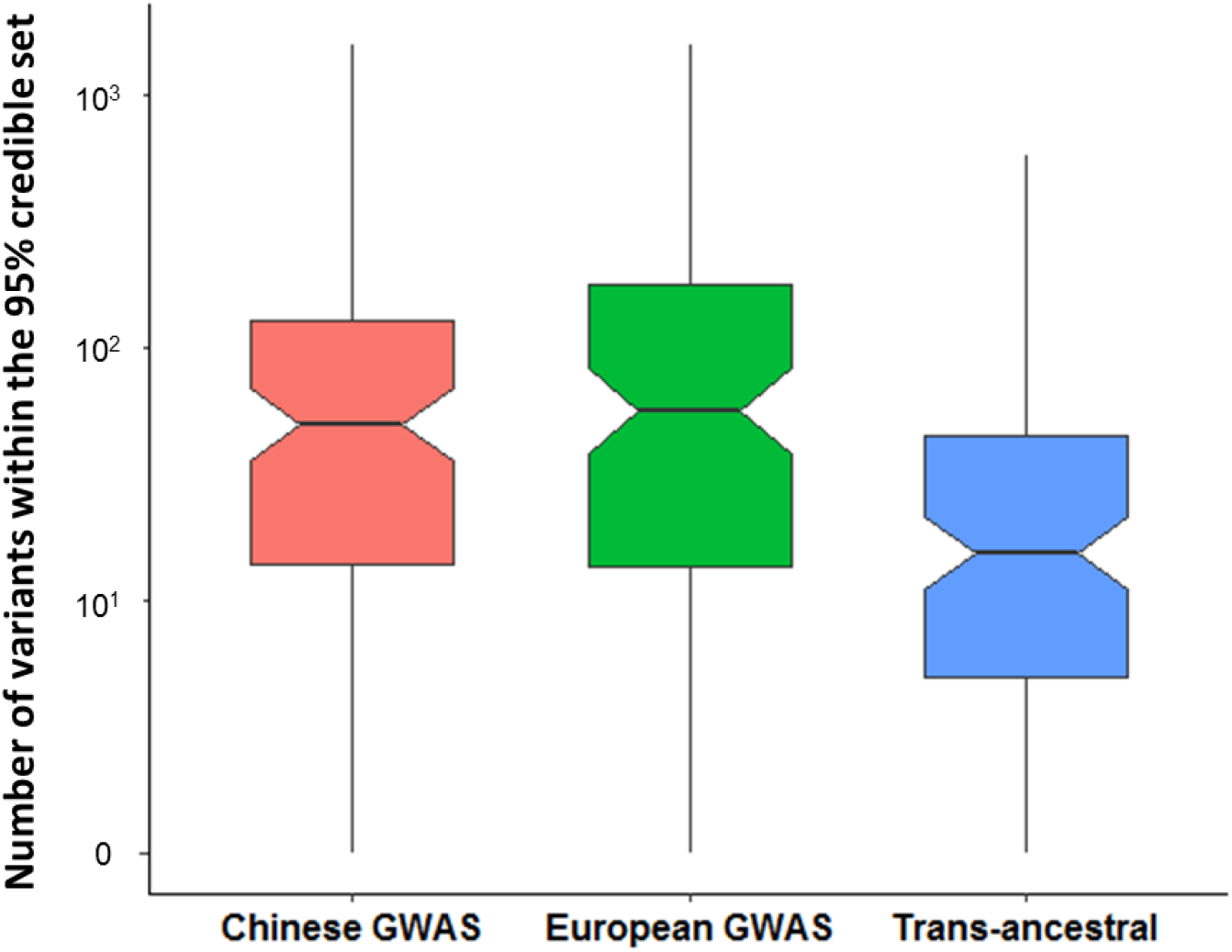
Fine-mapping across SLE-associated loci based on the association results from the Chinese SLE GWAS, the European GWAS, and the trans-ancestral meta-analyses. The Y-axis indicates the number of potential causal variants at each locus based on the 95% credible sets of the association results from the Chinese SLE GWAS (**red**), the European GWAS (**green**) and trans-ancestral meta-analyses (**blue**).

### Ancestral group differences

Based on the analysis of Cochran’s Q (CQ)-test that assesses heterogeneity of effect-size estimates from different ancestral groups, SLE-associated variants or loci were divided into four categories: 1) ancestry-shared disease loci tagged by variants with CQ-test *P* ≥ 0.05; 2) putative ancestry-heterogeneous disease loci with CQ-tests of *P* < 0.05 but FDR adjusted CQ-test *P* ≥ 0.05; 3) ancestry-heterogeneous disease loci with FDR adjusted CQ-test *P* < 0.05; and 4) disease loci tagged by associated variants with the risk allele absent in one of the two ancestries^15^ (**Extended Data Table 10**). Nine disease variants, other than those absent or rare (MAF < 0.01) in one of the two ancestries^15^, showed significant differences in effect-size estimates between the two ancestral groups and were considered ancestry-heterogeneous (FDR adjusted CQ-test *P* < 0.05, category 3; **Fig. 3a** and **Extended Data Table 8**). Within this category, variants in the *HIP1*, *TNFRSF13B*, *PRKCB*, *PRRX1*, *DSE* and *PLD4* loci were associated with SLE only in East Asians and variants in *TYK2* and *NEURL4-ACAP1* only in Europeans (*P* < 5.0E-08 in one ancestry but *P* > 0.01 in the other, with non-overlapping of the 95% CIs of the ORs). These eight loci were thus considered ancestry-specific. SNP rs4917014, a variant near *IKZF1*, showed a significantly stronger effect in East Asians (OR = 1.33, *P* = 5.18E-29) than in Europeans (OR = 1.16, *P* = 1.34E-06; CQ-test *P* = 4.02E-04). These findings were supported by analyses in each cohort (**Extended Data Fig. 9-10**).

**Fig. 3.**
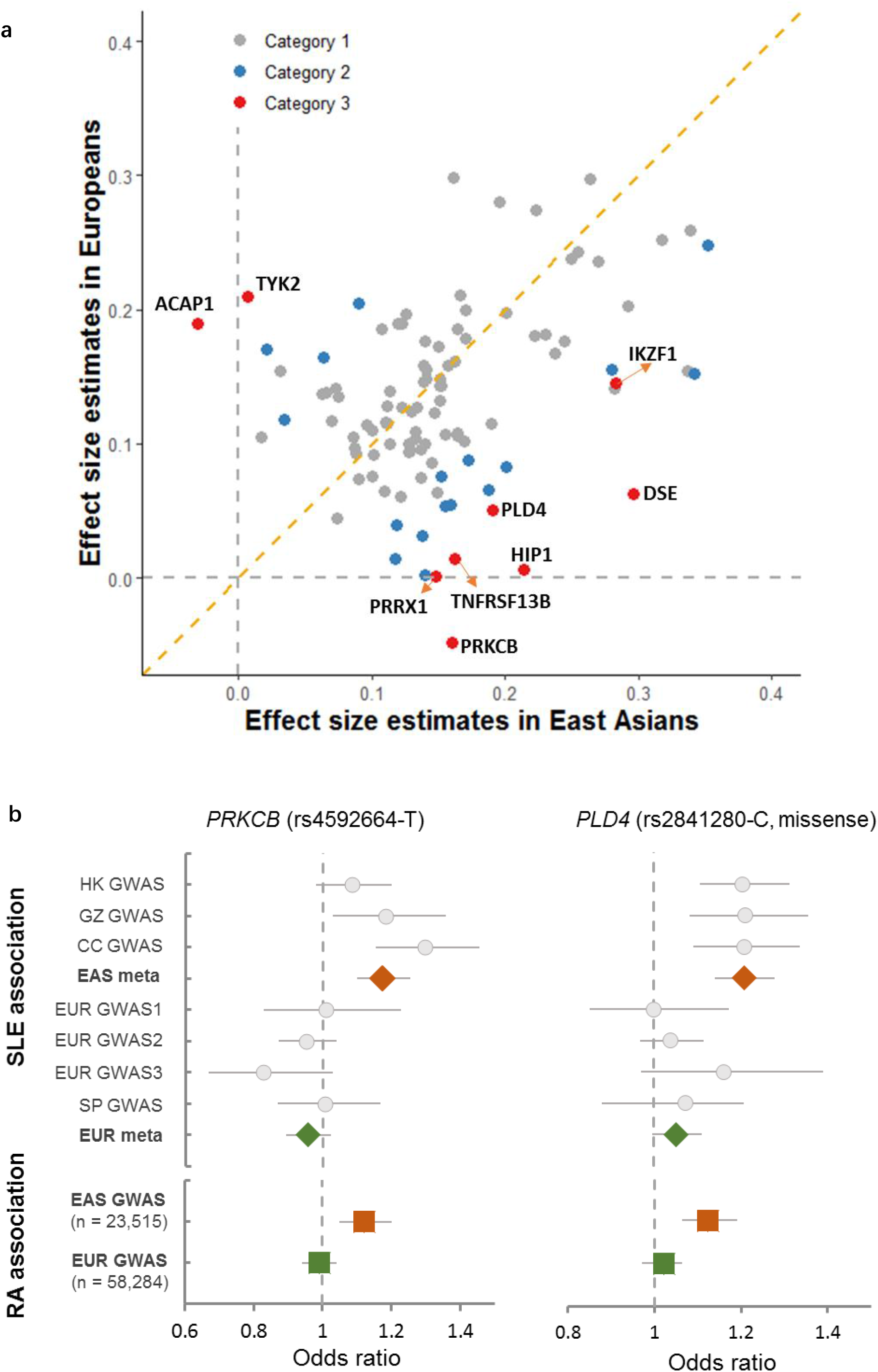
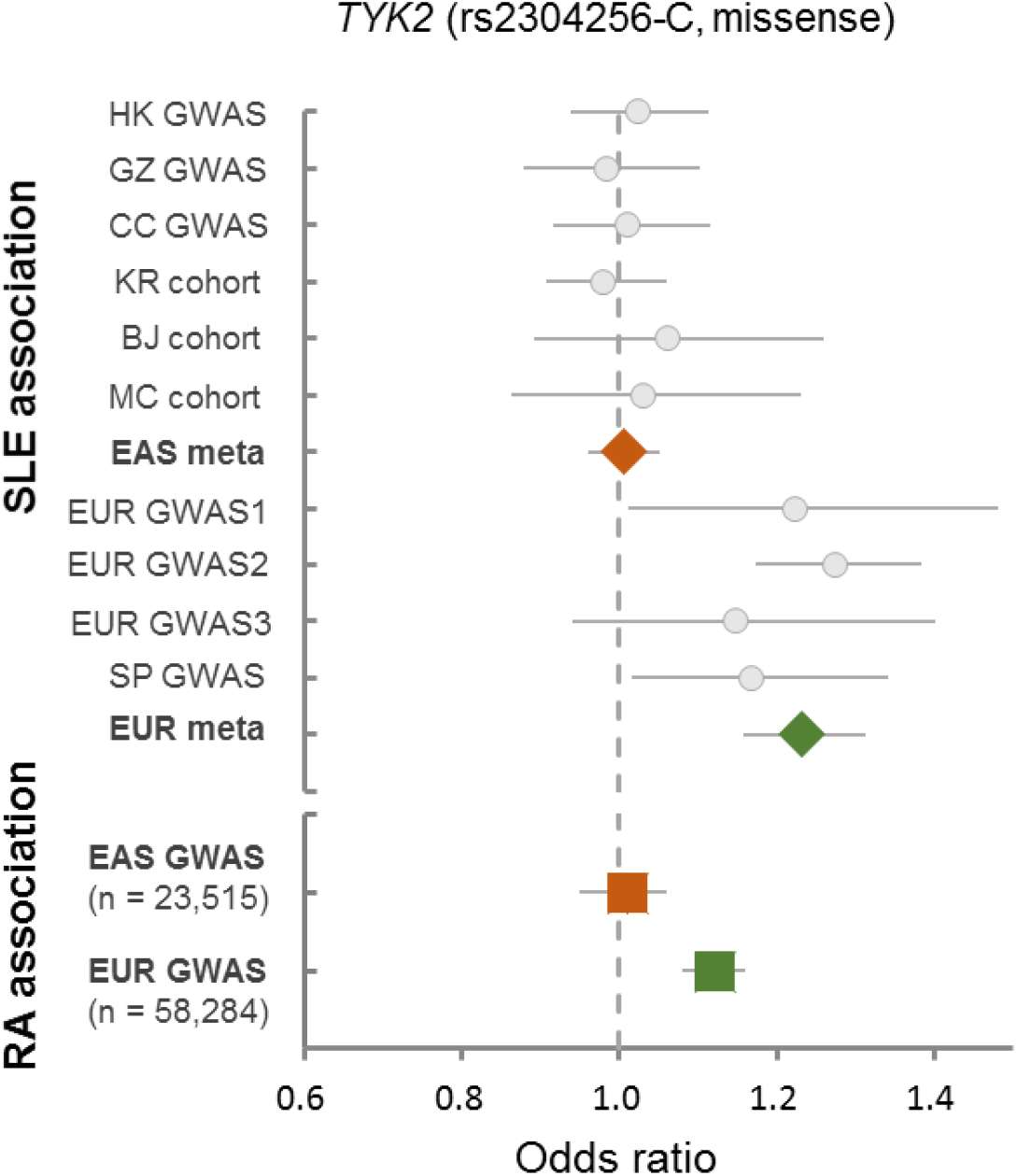
Genetic loci showing significant ancestral differences in effect-size estimates for SLE and RA. **a.** Correlation of effect-size estimates for SLE between East Asian (X-axis) and European (Y-axis) populations. Disease-associated variants with FDR adjusted CQ-test *P*-value < 0.05 (category 3) are labeled in **red**, and the variants with CQ-test *P*-value < 0.05 but FDR adjusted CQ-test *P*-value ≥ 0.05 (category 2) are labeled in **blue**. **b.** Forest plots of association from each cohort at the *PRKCB*, *PLD4* and *TYK2* loci. Diamonds represent the combined estimates of odds ratio (OR) in East Asians (EAS, **red**) and Europeans (EUR, **green**) for association with SLE. Error bars represent the 95% confidence intervals of the OR estimates. Squares represent OR estimates for rheumatoid arthritis (RA) in EAS (**red**) and EUR (**green**) populations. Regional plots for each locus are available in **Extended Data Fig. 10.** HK, Hong Kong; GZ, Guangzhou; CC, Central China; KR, Korean; BJ, Beijing; MC, Malaysian Chinese; SP, Spanish

On reanalyzing data from association studies on rheumatoid arthritis (RA)^29^, the variants in the *PRKCB* and *PLD4* loci were found to be associated with RA in East Asians but not in Europeans (CQ-test *P* = 0.004 and 0.010 for *PRKCB* and *PLD4*, respectively), while the variant in *TYK2* was found associated in Europeans only (CQ-test *P* = 0.002; **Fig. 3b**). This consistency with the differences found in the SLE study suggests that shared mechanisms could be responsible for ancestral group differences among autoimmune diseases.

### Colocalization of loci across ancestral groups

Colocalization methods consider many SNPs, rather than only the leading variant in a locus, to compare association signals between ancestral groups. The ancestry-shared disease loci showed much higher posterior probabilities (PP) of colocalization (mean of PP = 0.42) than the ancestry-specific loci (mean of PP = 0.03; **Extended Data Fig 11a**). For example, the *MTF1*, *IKBKE* and *TNIP1* loci in category 1 showed strong posterior probabilities (≥ 90%) of colocalization under a Bayesian test^30^(**Methods**), suggesting shared causal effects between the two ancestries. The eight ancestry-specific loci in category 3 showed low posterior probabilities of colocalization (1% – 10%), consistent with the CQ-test of the top variant of each locus (above and **Extended Data Fig. 10**).

Since LD differences between ancestries may affect the colocalization results, we compared the SLE association signals from the Chinese populations with those on 27 non-immune-related phenotypes studied in Europeans (**Extended Data Table 9**) to serve as colocalization baseline values. While posterior probabilities for colocalization at the ancestry-shared *MTF1*, *IKBKE* and *TNIP1* loci were much greater than the baseline values, there were no differences for the six Asian-specific disease loci (**Extended Data Fig. 11b**), thereby excluding the potential influence of LD. The European-specific loci (*TYK2* and *NEURL4-ACAP1*) were not evaluated this way due to lack of public data.

### Functional annotation of the ancestry-heterogeneous loci

The ancestry-heterogeneous loci appear to be enriched for functions related to antibody production. Two of the nine putative disease genes at ancestry-heterogeneous loci (category 3), *TNFRSF13B* and *IKZF1*, are causal genes for human primary immunodeficiency disorders (PID)^31^ presenting with primary antibody deficiencies (PADs), whereas none of the disease genes at the putative ancestry-heterogeneous loci (category 2, 0/22; Fisher exact test *P* = 0.07) or the ancestry-shared loci (category 1, 0/120; Fisher exact test *P* = 0.004) are known to cause PADs in humans. Two of the East Asian-specific disease variants, those in the *TNFRSF13B* and *PRKCB* loci, were associated with serum immunoglobulin levels in East Asian populations^32–34^, whereas none of the variants in loci belonging to category 1 and 2 were found to be associated.

*TNFRSF13B*, which encodes a BAFF receptor, TACI, plays a major role for immunoglobulin production^35–37^. In this study, a missense variant in *TNFRSF13B*, rs34562254, was specifically associated with SLE in Chinese populations (OR = 1.18, *P* = 2.88E-08 in Chinese; OR = 1.01, *P* = 0.75 in Europeans). In European populations, an SLE-associated variant in the 3’-UTR of *TNFSF13B* (encoding BAFF), which is absent in Asian populations (category 4), was associated with serum levels of total IgG, IgG1, IgA and IgM^38^.

Mice deficient in the orthologs of four of the nine putative disease genes (44.4%) at the ancestry-heterogeneous loci for SLE, *Tnfrsf13b-*, *Ikzf1-*, *Prkcb-* and *Tyk2*-, demonstrated abnormal IgG levels^39^ (MP:0020174), while at putative ancestry-heterogeneous loci (4/22 or 18.2%; OR = 3.43, Fisher exact test *P* = 0.185) or ancestry-shared loci (14/120 or 11.6%; OR = 5.92, Fisher exact test *P* = 0.027), proportionately fewer genes caused aberrant IgG levels in mice. Orthologs of four of the twelve (33.3%) putative disease genes from disease loci where the risk allele is monomorphic in one of the ancestries (*PTPN22, TNFSF13B, IKZF3* and *IGHG1*; category 4) also demonstrated abnormal immunoglobulin production in gene knockout mouse models^35,38,40–42^.

### Evolutionary signatures for the disease loci

Disease loci with heterogeneity between East Asians and Europeans might have undergone differential selection pressures in recent human history, as has been shown for the SLE risk variant in *TNFSF13B*^38^. Frequency variances, as fixation indexes (F_st_), for the variants of the first three categories were calculated using 3,324 controls from the HK cohort and 5,379 controls from EUR GWAS 2 cohort (**Methods**). Higher F_st_ would indicate a larger frequency difference between the two ancestries. Mean F_st_ values for the ancestry-shared, putative ancestry-heterogeneous and ancestry-heterogeneous variants were 0.054, 0.061 and 0.084, respectively. Although a small sample, three (*DSE*, *HIP1*, *TNFRSF13B*) of the nine ancestry-heterogeneous variants (33.3%) showed F_st_ ≥ 0.15 (empirical *P* < 0.03), while only 10% of the putative ancestry-heterogeneous variants and 8.8% for the ancestry-shared disease variants had F_st_ ≥ 0.15 (**Extended Data Fig. 12a**).

A significant correlation of standardized integrated Haplotype Scores (iHS)^43^, estimated using control subjects from HK and EUR GWAS 2 cohorts (**Methods**), gave evidence of shared recent positive selection for the ancestry-shared disease variants (categories 1; r = 0.28, *P* = 0.03; **Extended Data Fig. 12b**). This is consistent with results using data from Southern Han Chinese (CHS) and Utah residents of European ancestry^44^ (CEU; r = 0.32, *P* = 0.008). However, there was no evidence of such a correlation for disease variants that showed ancestry heterogeneity (category 2 and 3; **Extended Data Fig. 12b**). For example, in the BAFF system, the derived risk allele rs34562254-A in *TNFRSF13B* is much more prevalent in East Asians than in other populations (**Figure 4a**) and has a significantly longer haplotype for the derived risk allele than the ancestral allele (more negative standardized iHS score) in East Asian populations than in Africans (*P* = 3.2E-04) or Europeans (*P* = 4.4E-04) (**Figure 4b**), suggesting recent positive selection for the risk allele in East Asians.

**Fig. 4.**
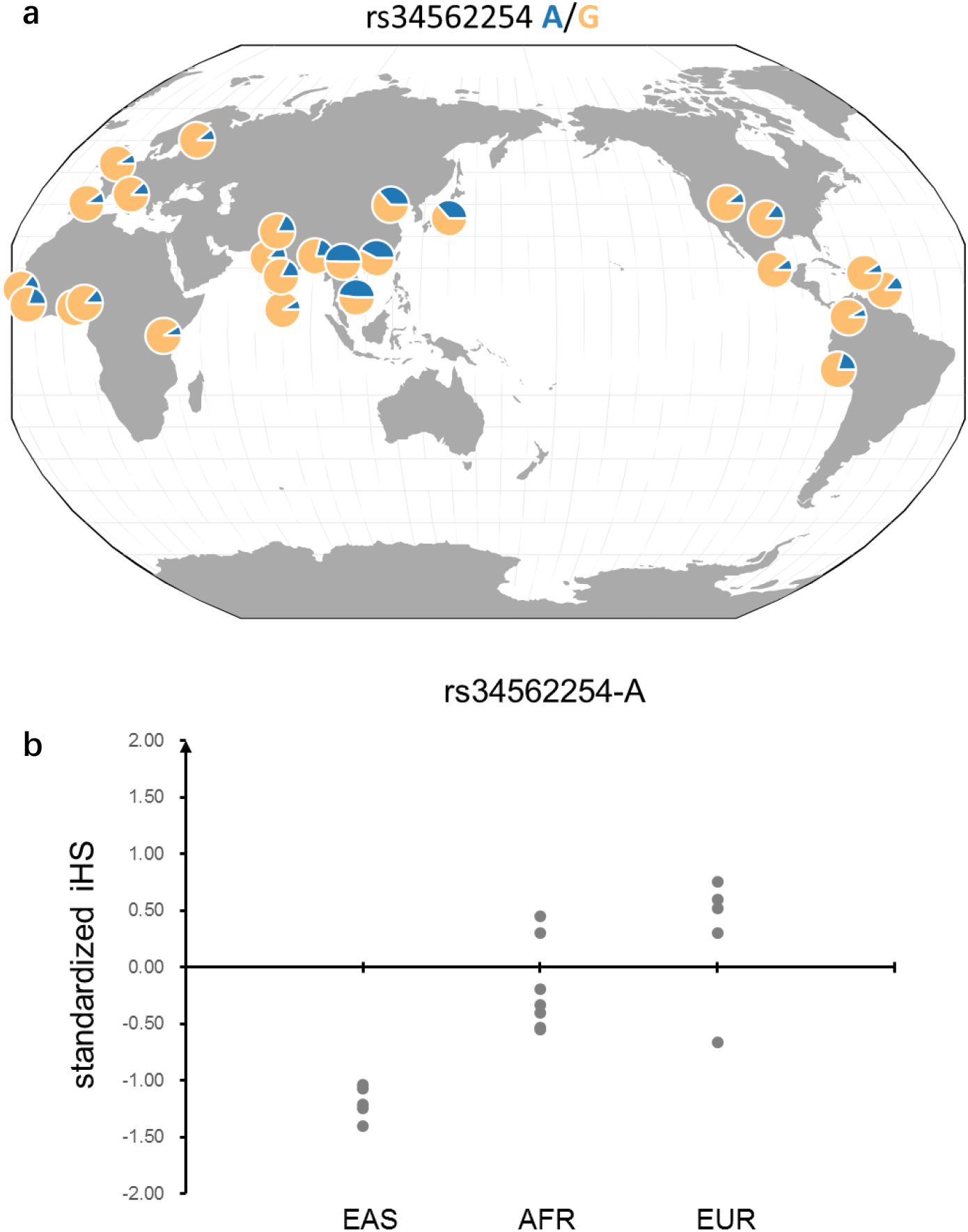
Risk allele frequency and standardized iHS scores across different populations for the Asian-specific SLE variant at the *TNFRSF13B* locus. **a**. Frequency for the risk allele rs34562254-A across populations. **b**. Standardized iHS for the variants across different continental populations (EAS: East Asians; AFR: Africans; EUR: Europeans). The risk allele rs34562254-A is a derived allele, and negative iHS value indicates that the haplotypes carrying the derived allele are longer than the haplotypes carrying the ancestral allele. The frequency and iHS were calculated using data from the 1,000 Genomes Project.

### Polygenetic risk scores for SLE and their accuracies across ancestries

Polygenic risk scores (PRS) have been used to estimate individual risk to complex diseases, such as coronary artery disease^45^ and schizophrenia^46^. However, as the majority of GWAS findings used to calculate these scores are based on European populations, their accuracy in other populations may be limited. PRS for SLE, trained by data on European populations, were tested on individuals of three Chinese cohorts using the lassosum algorithm^47^ (**Methods**). The area under the receiver-operator curve (AUC) ranged from 0.62 to 0.64 for the three Chinese cohorts. Similar results were observed in the reverse case (**Extended Data Fig. 13a**). The LDpred^48^ algorithm produced similar results (**Extended Data Fig. 13b**). These analyses suggest a partial transferability of PRS between the two ancestries.

Using samples from the GZ cohort as the validation dataset, performance of predictors trained using GWAS summary statistics from the HK and CC cohorts (2,618 cases and 7,446 controls) or from the European cohorts (4,576 cases and 8,039 controls) were evaluated. Ancestry-matched predictors significantly outperformed (AUC = 0.76, 95% CI: 0.74-0.78) ancestry mismatched predictors (AUC = 0.62, 95%CI: 0.60-0.64) (**Fig 5a)**. When the analysis was repeated by randomly choosing the same number of samples (1,500 cases and 1,500 controls) from each of the Chinese and European GWAS as training data, a similar difference was observed (**Extended Data Fig. 14**). Ancestry-matched PRS for samples in the GZ cohort had a mean difference of 0.89 (standard deviation) between the SLE case and control groups (*t*-test *P* = 9.01E-116) and disease classification using the optimal threshold achieved 73.4% sensitivity and 65.4% specificity (**Fig 5b**; **Methods**). Disease risk increased with higher PRS, with individuals in the highest PRS decile having a much higher disease risk than those in the lowest decile (OR = 30.3, Chi-square test *P* = 6.23E-54; **Fig 5c**).

**Fig. 5.**
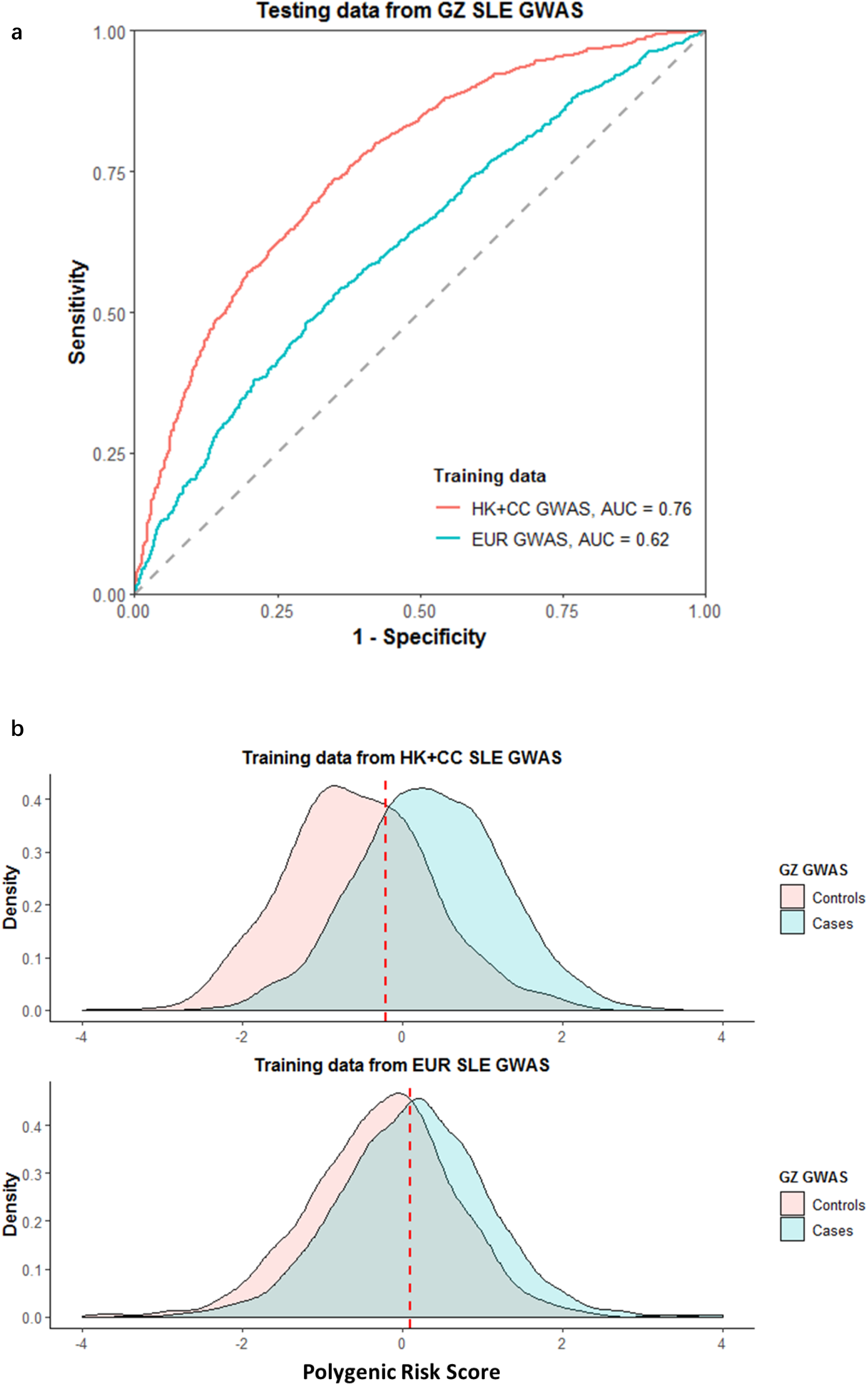
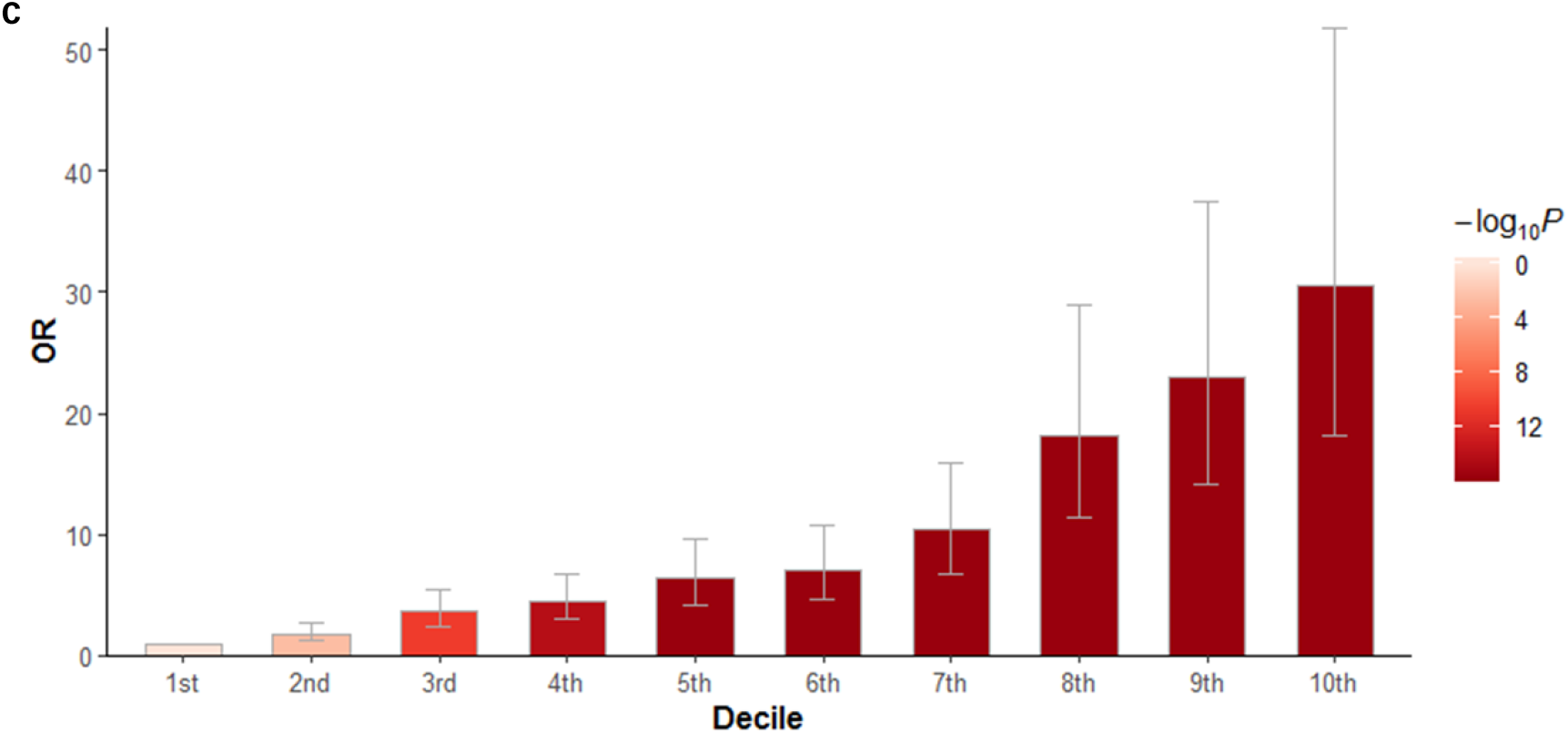
Performance of polygenic risk scores (PRS) calculated by summary statistics from different ancestral groups. **a**. Performance of PRS evaluated by area under the ROC curve (AUC). PRS for individuals from the GZ cohort were calculated using summary statistics from ancestry-matched Chinese populations (HK and CC GWAS, 2,618 cases and 7,446 controls; **red**) and European populations (4,576 cases and 8,039 controls; **blue**). **b**. Distribution of PRS for SLE cases (**blue**) and controls (**pink**) from the GZ cohort. The PRS distribution was estimated using summary statistics from the ancestry-matched Chinese cohorts (upper panel) or from the mismatched European GWAS (lower panel). The optimal PRS threshold for disease risk prediction is indicated by the red vertical line. **c.** Odds ratios (ORs) of disease risk for individuals in the GZ GWAS across different deciles of PRS calculated by ancestry-matched data. ORs and 95% confidence intervals (bars) were calculated by reference to the 1^st^ PRS decile.

## Discussion

Despite evidence of ancestral group differences in the disease manifestations of SLE, genetic studies to date have been unable to examine these differences in detail as most earlier data focused on European ancestral groups. As non-European groups appear to be more severely affected by SLE, greater genetic data on these groups are likely to be highly informative. By increasing the number of subjects of East-Asian ancestry to levels roughly equivalent to those of European subjects, and including previously published data, we have made cross ancestral group studies possible.

Through systematic comparisons between the two ancestral groups, it was found that there was a significant, but not complete, sharing of the genetic architecture of SLE between Chinese and Europeans. SLE showed a r_ge_ of 0.64. This represents a much lesser level of genetic sharing than seen for schizophrenia^49^ between East Asian and European ancestries (r_ge_ = 0.98).

Through ancestry-dependent and trans-ancestral meta-analyses, we identified 38 novel loci associated with SLE, bringing the total number of SLE-associated loci to 132. High level functional annotation of these SLE associated loci implicated hematological cells, particularly B and T cells, cytokine signaling and other immune system pathways. Consistent with previous findings^50^, we demonstrated the value of trans-ancestral data in significantly reducing the number of putative causal variants at each disease-associated locus, which may facilitate future functional and mechanistic studies.

There was strong evidence of heterogeneity on SLE associations between the two ancestries, which were not likely to be artefacts of study power. Eight variants (that are common in both ancestral groups) were associated with disease in only one of the ancestral groups. For three of these, a similar ancestral difference was found by re-analyzing association data on RA^29^. This might suggest that common mechanisms account for ancestral differences in autoimmune diseases.

Genes at the ancestry-heterogeneous disease loci seemed more likely to be involved in regulation of immunoglobulins than genes at the ancestry-shared disease loci. Immunoglobulin levels are highly heritable^51^ and have been found to differ between ancestries, with African Americans and Asians having higher serum immunoglobulin levels than people of European ancestry^52–55^. Higher antibody levels in non-European populations might have contributed to their higher prevalence of SLE and further study of intrinsic differences in immune function among ancestries may be informative.

That differential mechanisms may exist for antibody regulation between East Asian and European populations is supported by the association of SLE with the BAFF signaling system. This system, which may be part of the initial reaction to host-pathogen interactions^36^, is under positive selection based on different environmental exposures^36,38^. BAFF and its receptors, including TACI, play essential roles in B cell survival and differentiation^35–37^. The SLE risk allele in the gene encoding BAFF is completely absent in Chinese populations^38^ and a missense variant in the gene encoding TACI was found to be specifically associated with SLE in East Asians in this study. Both these genes, *TNFSF13B*^38^ in Europeans and *TNFRSF13B* in East Asians, were found to have undergone positive selection in recent human history. Adaptation of the host to resist pathogens may underlie some of the ancestral group-heterogeneity.

TACI is expressed at very low levels in human newborns and mice before exposure to pathogens^56,57^ and previous studies have shown that certain pathogens can ablate B cell responses by modulating the expression of TACI^58–60^. The BAFF risk allele was shown to significantly up-regulate humoral immunity^38^, and whether this is the case for the risk allele in TACI should be investigated. TACI blockers, such as Atacicept^61^, might give better responses to SLE in patients of Asian ancestry and the variant found in *TNFRSF13B* may be a useful genetic marker for the prescription of Belimumab, a hypothesis that may warrant further study.

Our analyses have identified a substantial number of novel SLE-associated genetic loci and deepened our understanding of the genetic factors that may underly the differences in the manifestation of SLE between peoples of European and non-European ancestry. Like a recent PRS study in SLE^62^, but to a greater extent, we have shown that PRS achieved a far better performance when based on ancestry-matched populations. Our findings contribute new insights into precise treatments, and to risk prediction and prevention of SLE.

## Methods

### Overview of samples

8,252 subjects of Han Chinese descent from Hong Kong (HK), Guangzhou (GZ) and Central China (CC) were genotyped in this study. The institutional review boards of the institutes collecting the samples (The University of Hong Kong, Hospital Authority Hong Kong West Cluster and Guangzhou Women and Children’s Medical Center) approved the study and all subjects gave informed consent. These subjects were genotyped by the Infinium OmniZhongHua-8, the Infinium Global Screening Array-24 v2.0 (GSA) and the Infinium Asian Screening Array-24 v1.0 (ASA) platforms. Fourteen samples were randomly selected and genotyped by different platforms. High concordance rates (> 99.9%) were observed for genotypes derived from the different platforms (**Extended Data Table 1**). Principal component (PC) analysis was performed to examine potential batch effects and no significant differences were observed from the PCs for data genotyped by different platforms and from different batches **(Extended Data Fig. 1)**. These new data were analyzed with data from our existing SLE GWAS^13^, which contained 5,057 Chinese subjects.

### Quality control and association study

Genotype Harmonizer^63^ was used to align the strands of variants of the Chinese GWAS to the reference of the 1,000 Genomes Project Phase 3 panel. Variants with a low call rate (< 90%), low minor allele frequency (< 0.5%) and violation of Hardy-Weinberg equilibrium (*P*-value < 1E-04) were removed. Quality control required the following criteria: i) missing genotypes were below 5%, ii) hidden relatedness (identity-by-descent) with other samples was ≤ 12.5% iii) inbreeding coefficients with other samples ranged from −0.05 to 0.05 and iv) not having extreme PC values as computed for individuals using EIGENSTRAT embedded in PLINK^64,65^. After quality control, pre-phasing used SHAPEIT^66^ and individual-level genotype data were imputed to the density of the 1,000 Genomes Project Phase 3 reference using IMPUTE2^67^. For association analysis SNPTEST^68^ was used to fit an additive model. Top PCs and the BeadChip types were included as covariates. The number of PCs to be adjusted for in each analysis was determined using a scree plot with a cutoff when the plot levels off. Variants with imputed INFO scores < 0.7 were excluded. The genomic inflation factors (λ_GC_) for the HK, GZ and CC GWAS were 1.04, 1.03 and 1.04, respectively, and the LD score regression (LDSC)^69^ intercepts were 1.03, 1.02 and 1.03, respectively. Manhattan plots for each cohort are shown in **Extended Data Fig. 3**.

For the European SLE GWAS data, the λ_GC_ and LDSC intercept listed in LD hub seemed inflated (λ_GC_ = 1.17 and LDSC intercept = 1.10)^20^. Thus, the data were reanalyzed to minimize the potential influence of sub-population stratification (and see below). PCA analysis showed that subjects from the existing European data were more diverse than the Chinese subjects used in this study (**Extended Data Fig. 2a**). The European individuals were grouped into three cohorts by their PCs relative to the subjects of the 1,000 Genomes Project. Subjects in the EUR GWAS 1, EUR GWAS 2, EUR GWAS 3 cohorts shared similar PCs with individuals of Spanish (IBS), northern and western European (CEU and GBR) and Italian (ITS) origins, respectively (**Extended Data Fig. 2b** and **Extended Data Table 2**). Quality control, imputation and association analyses were conducted, as for the Chinese datasets, in each cohort. λ_GC_ for the three European GWAS datasets were 1.05, 1.08 and 1.03, respectively, and the LDSC intercepts were 1.03, 1.04 and 1.00, respectively. Manhattan plots for each cohort are shown in **Extended Data Fig. 4**.

### Genetic correlation between the two ancestries

Trans-ancestral genetic correlation from the meta-analysis results for Chinese and Europeans were estimated using the Popcorn algorithm^23^ based on common SNPs in the autosomes. The disease prevalence in Chinese and European populations were set to be 1‰ and 0.3‰, respectively^4^. SNPs were removed from this analysis according the following criteria: 1) SNPs with strand-ambiguities (A/T or C/G alleles); 2) having MAF < 5%; 3) having imputed INFO score < 0.9. The cross-population LD scores were estimated using control subjects from the HK cohort (n = 3,324) and EUR GWAS 2 cohort (n = 5,379).

### Meta-analyses of SLE association studies

Meta-analyses for the Chinese and European SLE GWAS were conducted independently. The summary association statistics from HK, GZ and CC GWAS (4,222 SLE cases and 8,431 controls) were combined in a meta-analysis using a fixed-effect model, weighted by the inverse-variance^70^. The λ_GC_ for the Chinese meta-analysis was 1.09 and the LDSC intercept was 1.04. For the European data, the EUR GWAS 1-3 datasets were combined with a new GWAS on a Spanish population^21^ in the meta-analysis (4,576 cases and 8,039 controls). λ_GC_ and the LDSC intercept for the European SLE meta-analysis reduced to 1.11 and 1.03, respectively.

Trans-ancestral meta-analysis across the Chinese and European GWAS cohorts used the fixed-effect model. To increase statistical power, summary association statistics from an Immunochip study were included as an *in silico* replication, adding 2,485 SLE cases and 7,616 controls from Korea (KR), Han Chinese in Beijing (BJ) and Malaysian Chinese (MC)^22^. 11,283 SLE cases and 24,086 controls were involved in this study (**Extended Data Table 2**).

### Heritability explained by the SLE-associated variants

The variance in liability explained by the SLE-associated variants was measured using VarExplained program^71^. Variants in the HLA region were excluded in the analysis. The disease prevalence was set to be 1‰ for East Asians and 0.3‰ for Europeans^4^. The novel loci increased the heritability explained from 0.10 to 0.13 for East Asians, and from 0.08 to 0.09 for Europeans.

### Functional annotations of SLE associated SNPs

The stratified LD score regression method^24^ was applied on the trans-ancestral meta-analysis result to partition SNP-heritability across functional annotations. Twenty-eight categories of annotations that are not cell type specific (**Extended Data Fig. 5**) provided by this source were studied. For cell type-specific analyses, H3K4me1 and H3K4me3 modifications across 127 cell types **(Extended Data Fig. 6**) were downloaded from the Roadmap Epigenomics Project^72^. The cell type-specific enrichment was performed under the “full baseline” model^24^, which aimed to control for overlaps with annotations that are not cell type-specific.

### Identification of putative SLE genes and gene-set enrichment analysis

Putative causal gene(s) across all the SLE-associated loci outside of the HLA region were identified using DEPICT^25^. The default setting (r^2^ > 0.3) was used to set boundaries for each SLE associated locus. Genes within (or overlapping) the boundaries were examined and those with a *P*-value < 0.05 were defined as putative causal genes. If no genes were selected at that locus, gene(s) identified from eQTL data from human whole blood^73,74^ were considered to be putatively causal. Gene-set enrichment analysis was performed using ToppGene^26^, with the August, 2019 versions of the KEGG^75^, Reactome^76^ and mouse knockout phenotype^39^ databases. The 2017 IUIS Phenotypic Classification for Primary Immunodeficiencies^31^ (PID) was used to obtain 320 human PID genes categorized into nine phenotypic classifications.

### Trans-ancestral fine-mapping of the associated loci

The HLA region was excluded from this analysis, as extensive LD and limited genotyping of SNPs in both ancestries makes defining the best model of association difficult for this region. Disease loci with rare risk alleles (MAF < 0.01) or absent in one ancestry were also excluded, leaving 108 SLE-associated loci in the autosomes for this study. For each disease locus, all variants within the region were extracted for both ancestries. The genetic interval was determined by the closest recombination hotspots around a given disease-associated variant (defined as a recombination rate < 10 cM/Mb). A fine-mapping algorithm, PAINTOR (version 3.0)^27^, was used to estimate the posterior probability of causality for each variant at a given locus based on the trans-ancestral model. For comparison, we also applied the fine-mapping algorithm on the Chinese and European SLE GWAS, separately. All analyses were run under the assumption of a single causal variant per locus. The LD matrix was calculated using control samples from HK (n = 3,324) and EUR GWAS 2 (n = 5,379) for Chinese and European populations. Variants with a cumulative posterior probability greater than 95% were defined as putative causal variants (95% credible set).

### Identification of loci with differential effects between the two ancestries

Cochran’s Q (CQ)-test^77^ was used to examine effect-size differences between the two ancestries for all the disease-associated variants in the autosomes. If the variants were also interrogated by the Immunochip^22^ system, the association results derived from the KR, BJ and MC cohorts were also included. CQ-test *P*-values were adjusted for a cutoff of 0.05 using the Benjamini-Hochberg method^78^. For comparison, summary association statistics on RA were downloaded from a previous study^29^ of 4,873 RA cases and 17,642 controls of Asian ancestry and 14,361 RA cases and 43,923 controls of European ancestry.

### Colocalization analysis

Colocalization of association signals from the two ethnicities was determined using the R package coloc^30^ on all variants with a MAF > 1% and imputed (IMPUTE2^67^) INFO score > 0.9 within a given disease locus. To control for LD differences between ancestries, SLE association signals from Chinese populations were compared with those from 27 immune-unrelated phenotypes from European populations (LD hub^20^; **Extended Data Table 9**) to generate baseline posterior probabilities of colocalization in the absence of a phenotypic relationship. Ancestry-shared causal effects for SLE were expected to be significantly greater than the baseline values.

### Analysis of selection signatures for the associated variants

The fixation index (F_st_) was used to test allele-frequency differences between the two ancestries. F_st_ was calculated based on the following formula^79^:

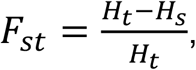

where *H*_*t*_ is the expected proportion of heterozygosity in the pooled samples from all ethnicities based on Hardy-Weinberg equilibrium: 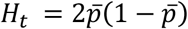,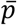 is the allele frequency in the overall pool. *H*_*s*_, the expected proportion of heterozygosity in a subpopulation (either Chinese or Europeans), is estimated as 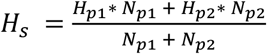 where *H*_*pi*_ is the expected heterozygosity in the i^th^ subpopulation estimated by the allele frequency in that subpopulation under Hardy-Weinberg equilibrium. *N*_*pi*_ is the sample size of the i^th^ subpopulation.

Potential selective sweeps in respective ancestries were examined using the Integrated Haplotype Score (iHS) method, which measures the extended haplotype homozygosity for the ancestral allele relative to the derived allele^80^. Raw iHS scores were computed using the R package rehh^81^ and normalized by different frequency bins (50 bins over the range 0 to 1). Large negative standardized iHS values indicate long haplotypes carrying the derived allele, while large positive values suggest long haplotypes with the ancestral allele. The F_st_ and iHS values analyzed in this study were estimated using control subjects from HK (n = 3,324) and EUR GWAS 2 (n = 5,379). Standardized iHS scores based on the 1,000 Genomes Project were downloaded from a previous study^44^ for comparison.

### Calculation of polygenic risk scores

Polygenic risk scores (PRS) for individuals were computed using lassosum^47^, a penalized regression framework. The meta-analysis results on Europeans were used to calculate PRS for individuals of Chinese ancestry, and vice versa. LD information among SNPs was calculated from the testing dataset. These analyses were repeated using LDpred^48^.

The GZ SLE GWAS cohort was used as a test dataset to evaluate the influence of training data from different ancestries. Two predictors were constructed using lassosum based on meta-analysis results from: 1) HK and CC GWAS, 2,618 cases and 7,446 controls; 2) European GWAS, 4,576 cases and 8,039 controls. To control for influence from different sample sizes, 1,500 cases and 1,500 controls were randomly chosen from the Chinese and European populations to train two same-size predictors (repeated 3 times). PRS values generated from each test were scaled to a mean of 0 and a standard deviation of 1, and then evaluated based on the area under the ROC curve (AUC). The values and the 95% confidence intervals were calculated using the R package pROC^82^ and the optimal cut-off, the point that maximizes the sum of sensitivity and specificity, for case-control classification was estimated using the coords function.

## Acknowledgements

We thank grant support from National Key Research and Development Program of China (2017YFC0909001), Hong Kong PhD fellowship scheme (HKPF), HKU Postgraduate Scholarships and the Edward & Yolanda Wong Fund for supporting postgraduate students who participated in this work. We also thank the Hong Kong Area of Excellence (AoE) NPC case-control study partially funded by the World Cancer Research Fund (WCRF) for sharing their GWAS data. W. Yang and Y. L. Lau thank Research Grant Council of Hong Kong for support on genetic studies of SLE (GRF 17146616, 17125114, HKU783813M). Y. Zhang thanks grant support from National Natural Science Foundation of China (Grant No. 81601423). Y.-F. Wang thanks grant support from National Natural Science Foundation of China (Grant No. 81801636).

## Author contributions

W. Yang and Y.-F. Wang conceived the study. Y. L. Lau and Y. Zhang took the lead in data collection and management. Z. Lin, H. Zhang, Y. Sheng, X. Yin, S.-L. Zhong, X. Gu, J. He, Q. Wu, T.-H. Lam, J.-H. Lin, Z.-M. Mai, Y. Tang, Y. Chen, Q. Song, B. Ban, C.C. Mok, C. Yong, L. Lu, N. Shen, P.C. Sham, C.C. Lau, J. Yang and X. Zhang undertook subject recruitment and collected phenotype data. D.M. and T.J.V shared SLE GWAS data on European populations. Y.-F. Wang, T.-Y. Wang, Y. Cao, M. Guo, J.J. Shen carried out data analyses including quality control, genotype imputation, association and meta-analyses. Y.-F. Wang, T.-Y. Wang and Y. Lei carried out fine-mapping, selection signatures analyses and PRS comparison between the two ancestral groups. Y.-F. Wang, W. Yang, Y. Zhang, Y. L. Lau, P.C. Sham and D.K. Smith wrote the manuscript. All authors read and contributed to the manuscript.

