## Extended Data Figures for "Identification of 38 novel loci for systemic lupus erythematosus and genetic heterogeneity that may underly population disparities in this disease"

a

**HK GWAS**

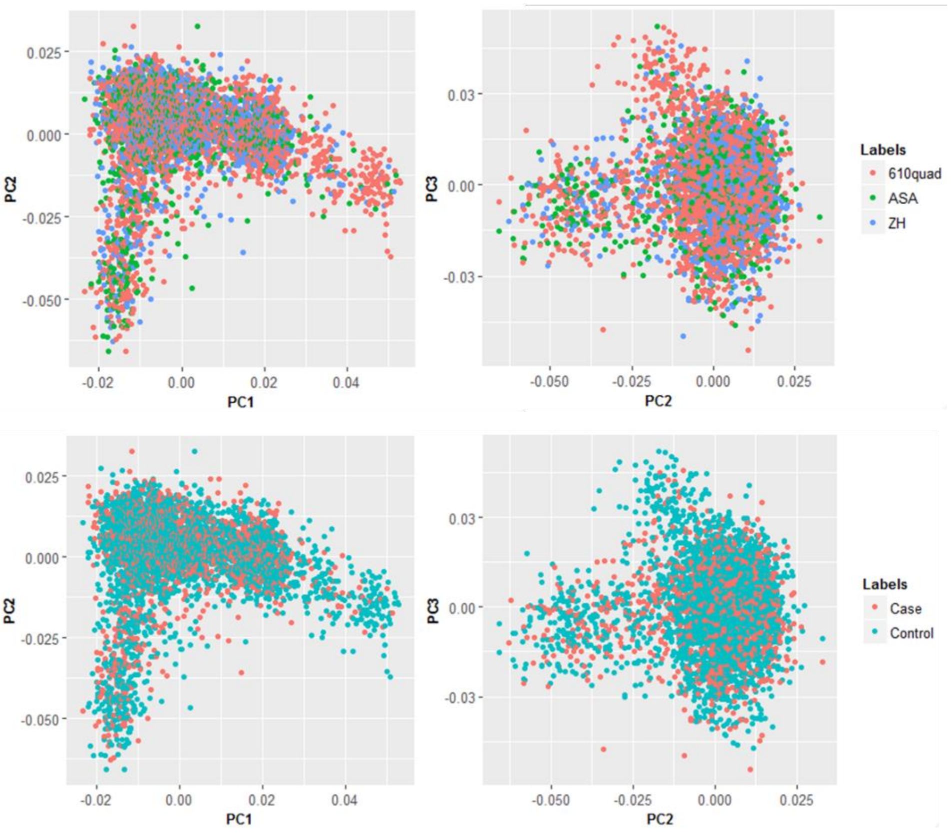

b

**GZ GWAS**

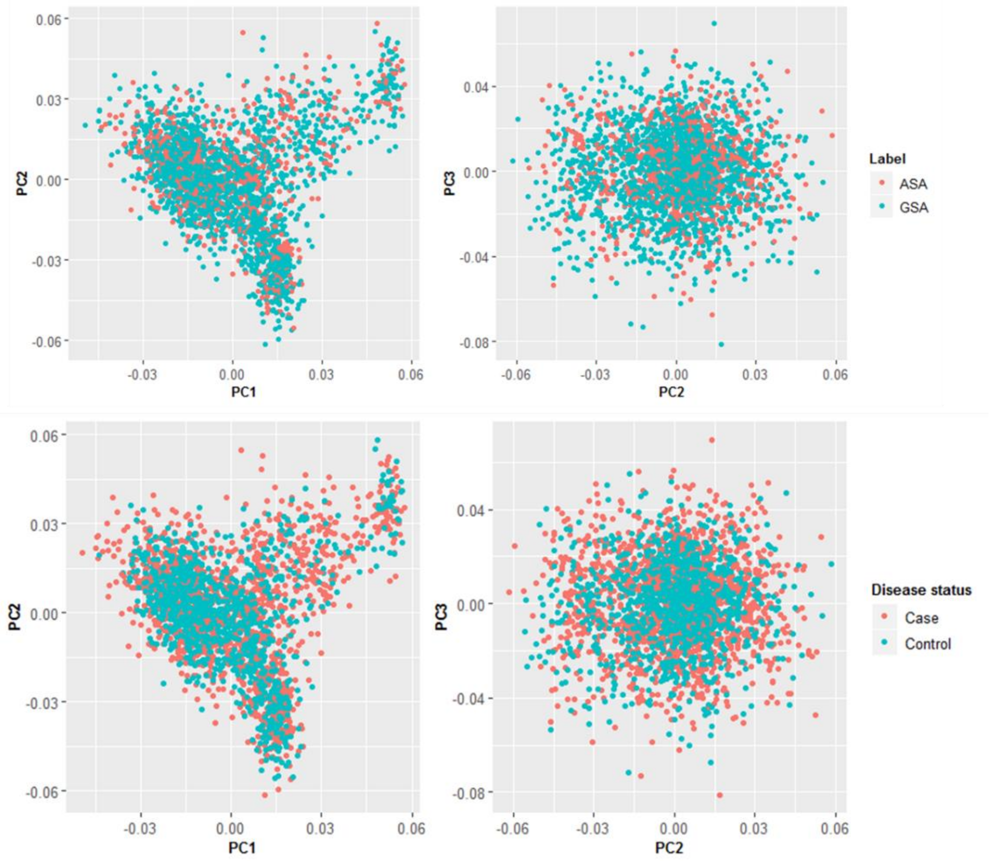

**c** CC GWAS

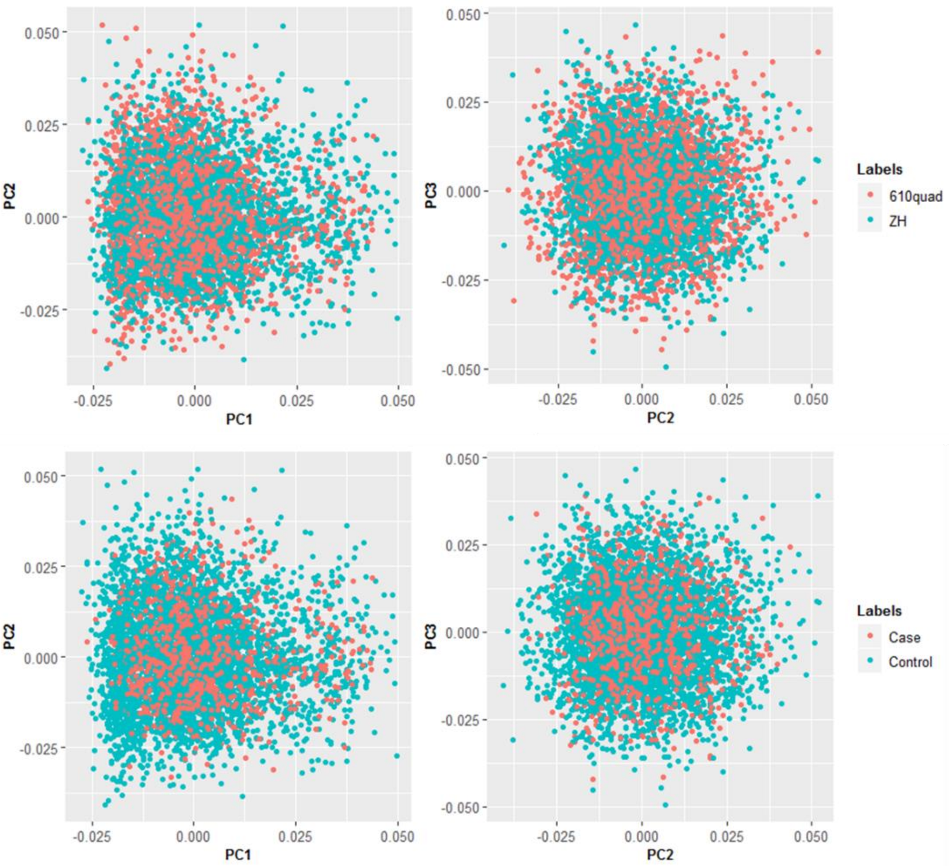

**Extended Data Fig. 1** Principal component (PC) analysis for individuals from Hong Kong (HK, **a**), Guangzhou (GZ, **b**) and Central China (CC, **c**) GWAS.

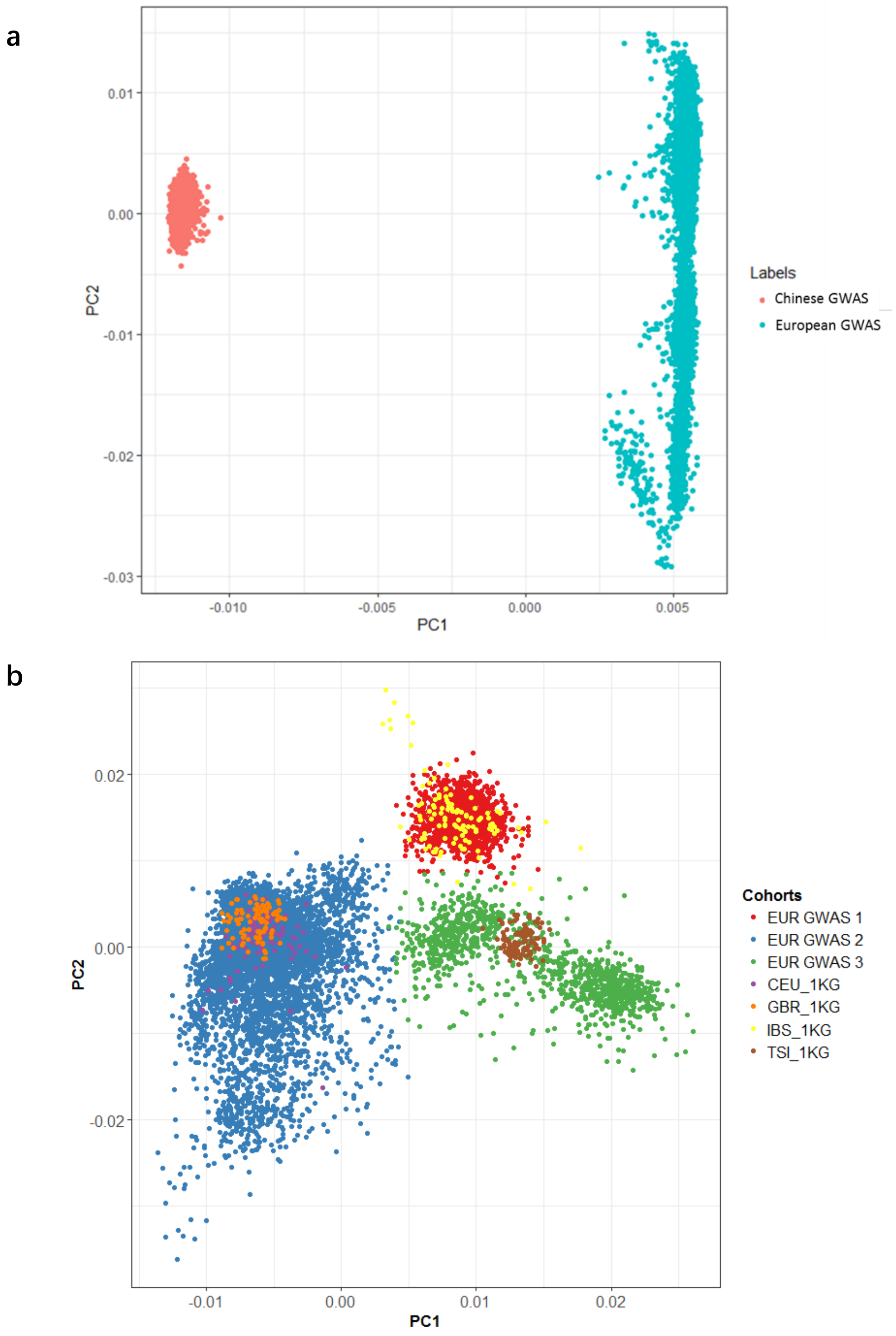

**Extended Data Fig. 2** PC analysis for individuals from the Chinese and European GWAS. **a**, comparison of PC for individuals of Chinese and European ancestries used in this study. **b**, Comparing the PCs for individuals in European GWAS with subjects from the 1,000 Genomes Project. Samples in EUR GWAS 1 overlapped individuals from Iberian population of Spain in the 1000 Genomes Project (IBS_1KG). Samples in EUR GWAS 2 overlapped individuals of northern and western European Ancestry (CEU_1KG and GBR_1KG). Samples in EUR GWAS 3 overlapped individuals from Toscani in Italy (ITS_1KG).

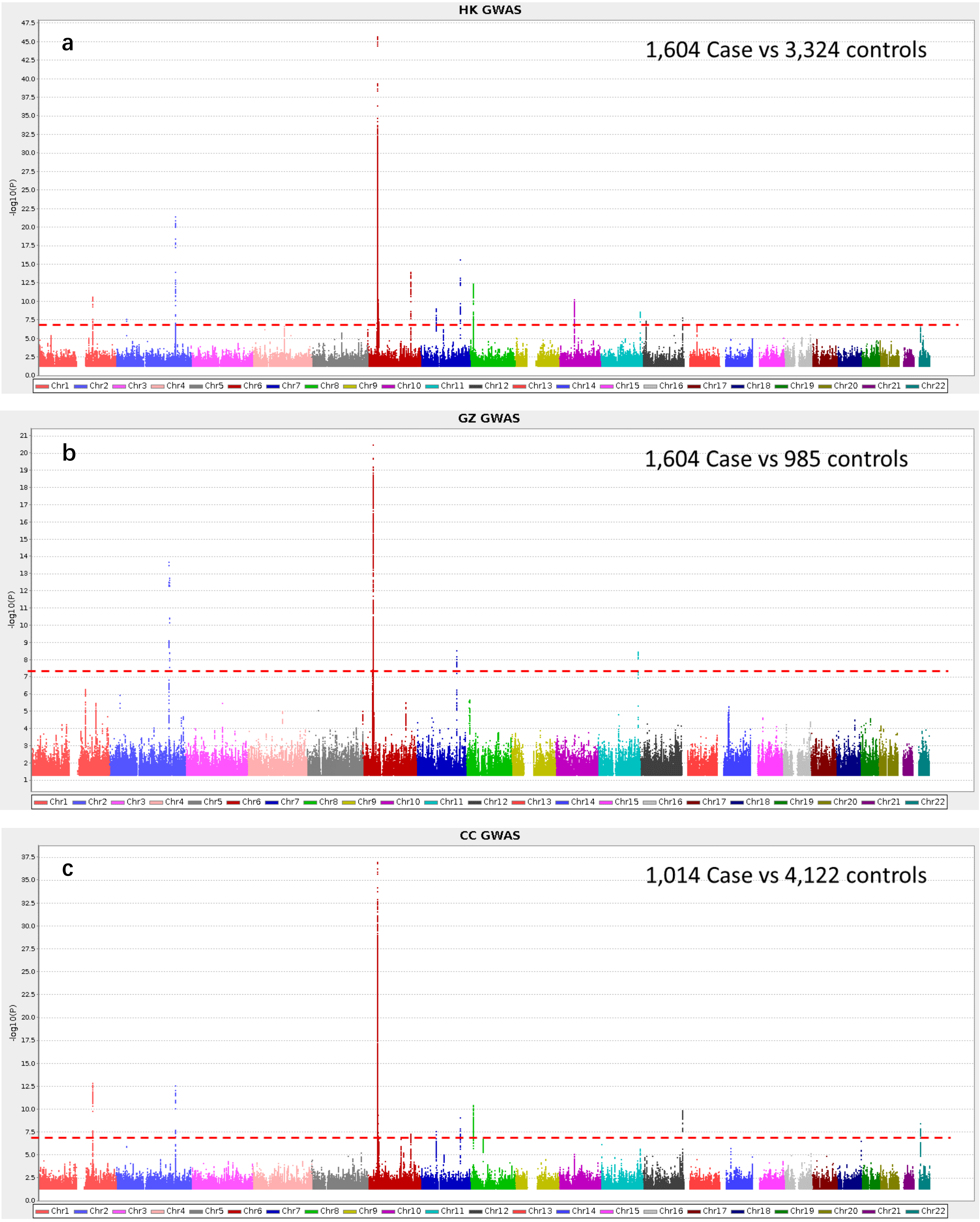

**Extended Data Fig. 3** Manhattan plots for the SLE GWAS from Chinese populations.

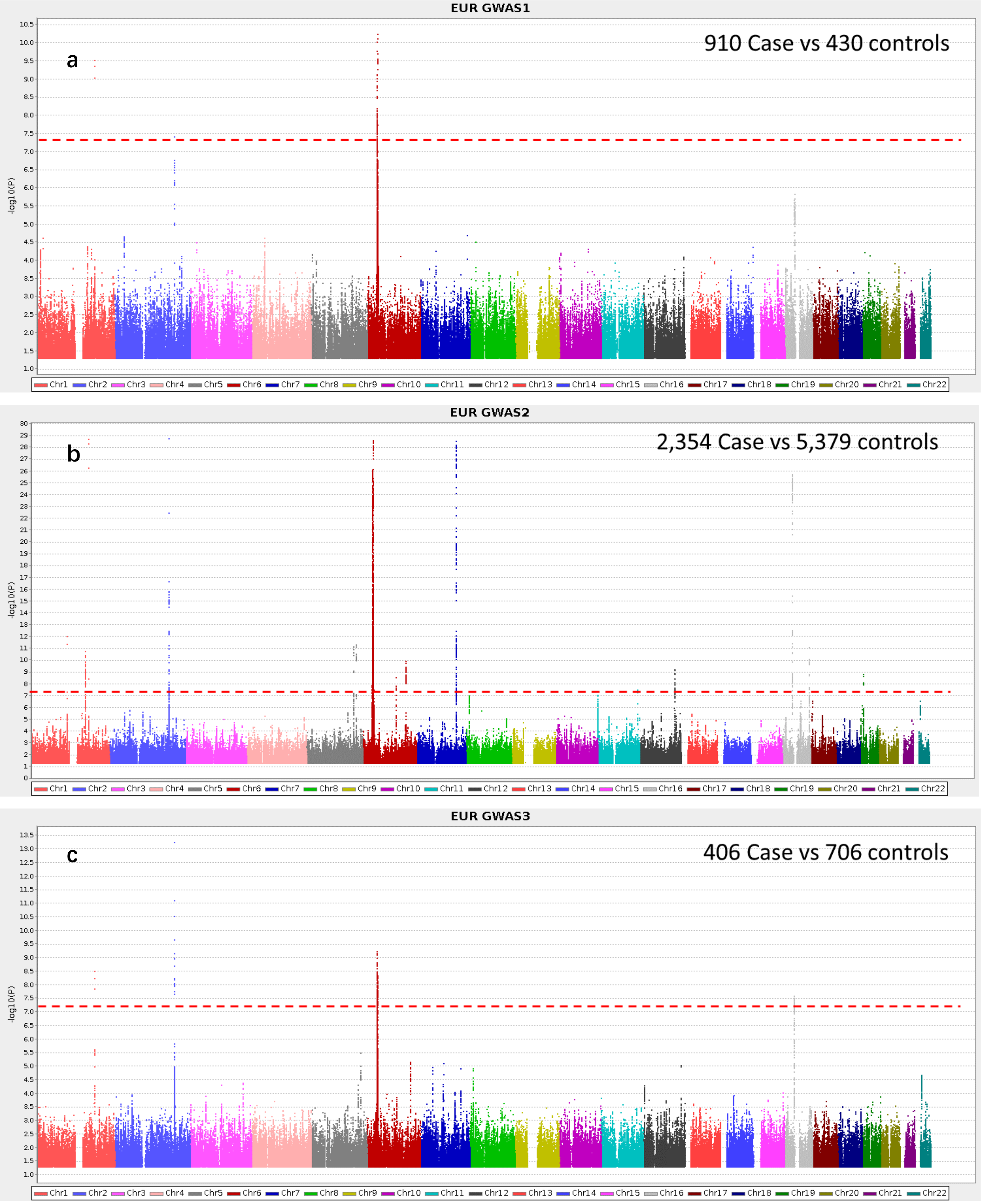
EUR GWAS1
a
910 Case vs 430 controls
-log10(P)
Chr1 Chr2 Chr3 Chr4 Chr5 Chr6 Chr7 Chr8 Chr9 Chr10 Chr11 Chr12 Chr13 Chr14 Chr15 Chr16 Chr17 Chr18 Chr19 Chr20 Chr21 Chr22
EUR GWAS2
b
2,354 Case vs 5,379 controls
-log10(P)
Chr1 Chr2 Chr3 Chr4 Chr5 Chr6 Chr7 Chr8 Chr9 Chr10 Chr11 Chr12 Chr13 Chr14 Chr15 Chr16 Chr17 Chr18 Chr19 Chr20 Chr21 Chr22
EUR GWAS3
c
406 Case vs 706 controls
-log10(P)
Chr1 Chr2 Chr3 Chr4 Chr5 Chr6 Chr7 Chr8 Chr9 Chr10 Chr11 Chr12 Chr13 Chr14 Chr15 Chr16 Chr17 Chr18 Chr19 Chr20 Chr21 Chr22

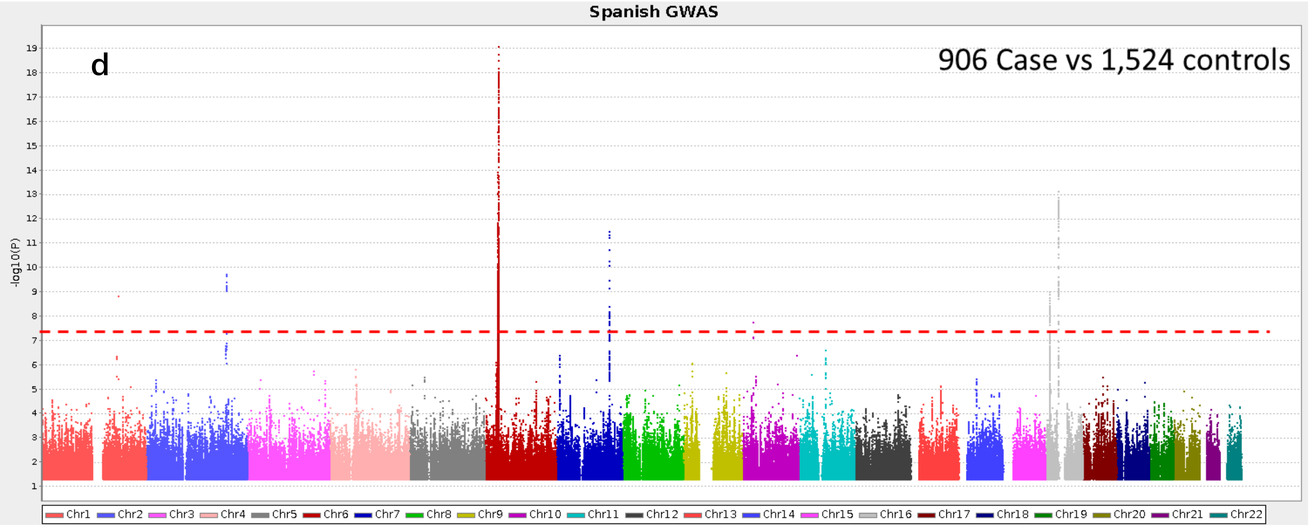

**Extended Data Fig. 4** Manhattan plots for the SLE GWAS from European populations.

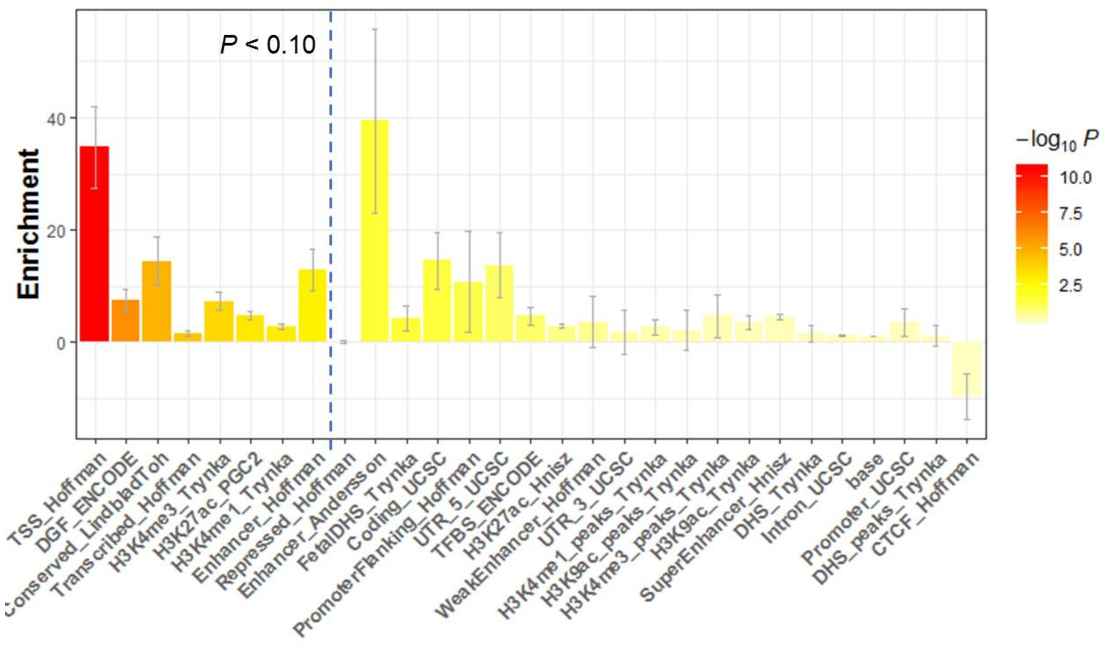

**Extended Data Fig. 5** Enrichment of SLE heritability across 28 core annotations (not specific for cell-type). Error bars represent jackknifed standard errors around the estimates of enrichment.

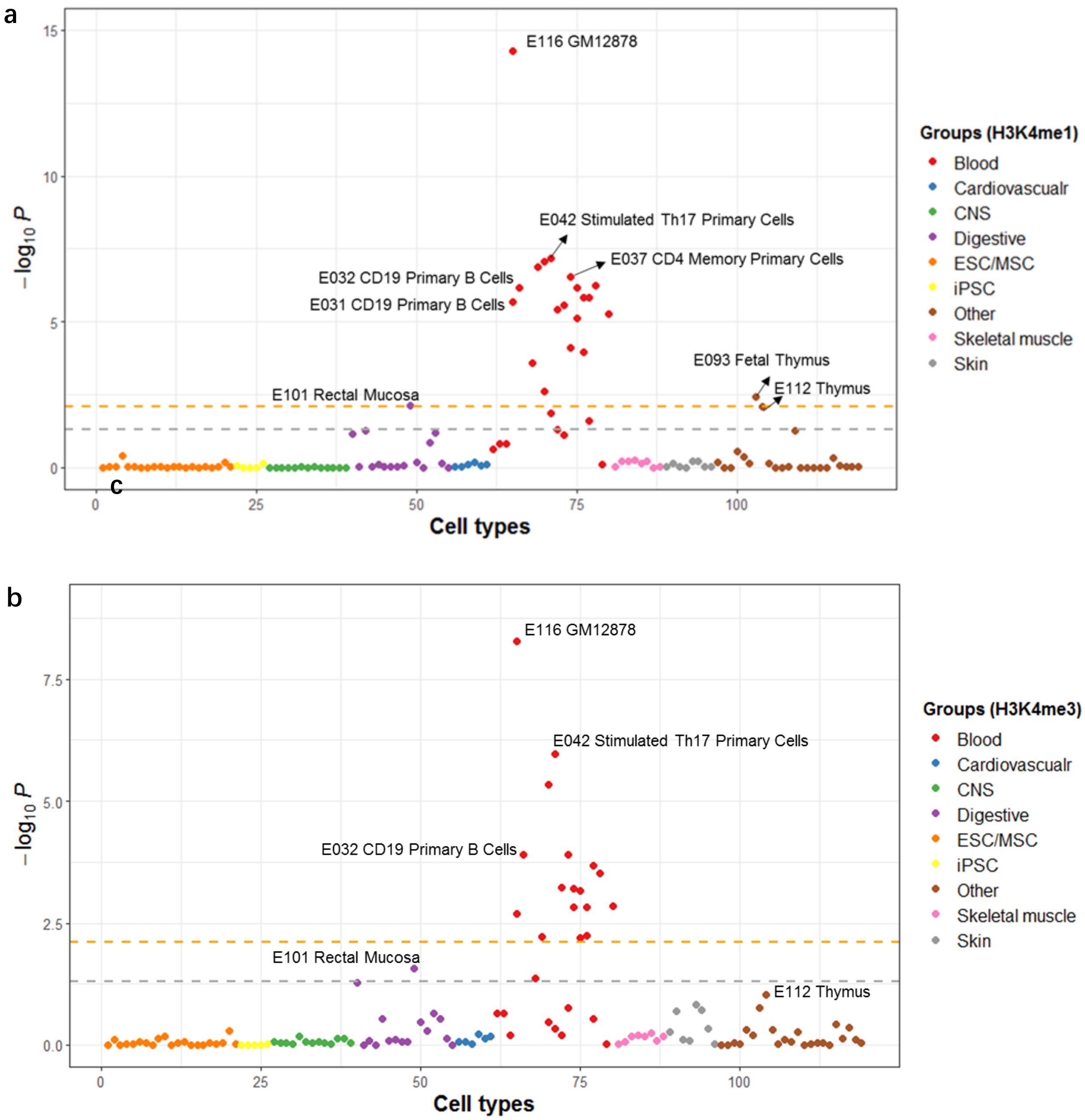

**Extended Data Fig. 6** Enrichment of SLE heritability across different cell types based on H3K4me1 **(a)** and H3K4me3 modifications **(b)**. Epigenomic data were obtained from the Roadmap Epigenomics Project and categorized into nine tissue/organ groups: blood, cardiovascular, gastrointestinal, central nervous system (CNS), skeletal muscle, skin, embryonic (ESC) or mesenchymal (MSC) stem cells, induced pluripotent stem cells (iPS) and others. The orange lines indicate a FDR threshold of 0.05 and the gray lines indicate a *P*-value threshold 0.05.

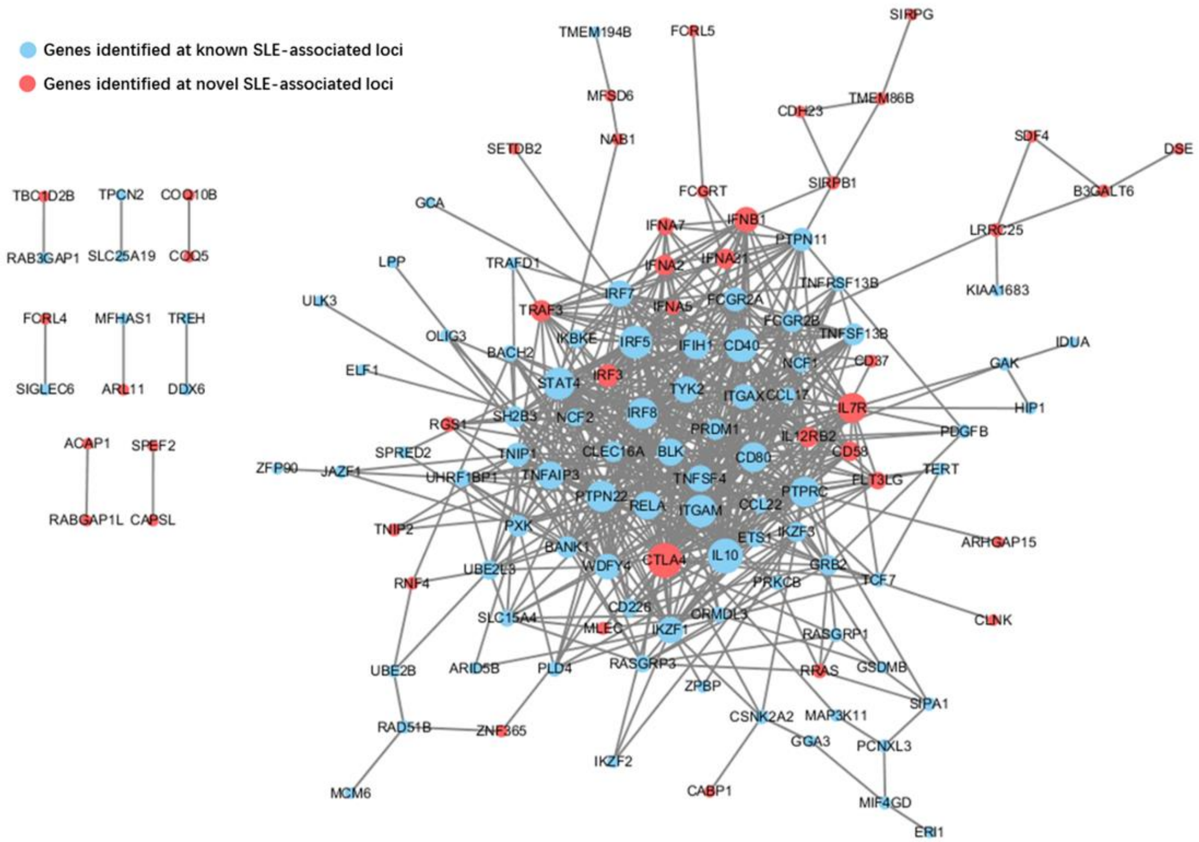

**Extended Data Fig. 7** Protein-Protein interaction (PPI) network corresponding to putative disease genes across SLE susceptibility loci. Genes identified at the novel SLE-associated loci are labeled in red, and genes found at the known SLE-associated loci are labeled in blue. The PPI information was retrieved from the STRING database (version 11.0).

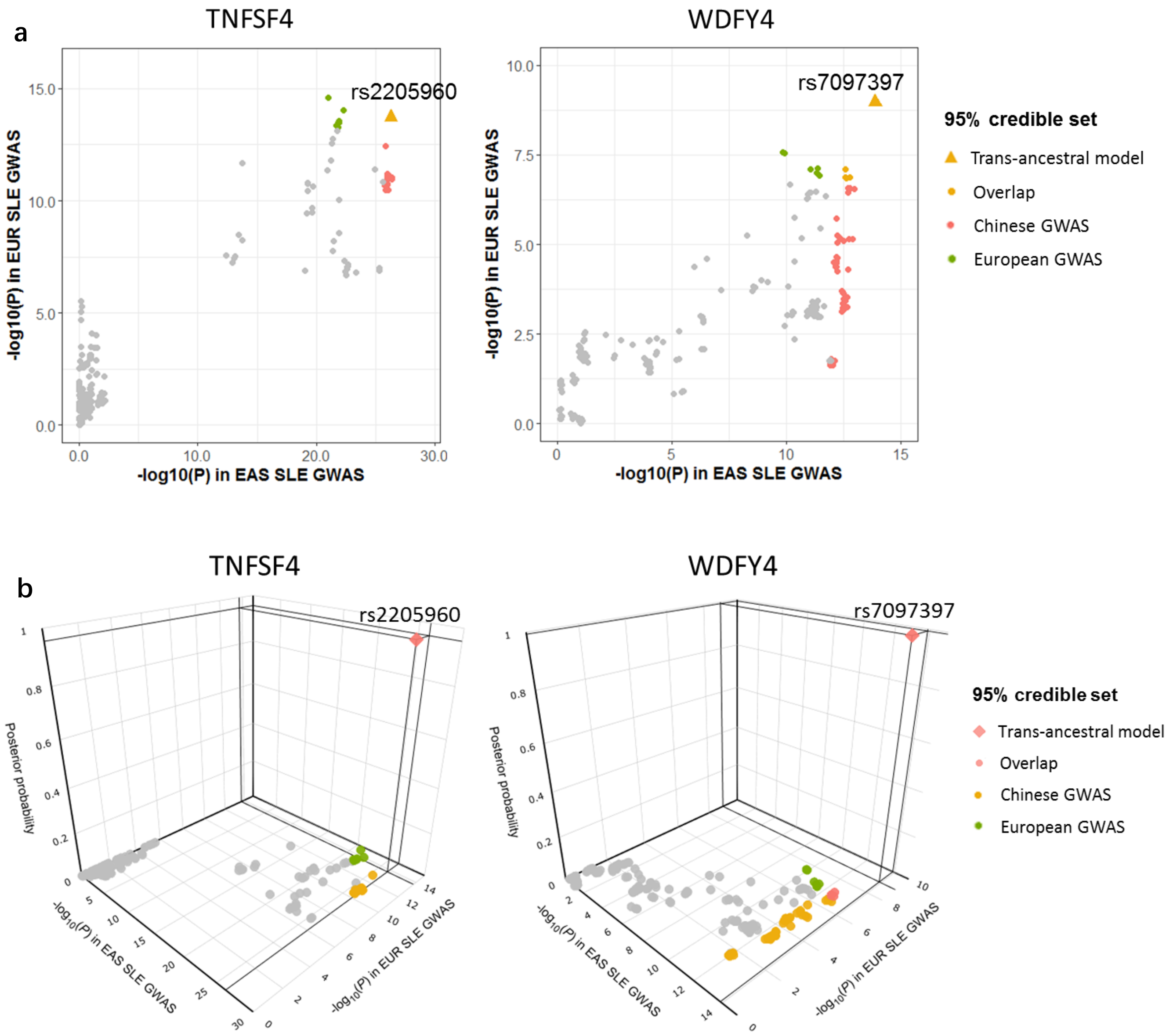

**Extended Data Fig. 8** Trans-ancestral fine-mapping results at the *TNFSF4* and *WDFY4* loci. **a,** X-axis indicates the log10 transformed association signal in the Chinese SLE GWAS, and Y-axis indicates the log10 transformed association signal in the European SLE GWAS. SNPs included in the 95% credible sets that were identified using the Chinese and European GWAS alone are labeled in red and green, respectively and yellow dots indicate variants found in both ancestral groups. The variant within 95% credible set that was identified using trans-ancestral fine-mapping model is labeled as a triangle. **b,** As in **a**, with the additional Z-axis showing the posterior probability of causality for each variant based on trans-ancestral fine-mapping model. For the analysis of the *TNFSF4* locus, the 95% credible set was reduced from 9 variants using the European GWAS alone to a single variant after the trans-ancestral analysis. The posterior probability of causality for the top variant, rs2205960, increased from 0.07 to 0.96, owing to the differential LD with other variants between the two ancestral groups. Similarly, a unique putative variant was also identified at the *WDFY4* locus (**Extended Data Table 7**).

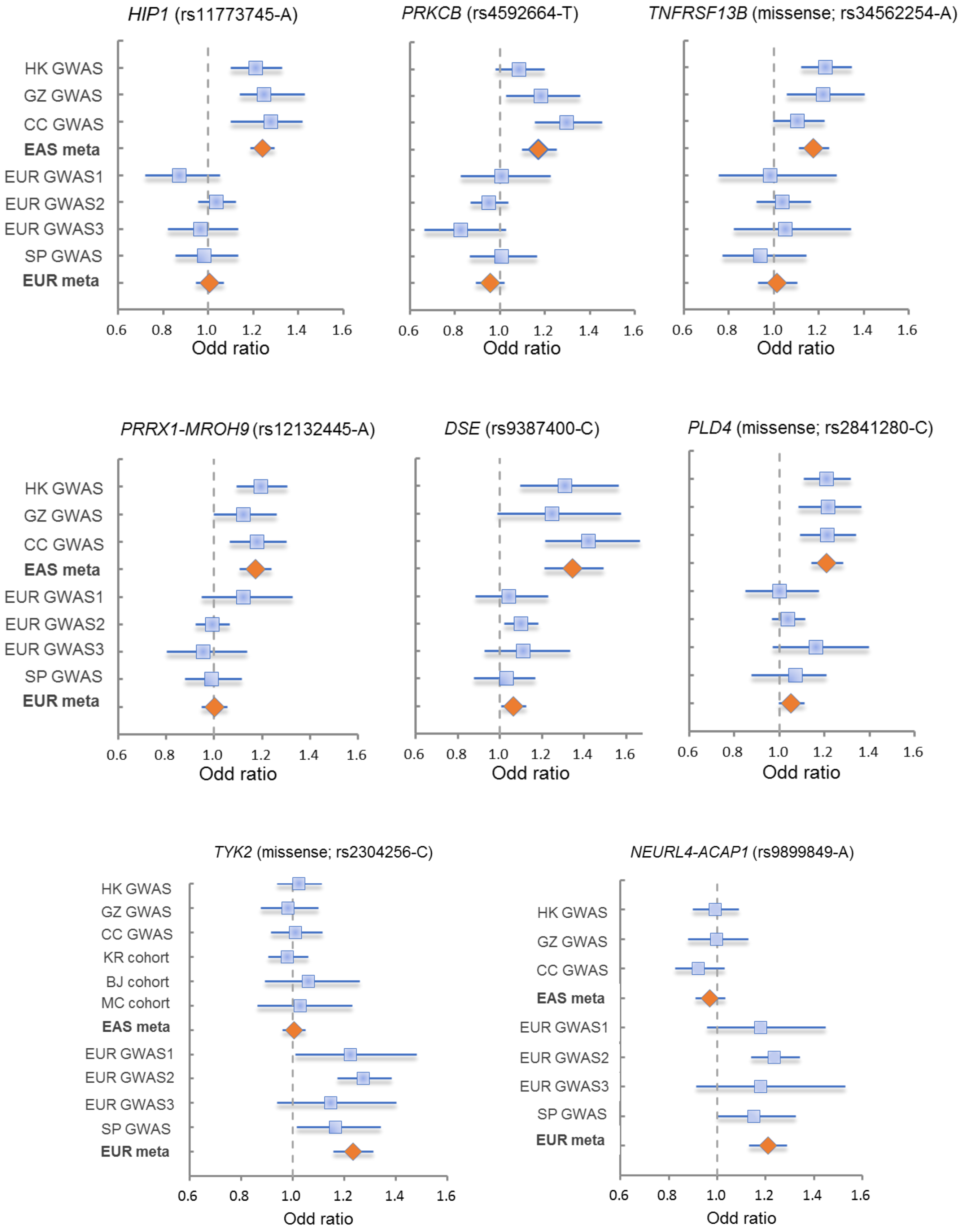

**Extended Data Fig. 9** Forest plots for the disease-associated loci with heterogeneity between East Asian (EAS) and European (EUR) populations.

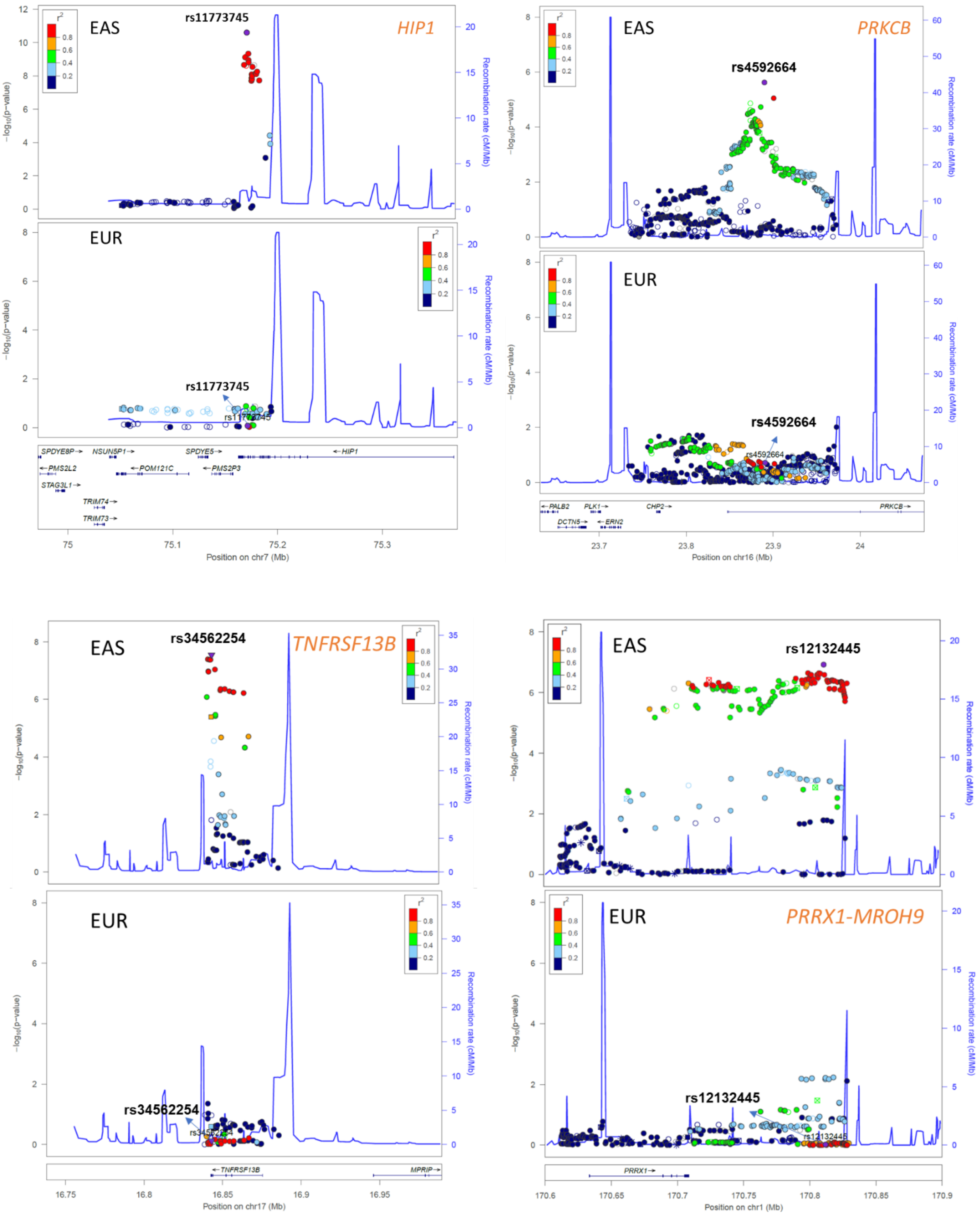
EAS
EUR
rs11773745
HIP1
Position on chr7 (Mb)
rs4592664
PRKCB
Position on chr16 (Mb)
rs34562254
TNFRSF13B
Position on chr17 (Mb)
rs12132445
PRRX1-MROH9
Position on chr1 (Mb)
−log10(p−value)
Recombination rate (cM/Mb)

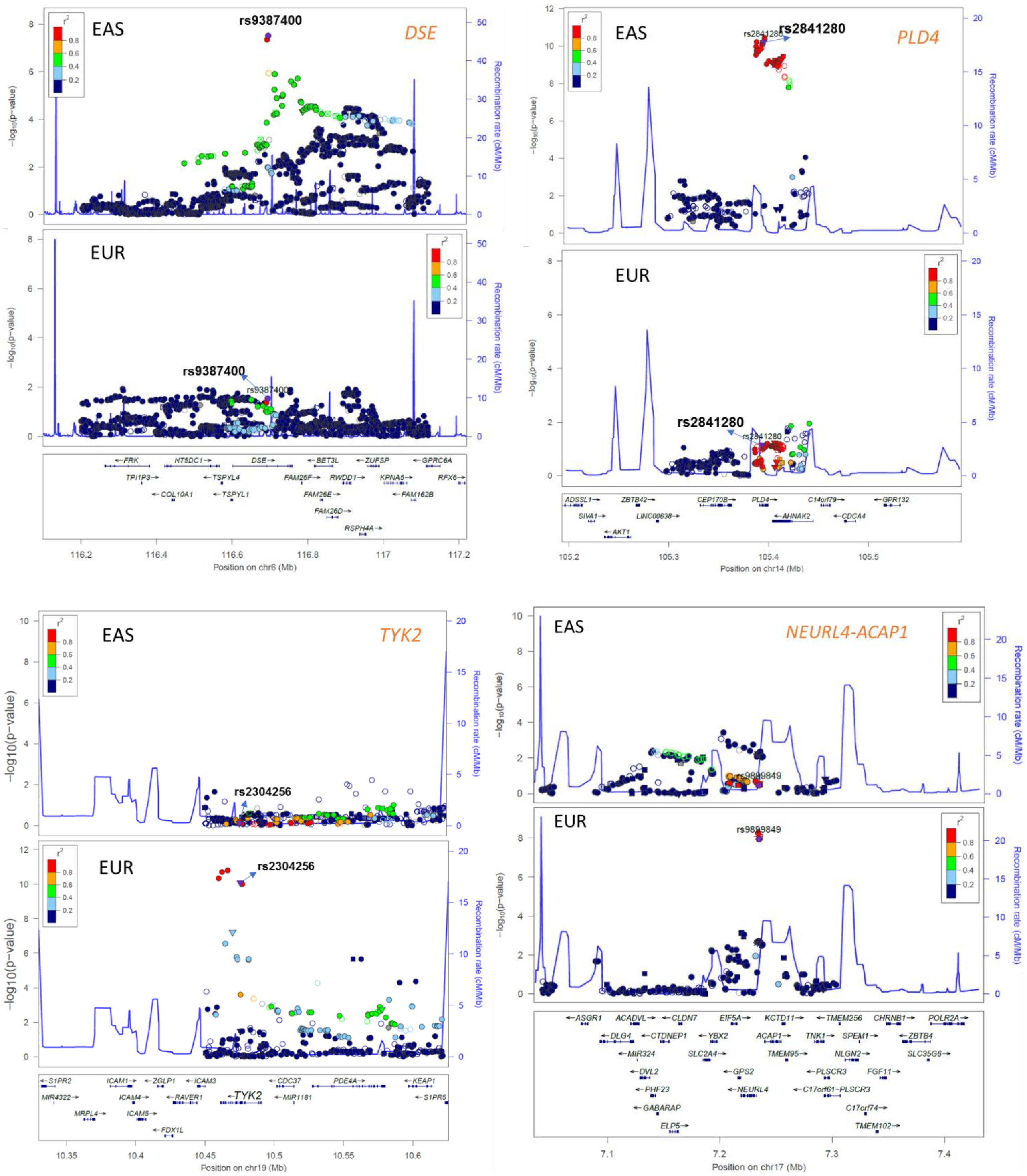

**Extended Data Fig. 10** Regional plots for the loci showing significant differences in effect-size estimates between the two ancestral groups.

a

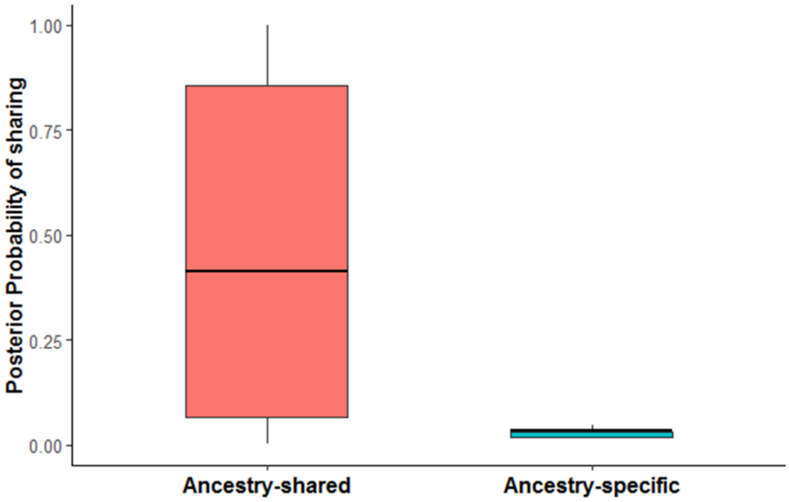

b

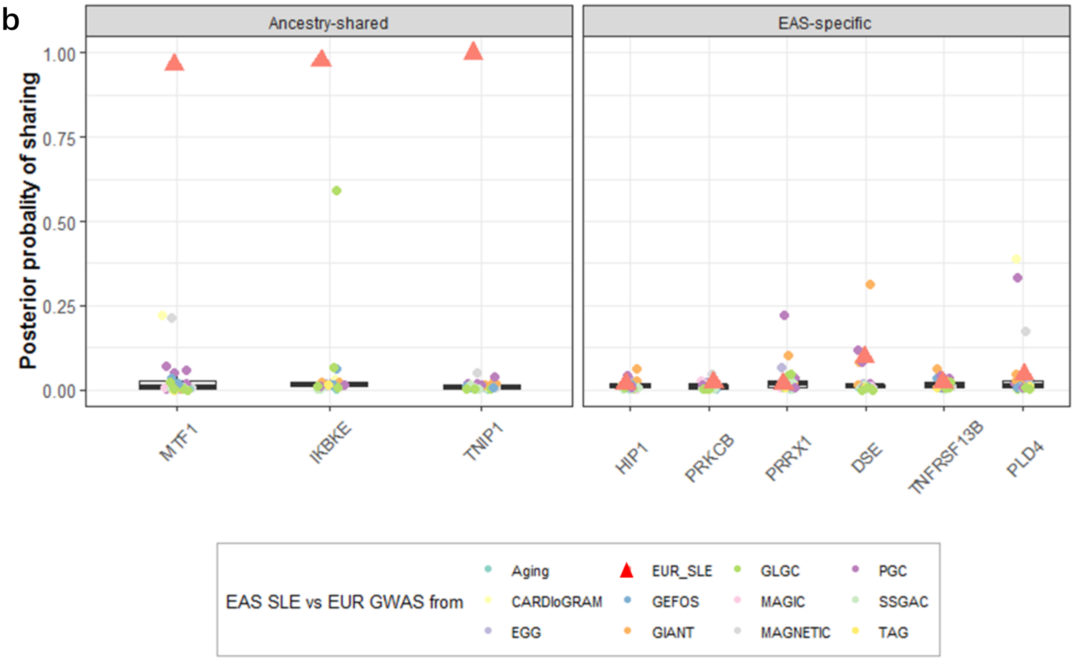

**Extended Data Fig. 11** Colocalization of East Asian (EAS) SLE association signals with European SLE associations and other non-immune system related diseases. **a**, Posterior probability of sharing for each ancestry-shared (left) and ancestry-specific SLE loci (right). **b**, Red triangles represent the posterior probability of SLE association signals between East Asian and European ancestries. Other symbols represent the colocalization of Chinese SLE association signals with those in European populations for 27 phenotypes unrelated to immunity. Summary statistics for the 27 phenotypes were downloaded from the results of 11 consortium studies: Aging, Global Lipids Genetics Consortium (GLGC), Psychiatric Genomics Consortium (PGC), Coronary Artery Disease Genome-wide Replication and Meta-analysis Consortium (CARDIoGRAM), Genetic Factors for Osteoporosis Consortium (GEFOS), Meta-analyses of Glucose and Insulin-related traits Consortium (MAGIC), Social Science Genetic Association Consortium (SSGAC), Early Growth Genetics (EGG), Genetic Investigation of Anthropometric Traits (GIANT), Myocardial Applied Genomics Network consortium (MAGNET) and Tobacco and Genetics Consortium (TAG).

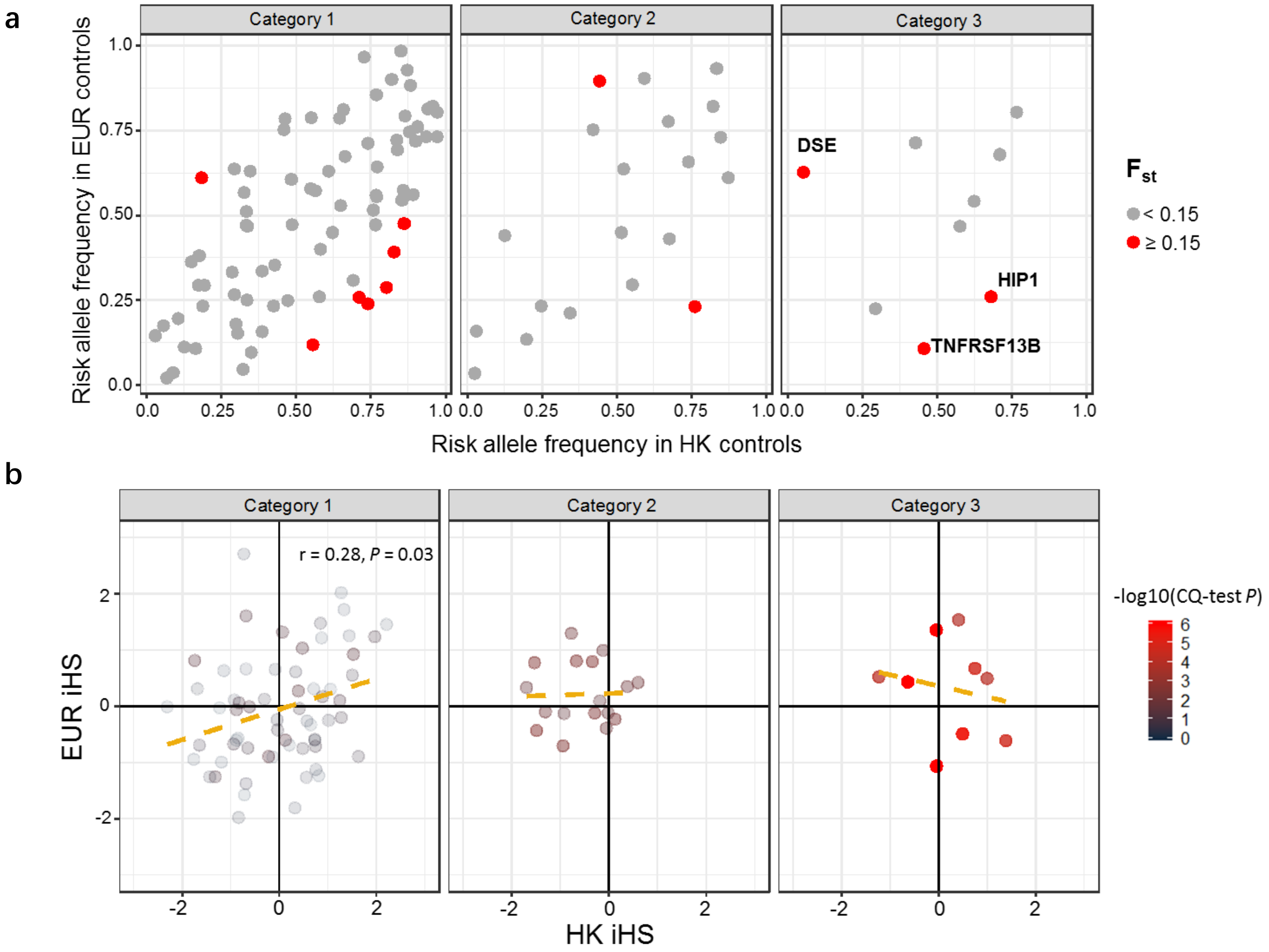

**Extended Data Fig. 12** Comparison of risk allele frequency and standardized iHS scores between East Asians and Europeans for SLE-associated variants. **a**. X-axis is the SLE risk allele frequency in a Hong Kong population (n = 3,324), and the y-axis is the risk allele frequency in EUR GWAS 2 (n = 5,379). The disease variants showing significant frequency variation ($F_{st}$ > 0.15) between ethnicities are shown in red. **b**. X-axis is the standardized iHS score estimated using data from a Hong Kong population (n = 3,324), y-axis is the score estimated using data from the European populations (n = 5,379).

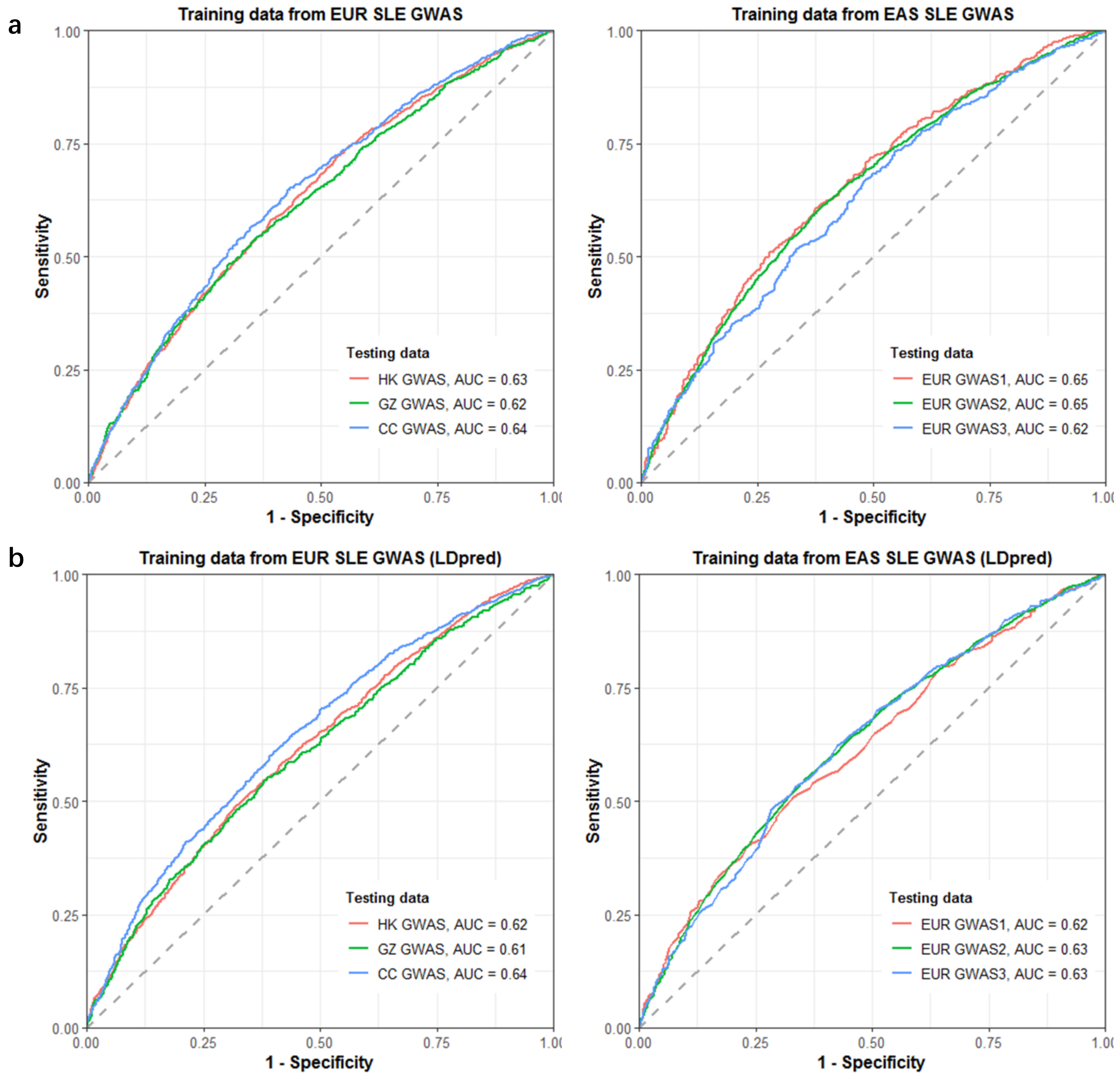

**Extended Data Fig. 13** Disease risk prediction accuracy based on polygenetic risk scores (PRS) between the two ancestral group populations. SLE PRS for individuals in the East Asian cohorts were calculated based on the summary statistics from the European GWAS (**left panel**), and vice versa (**right panel**). Data on a total of 4,222 SLE cases and 8,431 controls was included in the summary statistics of Chinese SLE GWAS, and data on a total of 4,576 cases and 8,039 controls were included in the summary statistics of European SLE GWAS. **a**. Performance of PRS calculated by lassosum for individuals from different cohorts; **b**. Performance of PRS calculated by LDpred, another algorithm for PRS estimation.

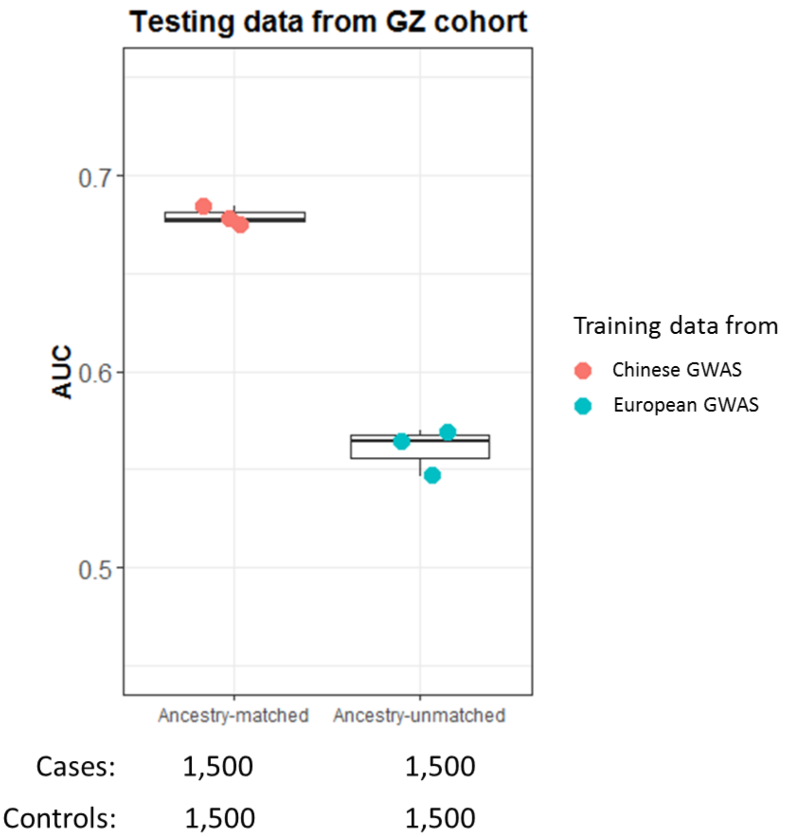

**Extended Data Fig. 14** Performance of PRS for GZ samples based on data from Chinese and European populations with equivalent sample size. To control for the influence of sample size 1,500 cases and 1,500 controls were randomly selected to train the predictors for the populations (**red**, Chinese; **blue**, European). These procedures were repeated 3 times, and the performance measured by AUC for each test is represented by the dot.
