## Extended Data Tables for "Identification of 38 novel loci for systemic lupus erythematosus and genetic heterogeneity that may underly population disparities in this disease"

**Extended Data Table 1** Evaluation of genotyping accuracy between different BeadChips

| Sample | BeadChips | #discordant SNPs | #overlapped SNPs | %concordant rate |
|---|---|---|---|---|
| 1 | 610-Quad vs ZhongHua | 11 | 293103 | 100.00% |
| 2 | 610-Quad vs ZhongHua | 10 | 293113 | 100.00% |
| 3 | 610-Quad vs ZhongHua | 35 | 292529 | 99.99% |
| 4 | 610-Quad vs ZhongHua | 10 | 292855 | 100.00% |
| 5 | 610-Quad vs ZhongHua | 5 | 293078 | 100.00% |
| 6 | 610-Quad vs ZhongHua | 28 | 292571 | 99.99% |
| 7 | 610-Quad vs ZhongHua | 4 | 293110 | 100.00% |
| 8 | 610-Quad vs ZhongHua | 10 | 293119 | 100.00% |
| 9 | GSA vs ASA | 1 | 145804 | 100.00% |
| 10 | GSA vs ASA | 6 | 145722 | 100.00% |
| 11 | GSA vs ASA | 9 | 145626 | 99.99% |
| 12 | GSA vs ASA | 8 | 145542 | 99.99% |
| 13 | GSA vs ASA | 7 | 145616 | 100.00% |
| 14 | GSA vs ASA | 5 | 145537 | 100.00% |

**Extended Data Table 2** Summary of SLE cohorts from East Asian (EAS) and European (EUR) populations

| Cohorts | Ancestry | #Case | #Controls | #Total |
|---|---|---|---|---|
| HK GWAS | EAS | 1,604 | 3,324 | 4,928 |
| GZ GWAS | EAS | 1,604 | 985 | 2,589 |
| CC GWAS | EAS | 1,014 | 4,122 | 5,136 |
| KR cohort | EAS | 1,710 | 6,836 | 8,546 |
| BJ cohort | EAS | 490 | 493 | 983 |
| MC cohort | EAS | 285 | 287 | 572 |
| EUR GWAS1 | EUR | 910 | 430 | 1,340 |
| EUR GWAS2 | EUR | 2,354 | 5,379 | 7,733 |
| EUR GWAS3 | EUR | 406 | 706 | 1,112 |
| SP GWAS | EUR | 906 | 1524 | 2,430 |
| **Total** | | **11,283** | **24,086** | **35,369** |

HK: Hong Kong; GZ: Guangzhou; CC: Central China; KR: Korea; BJ: Beijing; MC: Chinese in Malaysia; SP: Spain

**Extended Data Table 3** Summary association statistics of reported SLE-associated variants across different genetic cohorts

| Order | rsid | Chr | Pos (hg19) | AlleleA | AlleleB | Gene context | Annotation | Cohorts | FA AlleleB | FU AlleleB | OR | SE | P | Cochran's Q-test *P* | References (PMID) |
|---|---|---|---|---|---|---|---|---|---|---|---|---|---|---|---|
| **#1** | **rs2476601** | **1** | **114377568** | **A** | **G** | **PTPN22** | **missense** | **EUR meta** | **-** | **-** | **0.713** | **0.045** | **3.04E-14** | Mononorphic | (1)19838195, |
| | | | | | | | | *EUR GWAS1* | *0.909* | *0.907* | *1.028* | *0.141* | *8.46E-01* | SNV in EAS | (2)26502338 |
| | | | | | | | | *EUR GWAS2* | *0.858* | *0.901* | *0.676* | *0.055* | *9.48E-13* | | |
| | | | | | | | | *EUR GWAS3* | *0.917* | *0.941* | *0.687* | *0.171* | *2.80E-02* | | |
| | | | | | | | | *SP GWAS* | *-* | *-* | *0.720* | *0.106* | *7.03E-04* | | |
| **#2** | **rs1801274** | **1** | **161479745** | **A** | **G** | **FCGR2A** | **missense** | **Trans meta** | **-** | **-** | ***1.167*** | ***0.020*** | **5.05E-15** | 0.076 | 26502338 |
| | | | | | | | | EAS meta | - | - | *1.127* | *0.028* | 1.79E-05 | | |
| | | | | | | | | *HK GWAS* | *0.368* | *0.334* | *1.169* | *0.045* | *5.88E-04* | | |
| | | | | | | | | *CC GWAS* | *0.362* | *0.346* | *1.063* | *0.052* | *2.40E-01* | | |
| | | | | | | | | *GZ GWAS* | *0.373* | *0.342* | *1.160* | *0.062* | *1.70E-02* | | |
| | | | | | | | | EUR meta | - | - | 1.209 | 0.028 | 1.19E-11 | | |
| | | | | | | | | *EUR GWAS1* | *0.577* | *0.502* | *1.362* | *0.084* | *2.19E-04* | | |
| | | | | | | | | *EUR GWAS2* | *0.547* | *0.511* | *1.159* | *0.037* | *6.20E-05* | | |
| | | | | | | | | *EUR GWAS3* | *0.461* | *0.419* | *1.180* | *0.088* | *5.97E-02* | | |
| | | | | | | | | *SP GWAS* | *-* | *-* | *1.285* | *0.060* | *1.85E-04* | | |
| **#3** | **rs35426045** | **1** | **161649724** | **G** | **A** | **FCGR2B** | **intergenic** | **EAS meta** | **-** | **-** | **1.224** | **0.040** | **5.21E-07** | Rare variant in | 26808113 |
| | | | | | | | | *HK GWAS* | *0.856* | *0.824* | *1.285* | *0.060* | *3.27E-05* | EUR | |
| | | | | | | | | *CC GWAS* | *0.867* | *0.847* | *1.186* | *0.073* | *1.90E-02* | | |
| | | | | | | | | *GZ GWAS* | *0.869* | *0.849* | *1.167* | *0.082* | *5.78E-02* | | |
| **#4** | **rs2205960** | **1** | **173191475** | **G** | **T** | **TNFSF4** | **intergenic** | **Trans meta** | **-** | **-** | **1.368** | **0.020** | **2.52E-57** | 0.011 | (1)19838193 |
| | | | | | | | | EAS meta | - | - | 1.422 | 0.025 | 5.82E-46 | | (2)19838195 |
| | | | | | | | | *HK GWAS* | *0.310* | *0.248* | *1.365* | *0.048* | *8.95E-11* | | (3)22820624, |

| | | | | | | | | | | | | | | | |
|---|---|---|---|---|---|---|---|---|---|---|---|---|---|---|---|
| | | | | | | | | *CC GWAS* | *0.339* | *0.259* | *1.492* | *0.054* | *1.52E-13* | | (4)23874208 |
| | | | | | | | | *GZ GWAS* | *0.316* | *0.254* | *1.349* | *0.065* | *3.95E-06* | | |
| | | | | | | | | *KR IC* | *0.313* | *0.237* | *1.470* | *0.046* | *5.79E-16* | | |
| | | | | | | | | *BJ IC* | *0.340* | *0.249* | *1.560* | *0.098* | *8.85E-06* | | |
| | | | | | | | | *MC IC* | *0.289* | *0.254* | *1.190* | *0.135* | *1.82E-01* | | |
| | | | | | | | | EUR meta | - | - | 1.281 | 0.032 | 1.92E-14 | | |
| | | | | | | | | *EUR GWAS1* | *0.248* | *0.189* | *1.409* | *0.105* | *1.12E-03* | | |
| | | | | | | | | *EUR GWAS2* | *0.278* | *0.232* | *1.302* | *0.042* | *2.38E-10* | | |
| | | | | | | | | *EUR GWAS3* | *0.244* | *0.204* | *1.271* | *0.106* | *2.38E-02* | | |
| | | | | | | | | *SP GWAS* | - | - | *1.176* | *0.071* | *2.01E-02* | | |
| **#5** | **rs1418190** | **1** | **173361979** | **C** | **T** | **LOC100506023** | **ncRNA_intronic** | **Trans meta** | **-** | **-** | **1.223** | **0.021** | **2.14E-22** | 0.087 | 25890262 |
| | | | | | | | | EAS meta | - | - | 1.268 | 0.030 | 1.13E-15 | | |
| | | | | | | | | *HK GWAS* | *0.596* | *0.549* | *1.230* | *0.044* | *3.20E-06* | | |
| | | | | | | | | *CC GWAS* | *0.693* | *0.636* | *1.322* | *0.054* | *2.50E-07* | | |
| | | | | | | | | *GZ GWAS* | *0.609* | *0.550* | *1.275* | *0.059* | *3.84E-05* | | |
| | | | | | | | | EUR meta | - | - | 1.182 | 0.029 | 7.08E-09 | | |
| | | | | | | | | *EUR GWAS1* | *0.651* | *0.595* | *1.281* | *0.088* | *5.03E-03* | | |
| | | | | | | | | *EUR GWAS2* | *0.614* | *0.579* | *1.173* | *0.038* | *2.60E-05* | | |
| | | | | | | | | *EUR GWAS3* | *0.663* | *0.613* | *1.234* | *0.092* | *2.30E-02* | | |
| | | | | | | | | *SP GWAS* | - | - | *1.137* | *0.062* | *2.29E-02* | | |
| **#6** | **rs17849501** | **1** | **183542323** | **C** | **T** | **NCF2** | **synonymous** | **EUR meta** | **-** | **-** | **2.195** | **0.056** | **1.50E-45** | Mononorphic | 26502338 |
| | | | | | | | | *EUR GWAS1* | *0.171* | *0.067* | *2.455* | *0.143* | *3.07E-10* | SNV in EAS | |
| | | | | | | | | *EUR GWAS2* | *0.102* | *0.048* | *2.123* | *0.067* | *1.99E-29* | | |
| | | | | | | | | *EUR GWAS3* | *0.150* | *0.062* | *2.275* | *0.139* | *3.13E-09* | | |
| **#7** | **rs34889541** | **1** | **198594769** | **G** | **A** | **ATP6V1G3,** | **intergenic** | **Trans meta** | **-** | **-** | **0.833** | **0.035** | **1.59E-07** | 0.538 | 27399966 |

| | | | | | | | | | | | | | | | |
|---|---|---|---|---|---|---|---|---|---|---|---|---|---|---|---|
| | | | | | | **PTPRC** | | EAS meta | - | - | 0.847 | 0.044 | 1.38E-04 | | |
| | | | | | | | | *HK GWAS* | *0.111* | *0.128* | *0.836* | *0.069* | *9.41E-03* | | |
| | | | | | | | | *CC GWAS* | *0.131* | *0.157* | *0.811* | *0.073* | *4.26E-03* | | |
| | | | | | | | | *GZ GWAS* | *0.119* | *0.127* | *0.920* | *0.088* | *3.48E-01* | | |
| | | | | | | | | EUR meta | - | - | 0.810 | 0.058 | 2.61E-04 | | |
| | | | | | | | | *EUR GWAS1* | *0.061* | *0.067* | *0.914* | *0.172* | *6.00E-01* | | |
| | | | | | | | | *EUR GWAS2* | *0.056* | *0.072* | *0.758* | *0.078* | *3.56E-04* | | |
| | | | | | | | | *EUR GWAS3* | *0.076* | *0.077* | *0.989* | *0.166* | *9.47E-01* | | |
| | | | | | | | | *SP GWAS* | - | - | *0.806* | *0.125* | *1.80E-01* | | |
| **#8** | **rs2297550** | **1** | **206643772** | **C** | **G** | **IKBKE** | **UTR5** | **Trans meta** | **-** | **-** | **1.187** | **0.024** | **6.22E-13** | 0.676 | 27399966 |
| | | | | | | | | EAS meta | - | - | 1.179 | 0.029 | 1.57E-08 | | |
| | | | | | | | | *HK GWAS* | *0.585* | *0.557* | *1.119* | *0.044* | *1.07E-02* | | |
| | | | | | | | | *CC GWAS* | *0.566* | *0.526* | *1.168* | *0.051* | *2.14E-03* | | |
| | | | | | | | | *GZ GWAS* | *0.603* | *0.542* | *1.317* | *0.060* | *4.91E-06* | | |
| | | | | | | | | EUR meta | - | - | 1.204 | 0.042 | 7.82E-06 | | |
| | | | | | | | | *EUR GWAS1* | *0.140* | *0.139* | *1.009* | *0.122* | *9.38E-01* | | |
| | | | | | | | | *EUR GWAS2* | *0.132* | *0.119* | *1.177* | *0.056* | *3.55E-03* | | |
| | | | | | | | | *EUR GWAS3* | *0.150* | *0.123* | *1.266* | *0.131* | *7.17E-02* | | |
| | | | | | | | | *SP GWAS* | - | - | *1.357* | *0.086* | *6.07E-04* | | |
| **#9** | **rs3024505** | **1** | **206939904** | **G** | **A** | **IL10** | intergenic | Trans meta | - | - | 1.141 | 0.033 | 5.26E-05 | 0.043 | (1)19838195, |
| | | | | | | | | EAS meta | - | - | 1.021 | 0.064 | 7.41E-01 | | (2)26502338 |
| | | | | | | | | *HK GWAS* | *0.026* | *0.030* | *0.895* | *0.132* | *3.99E-01* | | |
| | | | | | | | | *CC GWAS* | *0.044* | *0.041* | *1.082* | *0.122* | *5.19E-01* | | |
| | | | | | | | | *GZ GWAS* | *0.028* | *0.025* | *1.118* | *0.179* | *5.33E-01* | | |
| | | | | | | | | *KR IC* | *0.028* | *0.027* | *1.060* | *0.127* | *6.72E-01* | | |

| | | | | | | | | | | | | | | | |
|---|---|---|---|---|---|---|---|---|---|---|---|---|---|---|---|
| | | | | | | | | BJ IC | 0.045 | 0.047 | 0.960 | 0.217 | 8.52E-01 | | |
| | | | | | | | | MC IC | 0.026 | 0.026 | 1.010 | 0.369 | 9.85E-01 | | |
| | | | | | | | | **EUR meta** | **-** | **-** | **1.186** | **0.038** | **6.53E-06** | | |
| | | | | | | | | EUR GWAS1 | 0.160 | 0.121 | 1.357 | 0.120 | 1.12E-02 | | |
| | | | | | | | | EUR GWAS2 | 0.175 | 0.158 | 1.114 | 0.049 | 2.64E-02 | | |
| | | | | | | | | EUR GWAS3 | 0.165 | 0.150 | 1.121 | 0.122 | 3.49E-01 | | |
| | | | | | | | | SP GWAS | - | - | 1.375 | 0.084 | 1.17E-04 | | |
| #10 | rs9782955 | 1 | 236039877 | T | C | LYST | intronic | **Trans meta** | **-** | **-** | **1.132** | **0.027** | **3.71E-06** | 0.232 | 26502338 |
| | | | | | | | | EAS meta | - | - | 1.184 | 0.046 | 2.60E-04 | | |
| | | | | | | | | HK GWAS | 0.901 | 0.880 | 1.244 | 0.071 | 1.97E-03 | | |
| | | | | | | | | CC GWAS | 0.894 | 0.884 | 1.108 | 0.080 | 1.96E-01 | | |
| | | | | | | | | GZ GWAS | 0.897 | 0.882 | 1.190 | 0.096 | 7.03E-02 | | |
| | | | | | | | | EUR meta | - | - | 1.107 | 0.033 | 2.05E-03 | | |
| | | | | | | | | EUR GWAS1 | 0.799 | 0.786 | 1.072 | 0.103 | 5.00E-01 | | |
| | | | | | | | | EUR GWAS2 | 0.767 | 0.746 | 1.123 | 0.042 | 6.40E-03 | | |
| | | | | | | | | EUR GWAS3 | 0.744 | 0.756 | 0.940 | 0.103 | 5.45E-01 | | |
| | | | | | | | | SP GWAS | - | - | 1.173 | 0.074 | 2.90E-02 | | |
| #11 | rs7579944 | 2 | 30445026 | T | C | LBH | intergenic | **Trans meta** | **-** | **-** | **1.155** | **0.018** | **1.17E-15** | 0.607 | 27399966 |
| | | | | | | | | EAS meta | - | - | 1.163 | 0.023 | 3.17E-11 | | |
| | | | | | | | | HK GWAS | 0.376 | 0.347 | 1.135 | 0.045 | 4.53E-03 | | |
| | | | | | | | | CC GWAS | 0.434 | 0.389 | 1.210 | 0.051 | 1.75E-04 | | |
| | | | | | | | | GZ GWAS | 0.382 | 0.344 | 1.164 | 0.060 | 1.10E-02 | | |
| | | | | | | | | KR IC | 0.414 | 0.376 | 1.170 | 0.042 | 2.78E-04 | | |
| | | | | | | | | BJ IC | 0.437 | 0.401 | 1.160 | 0.092 | 1.05E-01 | | |
| | | | | | | | | MC IC | 0.391 | 0.378 | 1.060 | 0.120 | 6.47E-01 | | |

| | | | | | | | | | | | | | | |
|---|---|---|---|---|---|---|---|---|---|---|---|---|---|---|
| | | | | | | | | EUR meta | - | - | 1.141 | 0.029 | 6.59E-06 | | |
| | | | | | | | | *EUR GWAS1* | *0.683* | *0.649* | *1.174* | *0.089* | *7.13E-02* | | |
| | | | | | | | | *EUR GWAS2* | *0.651* | *0.630* | *1.134* | *0.038* | *1.00E-03* | | |
| | | | | | | | | *EUR GWAS3* | *0.682* | *0.646* | *1.176* | *0.093* | *8.21E-02* | | |
| | | | | | | | | *SP GWAS* | - | - | *1.128* | *0.065* | *1.11E-01* | | |
| **#12** | **rs17321999** | **2** | **30479857** | **C** | **A** | **LBH** | **intronic** | **Trans meta** | **-** | **-** | **0.902** | **0.028** | **1.72E-04** | 0.181 | 27399966 |
| | | | | | | | | EAS meta | - | - | 0.939 | 0.041 | 1.24E-01 | | |
| | | | | | | | | *HK GWAS* | *0.124* | *0.136* | *0.904* | *0.066* | *1.25E-01* | | |
| | | | | | | | | *CC GWAS* | *0.189* | *0.204* | *0.920* | *0.063* | *1.90E-01* | | |
| | | | | | | | | *GZ GWAS* | *0.141* | *0.134* | *1.057* | *0.093* | *5.50E-01* | | |
| | | | | | | | | EUR meta | - | - | 0.872 | 0.037 | 2.34E-04 | | |
| | | | | | | | | *EUR GWAS1* | *0.109* | *0.166* | *0.603* | *0.120* | *2.40E-05* | | |
| | | | | | | | | *EUR GWAS2* | *0.193* | *0.208* | *0.902* | *0.046* | *2.40E-02* | | |
| | | | | | | | | *EUR GWAS3* | *0.103* | *0.114* | *0.895* | *0.143* | *4.38E-01* | | |
| | | | | | | | | SP GWAS | - | - | 0.932 | 0.090 | 2.55E-01 | | |
| **#13** | **rs13385731** | **2** | **33701890** | **T** | **C** | **RASGRP3** | **intronic** | **Trans meta** | **-** | **-** | **0.744** | **0.030** | **5.09E-23** | 6.16E-03 | 19838193 |
| | | | | | | | | EAS meta | - | - | 0.710 | 0.034 | 2.96E-23 | | |
| | | | | | | | | *HK GWAS* | *0.124* | *0.166* | *0.705* | *0.064* | *4.13E-08* | | |
| | | | | | | | | *CC GWAS* | *0.114* | *0.156* | *0.691* | *0.076* | *1.15E-06* | | |
| | | | | | | | | *GZ GWAS* | *0.120* | *0.165* | *0.678* | *0.084* | *3.32E-06* | | |
| | | | | | | | | *KR IC* | *0.102* | *0.127* | *0.780* | *0.073* | *3.22E-04* | | |
| | | | | | | | | *BJ IC* | *0.103* | *0.150* | *0.650* | *0.137* | *1.72E-03* | | |
| | | | | | | | | *MC IC* | *0.130* | *0.172* | *0.720* | *0.085* | *4.41E-02* | | |
| | | | | | | | | EUR meta | - | - | 0.859 | 0.060 | 1.12E-02 | | |
| | | | | | | | | *EUR GWAS1* | *0.048* | *0.049* | *1.022* | *0.188* | *9.06E-01* | | |

| | | | | | | | | | | | | | | | |
|---|---|---|---|---|---|---|---|---|---|---|---|---|---|---|---|
| | | | | | | | | *EUR GWAS2* | *0.060* | *0.068* | *0.885* | *0.075* | *1.03E-01* | | |
| | | | | | | | | *EUR GWAS3* | *0.036* | *0.045* | *0.782* | *0.231* | *2.88E-01* | | |
| | | | | | | | | *SP GWAS* | *-* | *-* | *0.733* | *0.137* | *3.48E-02* | | |
| **#14** | **rs6740462** | **2** | **65667272** | **C** | **A** | **SPRED2** | **intergenic** | **Trans meta** | **-** | **-** | **1.117** | **0.022** | **4.81E-07** | 0.022 | 26502338 |
| | | | | | | | | EAS meta | - | - | 1.066 | 0.030 | 3.38E-02 | | |
| | | | | | | | | *HK GWAS* | *0.862* | *0.848* | *1.112* | *0.062* | *8.52E-02* | | |
| | | | | | | | | *CC GWAS* | *0.800* | *0.797* | *1.013* | *0.063* | *8.33E-01* | | |
| | | | | | | | | *GZ GWAS* | *0.849* | *0.847* | *1.028* | *0.080* | *7.28E-01* | | |
| | | | | | | | | *KR IC* | *0.840* | *0.834* | *1.042* | *0.055* | *4.39E-01* | | |
| | | | | | | | | *BJ IC* | *0.794* | *0.765* | *1.176* | *0.108* | *1.31E-01* | | |
| | | | | | | | | *MC IC* | *0.849* | *0.816* | *1.266* | *0.145* | *1.39E-01* | | |
| | | | | | | | | EUR meta | - | - | 1.178 | 0.032 | 3.25E-07 | | |
| | | | | | | | | *EUR GWAS1* | *0.765* | *0.714* | *1.303* | *0.094* | *5.03E-03* | | |
| | | | | | | | | *EUR GWAS2* | *0.767* | *0.730* | *1.214* | *0.042* | *4.33E-06* | | |
| | | | | | | | | *EUR GWAS3* | *0.778* | *0.763* | *1.081* | *0.104* | *4.52E-01* | | |
| | | | | | | | | *SP GWAS* | *-* | *-* | *1.068* | *0.070* | *4.69E-01* | | |
| **#15** | **rs6705628** | **2** | **74208362** | **C** | **T** | **DGUOK-AS1** | **ncRNA_exonic** | **Trans meta** | **-** | **-** | **0.845** | **0.029** | **4.61E-09** | 0.284 | 23273568 |
| | | | | | | | | EAS meta | - | - | 0.851 | 0.029 | 4.71E-08 | | |
| | | | | | | | | *HK GWAS* | *0.123* | *0.149* | *0.804* | *0.065* | *8.51E-04* | | |
| | | | | | | | | *CC GWAS* | *0.183* | *0.212* | *0.839* | *0.063* | *5.65E-03* | | |
| | | | | | | | | *GZ GWAS* | *0.153* | *0.157* | *0.948* | *0.079* | *4.96E-01* | | |
| | | | | | | | | *KR IC* | *0.175* | *0.197* | *0.870* | *0.050* | *8.86E-03* | | |
| | | | | | | | | *BJ IC* | *0.199* | *0.222* | *0.870* | *0.110* | *2.09E-01* | | |
| | | | | | | | | *MC IC* | *0.113* | *0.169* | *0.620* | *0.283* | *6.35E-03* | | |
| | | | | | | | | EUR meta | - | - | 0.742 | 0.126 | 1.73E-02 | | |

| | | | | | | | | | | | | | | | |
|---|---|---|---|---|---|---|---|---|---|---|---|---|---|---|---|
| | | | | | | | | *EUR GWAS1* | *0.010* | *0.019* | *0.478* | *0.360* | *4.08E-02* | | |
| | | | | | | | | *EUR GWAS2* | *0.013* | *0.016* | *0.765* | *0.156* | *8.69E-02* | | |
| | | | | | | | | *EUR GWAS3* | *0.013* | *0.006* | *2.288* | *0.478* | *8.36E-02* | | |
| | | | | | | | | *SP GWAS* | *-* | *-* | *0.567* | *0.309* | *7.21E-02* | | |
| **#16** | **rs2322659** | **2** | **136555659** | **T** | **C** | **LCT** | **missense** | **EAS meta** | **-** | **-** | **1.106** | **0.023** | **7.38E-06** | QC fail in EUR | 29233832 |
| | | | | | | | | *HK GWAS* | *0.504* | *0.485* | *1.077* | *0.043* | *8.56E-02* | | |
| | | | | | | | | *CC GWAS* | *0.458* | *0.440* | *1.070* | *0.051* | *1.84E-01* | | |
| | | | | | | | | *GZ GWAS* | *0.520* | *0.475* | *1.195* | *0.057* | *1.71E-03* | | |
| | | | | | | | | *KR IC* | *0.532* | *0.505* | *1.111* | *0.043* | *1.28E-02* | | |
| | | | | | | | | *BJ IC* | *0.450* | *0.419* | *1.136* | *0.218* | *1.64E-01* | | |
| | | | | | | | | *MC IC* | *0.474* | *0.456* | *1.075* | *0.190* | *5.61E-01* | | |
| **#17** | **rs1990760** | **2** | **163124051** | **C** | **T** | **IFIH1** | **missense** | **Trans meta** | **-** | **-** | **1.110** | **0.020** | **1.82E-07** | 0.131 | 22046141 |
| | | | | | | | | EAS meta | - | - | 1.078 | 0.028 | 6.94E-03 | | |
| | | | | | | | | *HK GWAS* | *0.208* | *0.184* | *1.163* | *0.053* | *4.86E-03* | | |
| | | | | | | | | *CC GWAS* | *0.211* | *0.206* | *1.036* | *0.061* | *5.59E-01* | | |
| | | | | | | | | *GZ GWAS* | *0.201* | *0.178* | *1.151* | *0.073* | *5.43E-02* | | |
| | | | | | | | | *KR IC* | *0.184* | *0.182* | *1.010* | *0.053* | *8.90E-01* | | |
| | | | | | | | | *BJ IC* | *0.235* | *0.212* | *1.140* | *0.108* | *2.26E-01* | | |
| | | | | | | | | *MC IC* | *0.195* | *0.216* | *0.880* | *0.145* | *3.73E-01* | | |
| | | | | | | | | EUR meta | - | - | 1.145 | 0.029 | 2.44E-06 | | |
| | | | | | | | | *EUR GWAS1* | *0.625* | *0.595* | *1.122* | *0.085* | *1.78E-01* | | |
| | | | | | | | | *EUR GWAS2* | *0.638* | *0.610* | *1.091* | *0.038* | *2.14E-02* | | |
| | | | | | | | | *EUR GWAS3* | *0.634* | *0.562* | *1.360* | *0.091* | *7.43E-04* | | |
| | | | | | | | | *SP GWAS* | *-* | *-* | *1.218* | *0.063* | *7.18E-04* | | |
| **#18** | **rs11889341** | **2** | **191943742** | **C** | **T** | **STAT4** | **intronic** | **Trans meta** | **-** | **-** | **1.591** | **0.019** | **5.89E-137** | 0.022 | 26502338 |

| | | | | | | | | | | | | | | | |
|---|---|---|---|---|---|---|---|---|---|---|---|---|---|---|---|
| | | | | | | | | EAS meta | - | - | 1.543 | 0.023 | 3.79E-79 | | |
| | | | | | | | | *HK GWAS* | *0.443* | *0.344* | *1.525* | *0.045* | *4.01E-21* | | |
| | | | | | | | | *CC GWAS* | *0.424* | *0.337* | *1.465* | *0.052* | *2.79E-13* | | |
| | | | | | | | | *GZ GWAS* | *0.447* | *0.339* | *1.571* | *0.060* | *3.27E-14* | | |
| | | | | | | | | *KR IC* | *0.408* | *0.299* | *1.620* | *0.042* | *1.97E-27* | | |
| | | | | | | | | *BJ IC* | *0.409* | *0.334* | *1.380* | *0.094* | *5.30E-04* | | |
| | | | | | | | | *MC IC* | *0.453* | *0.331* | *1.670* | *0.122* | *2.51E-05* | | |
| | | | | | | | | EUR meta | - | - | 1.688 | 0.032 | 7.23E-61 | | |
| | | | | | | | | *EUR GWAS1* | *0.324* | *0.219* | *1.701* | *0.098* | *5.50E-08* | | |
| | | | | | | | | *EUR GWAS2* | *0.307* | *0.212* | *1.657* | *0.042* | *2.60E-33* | | |
| | | | | | | | | *EUR GWAS3* | *0.384* | *0.212* | *2.337* | *0.102* | *6.86E-17* | | |
| | | | | | | | | *SP GWAS* | - | - | *1.529* | *0.067* | *8.95E-10* | | |
| **#19** | **rs3768792** | **2** | **213871709** | **G** | **A** | **IKZF2** | **UTR3** | **Trans meta** | **-** | **-** | **0.869** | **0.026** | **8.36E-08** | 0.031 | 26502338 |
| | | | | | | | | EAS meta | - | - | 0.914 | 0.035 | 9.61E-03 | | |
| | | | | | | | | *HK GWAS* | *0.785* | *0.801* | *0.907* | *0.052* | *6.29E-02* | | |
| | | | | | | | | *CC GWAS* | *0.790* | *0.799* | *0.952* | *0.061* | *4.22E-01* | | |
| | | | | | | | | *GZ GWAS* | *0.787* | *0.808* | *0.873* | *0.073* | *6.26E-02* | | |
| | | | | | | | | EUR meta | - | - | 0.815 | 0.040 | 2.39E-07 | | |
| | | | | | | | | *EUR GWAS1* | *0.837* | *0.880* | *0.674* | *0.127* | *1.91E-03* | | |
| | | | | | | | | *EUR GWAS2* | *0.838* | *0.865* | *0.804* | *0.051* | *1.74E-05* | | |
| | | | | | | | | *EUR GWAS3* | *0.899* | *0.890* | *1.105* | *0.148* | *5.01E-01* | | |
| | | | | | | | | *SP GWAS* | - | - | *0.834* | *0.083* | *3.35E-02* | | |
| **#20** | **rs6445972** | **3** | **58321707** | **C** | **T** | **PXK** | **intronic** | **EUR meta** | **-** | **-** | **1.116** | **0.030** | **2.83E-04** | Mononorphic | 25620976 |
| | | | | | | | | EUR GWAS1 | 0.689 | 0.639 | 1.244 | 0.086 | 1.11E-02 | SNV in EAS | |
| | | | | | | | | EUR GWAS2 | 0.732 | 0.701 | 1.146 | 0.041 | 8.01E-04 | | |

| | | | | | | | | | | | | | | | |
|---|---|---|---|---|---|---|---|---|---|---|---|---|---|---|---|
| | | | | | | | | EUR GWAS3 | 0.661 | 0.658 | 1.014 | 0.094 | 8.81E-01 | | |
| | | | | | | | | SP GWAS | - | - | 1.027 | 0.065 | 4.32E-01 | | |
| **#21** | **rs6445975** | **3** | **58370177** | **G** | **T** | **PXK** | **intronic** | **Trans meta** | **-** | **-** | **0.914** | **0.020** | **9.31E-06** | 0.256 | 18204446 |
| | | | | | | | | EAS meta | - | - | 0.933 | 0.027 | 9.66E-03 | | |
| | | | | | | | | *HK GWAS* | *0.818* | *0.827* | *0.932* | *0.056* | *2.04E-01* | | |
| | | | | | | | | *CC GWAS* | *0.780* | *0.791* | *0.918* | *0.060* | *1.53E-01* | | |
| | | | | | | | | *GZ GWAS* | *0.801* | *0.807* | *0.969* | *0.071* | *6.56E-01* | | |
| | | | | | | | | *KR IC* | *0.762* | *0.770* | *0.952* | *0.046* | *3.73E-01* | | |
| | | | | | | | | *BJ IC* | *0.758* | *0.792* | *0.820* | *0.105* | *7.16E-02* | | |
| | | | | | | | | *MC IC* | *0.811* | *0.828* | *0.893* | *0.152* | *4.67E-01* | | |
| | | | | | | | | EUR meta | - | - | 0.890 | 0.031 | 1.61E-04 | | |
| | | | | | | | | *EUR GWAS1* | *0.763* | *0.783* | *0.902* | *0.098* | *2.93E-01* | | |
| | | | | | | | | *EUR GWAS2* | *0.674* | *0.706* | *0.881* | *0.039* | *1.07E-03* | | |
| | | | | | | | | *EUR GWAS3* | *0.773* | *0.774* | *0.992* | *0.106* | *9.38E-01* | | |
| | | | | | | | | *SP GWAS* | - | - | *0.875* | *0.070* | *5.04E-02* | | |
| **#22** | **rs9311676** | **3** | **58470351** | **C** | **T** | **LOC107986092** | **intronic** | **EUR meta** | **-** | **-** | **0.899** | **0.028** | **1.59E-04** | Rare in EAS | 26502338 |
| | | | | | | | | *EUR GWAS1* | *0.440* | *0.484* | *0.841* | *0.082* | *3.41E-02* | | |
| | | | | | | | | *EUR GWAS2* | *0.371* | *0.406* | *0.883* | *0.037* | *8.80E-04* | | |
| | | | | | | | | *EUR GWAS3* | *0.498* | *0.488* | *1.035* | *0.088* | *6.99E-01* | | |
| | | | | | | | | *SP GWAS* | - | - | *0.916* | *0.061* | *4.72E-02* | | |
| **#23** | **rs1131265** | **3** | **119222456** | **G** | **C** | **TIMMDC1;CD80** | **synonymous SNV** | **Trans meta** | **-** | **-** | **0.843** | **0.021** | **8.35E-16** | 0.121 | 28714469 |
| | | | | | | | | EAS meta | - | - | 0.827 | 0.025 | 1.37E-14 | | |
| | | | | | | | | *HK GWAS* | *0.312* | *0.342* | *0.878* | *0.046* | *4.81E-03* | | |
| | | | | | | | | *CC GWAS* | *0.281* | *0.324* | *0.821* | *0.055* | *3.27E-04* | | |
| | | | | | | | | *GZ GWAS* | *0.309* | *0.361* | *0.792* | *0.061* | *1.37E-04* | | |

| | | | | | | | | | | | | | | |
|---|---|---|---|---|---|---|---|---|---|---|---|---|---|---|
| | | | | | | | | *KR IC* | *0.263* | *0.308* | *0.800* | *0.049* | *2.94E-06* | | |
| | | | | | | | | *BJ IC* | *0.270* | *0.299* | *0.870* | *0.101* | *1.58E-01* | | |
| | | | | | | | | *MC IC* | *0.300* | *0.358* | *0.770* | *0.128* | *3.75E-02* | | |
| | | | | | | | | EUR meta | - | - | 0.891 | 0.041 | 4.89E-03 | | |
| | | | | | | | | *EUR GWAS1* | *0.133* | *0.170* | *0.752* | *0.114* | *1.24E-02* | | |
| | | | | | | | | *EUR GWAS2* | *0.174* | *0.189* | *0.903* | *0.047* | *3.09E-02* | | |
| | | | | | | | | *EUR GWAS3* | *0.169* | *0.170* | *0.986* | *0.120* | *9.04E-01* | | |
| **#24** | **rs564799** | **3** | **159728987** | **C** | **T** | **IL12A** | **ncRNA_intronic** | **Trans meta** | **-** | **-** | **0.864** | **0.022** | **3.19E-11** | 0.839 | 26502338 |
| | | | | | | | | EAS meta | - | - | 0.859 | 0.034 | 8.84E-06 | | |
| | | | | | | | | *HK GWAS* | *0.135* | *0.140* | *0.954* | *0.063* | *4.58E-01* | | |
| | | | | | | | | *CC GWAS* | *0.129* | *0.138* | *0.935* | *0.074* | *3.60E-01* | | |
| | | | | | | | | *GZ GWAS* | *0.113* | *0.158* | *0.688* | *0.083* | *6.78E-06* | | |
| | | | | | | | | *KR IC* | *0.091* | *0.107* | *0.840* | *0.073* | *1.49E-02* | | |
| | | | | | | | | *BJ IC* | *0.130* | *0.145* | *0.880* | *0.132* | *3.20E-01* | | |
| | | | | | | | | *MC IC* | *0.128* | *0.174* | *0.700* | *0.166* | *2.94E-02* | | |
| | | | | | | | | EUR meta | - | - | 0.867 | 0.029 | 8.02E-07 | | |
| | | | | | | | | *EUR GWAS1* | *0.331* | *0.345* | *0.939* | *0.090* | *4.82E-01* | | |
| | | | | | | | | *EUR GWAS2* | *0.395* | *0.426* | *0.877* | *0.037* | *3.90E-04* | | |
| | | | | | | | | *EUR GWAS3* | *0.303* | *0.341* | *0.842* | *0.097* | *7.47E-02* | | |
| | | | | | | | | *SP GWAS* | - | - | *0.814* | *0.065* | *1.70E-03* | | |
| **#25** | **rs10936599** | **3** | **169492101** | **C** | **T** | **MYNN** | **synonymous SNV** | **Trans meta** | **-** | **-** | **0.890** | **0.018** | **2.31E-10** | 0.676 | 28108556 |
| | | | | | | | | EAS meta | - | - | 0.895 | 0.022 | 4.50E-07 | | |
| | | | | | | | | *HK GWAS* | *0.521* | *0.540* | *0.925* | *0.043* | *7.14E-02* | | |
| | | | | | | | | *CC GWAS* | *0.541* | *0.556* | *0.954* | *0.050* | *3.41E-01* | | |
| | | | | | | | | *GZ GWAS* | *0.522* | *0.559* | *0.861* | *0.058* | *9.56E-03* | | |

| | | | | | | | | | | | | | | | |
|---|---|---|---|---|---|---|---|---|---|---|---|---|---|---|---|
| | | | | | | | | KR IC | 0.565 | 0.610 | 0.826 | 0.041 | 1.36E-05 | | |
| | | | | | | | | BJ IC | 0.508 | 0.540 | 0.885 | 0.091 | 1.64E-01 | | |
| | | | | | | | | MC IC | 0.544 | 0.516 | 1.124 | 0.122 | 3.40E-01 | | |
| | | | | | | | | EUR meta | - | - | 0.880 | 0.033 | 1.14E-04 | | |
| | | | | | | | | EUR GWAS1 | 0.194 | 0.221 | 0.857 | 0.102 | 1.31E-01 | | |
| | | | | | | | | EUR GWAS2 | 0.231 | 0.248 | 0.913 | 0.043 | 3.22E-02 | | |
| | | | | | | | | EUR GWAS3 | 0.196 | 0.249 | 0.737 | 0.107 | 4.53E-03 | | |
| | | | | | | | | SP GWAS | - | - | 0.869 | 0.075 | 8.31E-02 | | |
| **#26** | **rs6762714** | **3** | **188470238** | **C** | **T** | **LPP** | **intronic** | **Trans meta** | **-** | **-** | **1.100** | **0.023** | **3.43E-05** | 0.855 | 27399966 |
| | | | | | | | | EAS meta | - | - | 1.106 | 0.039 | 9.41E-03 | | |
| | | | | | | | | HK GWAS | 0.838 | 0.828 | 1.082 | 0.059 | 1.85E-01 | | |
| | | | | | | | | CC GWAS | 0.842 | 0.822 | 1.161 | 0.068 | 2.88E-02 | | |
| | | | | | | | | GZ GWAS | 0.831 | 0.822 | 1.080 | 0.079 | 3.31E-01 | | |
| | | | | | | | | EUR meta | - | - | 1.096 | 0.029 | 1.22E-03 | | |
| | | | | | | | | EUR GWAS1 | 0.441 | 0.385 | 1.268 | 0.087 | 6.26E-03 | | |
| | | | | | | | | EUR GWAS2 | 0.418 | 0.392 | 1.085 | 0.037 | 3.02E-02 | | |
| | | | | | | | | EUR GWAS3 | 0.374 | 0.358 | 1.075 | 0.092 | 4.33E-01 | | |
| | | | | | | | | SP GWAS | - | - | 1.060 | 0.061 | 5.25E-01 | | |
| **#27** | **rs4690229** | **4** | **970724** | **A** | **T** | **DGKQ;SLC26A1** | **intergenic** | **Trans meta** | **-** | **-** | **1.101** | **0.020** | **2.45E-06** | 0.652 | 28714469 |
| | | | | | | | | EAS meta | - | - | 1.090 | 0.030 | 4.14E-03 | | |
| | | | | | | | | HK GWAS | 0.342 | 0.333 | 1.044 | 0.046 | 3.58E-01 | | |
| | | | | | | | | CC GWAS | 0.407 | 0.372 | 1.174 | 0.051 | 1.69E-03 | | |
| | | | | | | | | GZ GWAS | 0.359 | 0.345 | 1.056 | 0.062 | 3.78E-01 | | |
| | | | | | | | | EUR meta | - | - | 1.110 | 0.028 | 1.65E-04 | | |
| | | | | | | | | EUR GWAS1 | 0.491 | 0.467 | 1.102 | 0.088 | 2.66E-01 | | |

| | | | | | | | | | | | | | | | |
|---|---|---|---|---|---|---|---|---|---|---|---|---|---|---|---|
| | | | | | | | | *EUR GWAS2* | *0.497* | *0.470* | *1.106* | *0.036* | *5.36E-03* | | |
| | | | | | | | | *EUR GWAS3* | *0.452* | *0.441* | *1.049* | *0.087* | *5.83E-01* | | |
| | | | | | | | | *SP GWAS* | *-* | *-* | *1.157* | *0.060* | *2.02E-02* | | |
| **#28** | **rs10028805** | **4** | **102737250** | **G** | **A** | **BANK1** | **intronic** | **Trans meta** | **-** | **-** | **0.854** | **0.019** | **2.67E-17** | 0.985 | 26502338 |
| | | | | | | | | EAS meta | - | - | 0.854 | 0.024 | 8.41E-11 | | |
| | | | | | | | | *HK GWAS* | *0.346* | *0.390* | *0.823* | *0.045* | *1.49E-05* | | |
| | | | | | | | | *CC GWAS* | *0.292* | *0.324* | *0.854* | *0.055* | *3.91E-03* | | |
| | | | | | | | | *GZ GWAS* | *0.353* | *0.371* | *0.937* | *0.060* | *2.81E-01* | | |
| | | | | | | | | *KR IC* | *0.235* | *0.280* | *0.790* | *0.049* | *2.18E-06* | | |
| | | | | | | | | *BJ IC* | *0.275* | *0.300* | *0.880* | *0.100* | *2.08E-01* | | |
| | | | | | | | | *MC IC* | *0.394* | *0.358* | *1.160* | *0.124* | *2.15E-01* | | |
| | | | | | | | | EUR meta | - | - | 0.854 | 0.029 | 5.85E-08 | | |
| | | | | | | | | *EUR GWAS1* | *0.355* | *0.370* | *0.937* | *0.086* | *4.48E-01* | | |
| | | | | | | | | *EUR GWAS2* | *0.331* | *0.370* | *0.858* | *0.039* | *7.13E-05* | | |
| | | | | | | | | *EUR GWAS3* | *0.353* | *0.397* | *0.834* | *0.091* | *4.57E-02* | | |
| | | | | | | | | *SP GWAS* | *-* | *-* | *0.811* | *0.063* | *1.99E-03* | | |
| **#29** | **rs7726159** | **5** | **1282319** | **C** | **A** | **TERT** | **intron** | **Trans meta** | **-** | **-** | **1.134** | **0.019** | **6.04E-11** | 0.150 | 26808113 |
| | | | | | | | | EAS meta | - | - | 1.156 | 0.023 | 4.72E-10 | | |
| | | | | | | | | *HK GWAS* | *0.418* | *0.386* | *1.155* | *0.046* | *1.68E-03* | | |
| | | | | | | | | *CC GWAS* | *0.434* | *0.399* | *1.182* | *0.054* | *1.92E-03* | | |
| | | | | | | | | *GZ GWAS* | *0.416* | *0.385* | *1.134* | *0.058* | *3.10E-02* | | |
| | | | | | | | | *KR IC* | *0.397* | *0.364* | *1.150* | *0.043* | *1.27E-03* | | |
| | | | | | | | | *BJ IC* | *0.442* | *0.420* | *1.090* | *0.094* | *3.26E-01* | | |
| | | | | | | | | *MC IC* | *0.479* | *0.416* | *1.290* | *0.119* | *3.37E-02* | | |
| | | | | | | | | EUR meta | - | - | 1.089 | 0.034 | 1.36E-02 | | |

| | | | | | | | | | | | | | | | |
|---|---|---|---|---|---|---|---|---|---|---|---|---|---|---|---|
| | | | | | | | | *EUR GWAS1* | *0.414* | *0.390* | *1.120* | *0.089* | *2.05E-01* | | |
| | | | | | | | | *EUR GWAS2* | *0.355* | *0.336* | *1.092* | *0.040* | *2.96E-02* | | |
| | | | | | | | | *EUR GWAS3* | *0.401* | *0.394* | *1.035* | *0.098* | *7.29E-01* | | |
| **#30** | **rs7726414** | **5** | **133431834** | **C** | **T** | **VDAC1;TCF7** | **intergenic** | **Trans meta** | **-** | **-** | **1.317** | **0.034** | **3.17E-16** | 0.638 | (1)26502338 |
| | | | | | | | | EAS meta | - | - | 1.302 | 0.042 | 2.26E-10 | | (2)26808113 |
| | | | | | | | | *HK GWAS* | *0.099* | *0.089* | *1.122* | *0.075* | *1.24E-01* | | |
| | | | | | | | | *CC GWAS* | *0.086* | *0.060* | *1.433* | *0.092* | *9.03E-05* | | |
| | | | | | | | | *GZ GWAS* | *0.107* | *0.086* | *1.306* | *0.103* | *9.76E-03* | | |
| | | | | | | | | *KR IC* | *0.068* | *0.052* | *1.330* | *0.088* | *1.25E-03* | | |
| | | | | | | | | *BJ IC* | *0.085* | *0.048* | *1.850* | *0.187* | *9.56E-04* | | |
| | | | | | | | | *MC IC* | *0.130* | *0.096* | *1.410* | *0.191* | *6.58E-02* | | |
| | | | | | | | | EUR meta | - | - | 1.346 | 0.058 | 2.39E-07 | | |
| | | | | | | | | *EUR GWAS1* | *0.096* | *0.085* | *1.131* | *0.146* | *4.00E-01* | | |
| | | | | | | | | *EUR GWAS2* | *0.055* | *0.037* | *1.454* | *0.086* | *1.26E-05* | | |
| | | | | | | | | *EUR GWAS3* | *0.089* | *0.072* | *1.247* | *0.162* | *1.74E-01* | | |
| | | | | | | | | *SP GWAS* | *-* | *-* | *1.356* | *0.111* | *2.11E-02* | | |
| **#31** | **rs10036748** | **5** | **150458146** | **C** | **T** | **TNIP1** | **intronic** | **Trans meta** | **-** | **-** | **1.278** | **0.021** | **5.65E-33** | 0.222 | 19838193 |
| | | | | | | | | EAS meta | - | - | 1.250 | 0.027 | 3.22E-16 | | |
| | | | | | | | | *HK GWAS* | *0.772* | *0.740* | *1.192* | *0.051* | *5.42E-04* | | |
| | | | | | | | | *CC GWAS* | *0.792* | *0.758* | *1.216* | *0.061* | *1.23E-03* | | |
| | | | | | | | | *GZ GWAS* | *0.798* | *0.752* | *1.296* | *0.069* | *1.57E-04* | | |
| | | | | | | | | *KR IC* | *0.820* | *0.781* | *1.282* | *0.056* | *7.01E-06* | | |
| | | | | | | | | *BJ IC* | *0.813* | *0.750* | *1.449* | *0.112* | *7.56E-04* | | |
| | | | | | | | | *MC IC* | *0.782* | *0.751* | *1.190* | *0.138* | *2.07E-01* | | |
| | | | | | | | | EUR meta | - | - | 1.315 | 0.031 | 1.09E-18 | | |

| | | | | | | | | | | | | | | |
|---|---|---|---|---|---|---|---|---|---|---|---|---|---|---|
| | | | | | | | | *EUR GWAS1* | *0.313* | *0.264* | *1.250* | *0.093* | *1.69E-02* | | |
| | | | | | | | | *EUR GWAS2* | *0.295* | *0.239* | *1.321* | *0.041* | *1.24E-11* | | |
| | | | | | | | | *EUR GWAS3* | *0.377* | *0.297* | *1.429* | *0.094* | *1.48E-04* | | |
| | | | | | | | | *SP GWAS* | *-* | *-* | *1.277* | *0.068* | *7.64E-04* | | |
| **#32** | **rs2431697** | **5** | **159879978** | **T** | **C** | **PTTG1;MIR146A** | **intergenic** | **Trans meta** | **-** | **-** | **0.784** | **0.022** | **5.98E-29** | 0.784 | (2)25890262, |
| | | | | | | | | EAS meta | - | - | 0.779 | 0.034 | 1.57E-13 | | (1)26502338 |
| | | | | | | | | *HK GWAS* | *0.086* | *0.108* | *0.789* | *0.075* | *1.52E-03* | | |
| | | | | | | | | *CC GWAS* | *0.140* | *0.184* | *0.728* | *0.070* | *5.89E-06* | | |
| | | | | | | | | *GZ GWAS* | *0.096* | *0.107* | *0.850* | *0.096* | *9.08E-02* | | |
| | | | | | | | | *KR IC* | *0.165* | *0.198* | *0.800* | *0.060* | *8.86E-05* | | |
| | | | | | | | | *BJ IC* | *0.145* | *0.209* | *0.640* | *0.120* | *2.00E-04* | | |
| | | | | | | | | *MC IC* | *0.116* | *0.110* | *1.060* | *0.187* | *7.47E-01* | | |
| | | | | | | | | EUR meta | - | - | 0.788 | 0.028 | 5.16E-17 | | |
| | | | | | | | | *EUR GWAS1* | *0.414* | *0.440* | *0.898* | *0.083* | *1.98E-01* | | |
| | | | | | | | | *EUR GWAS2* | *0.376* | *0.439* | *0.771* | *0.038* | *4.82E-12* | | |
| | | | | | | | | *EUR GWAS3* | *0.366* | *0.462* | *0.680* | *0.090* | *1.77E-05* | | |
| | | | | | | | | *SP GWAS* | *-* | *-* | *0.834* | *0.061* | *7.93E-03* | | |
| **#33** | **rs9270984** | **6** | **32573991** | **T** | **G** | **HLA region** | **intergenic** | **Trans meta** | **-** | **-** | **0.634** | **0.027** | **6.51E-66** | 8E-13 | (1)23273568, |
| | | | | | | | | EAS meta | - | - | 0.543 | 0.034 | 4.37E-71 | | (2)20169177 |
| | | | | | | | | *HK GWAS* | *0.711* | *0.811* | *0.558* | *0.052* | *1.91E-29* | | |
| | | | | | | | | *CC GWAS* | *0.752* | *0.847* | *0.517* | *0.062* | *4.39E-26* | | |
| | | | | | | | | *GZ GWAS* | *0.665* | *0.780* | *0.547* | *0.067* | *3.42E-19* | | |
| | | | | | | | | EUR meta | - | - | 0.801 | 0.042 | 1.55E-07 | | |
| | | | | | | | | *EUR GWAS1* | *0.837* | *0.904* | *0.523* | *0.137* | *2.25E-06* | | |
| | | | | | | | | *EUR GWAS2* | *0.808* | *0.835* | *0.843* | *0.047* | *2.95E-04* | | |

| | | | | | | | | | | | | | | | |
|---|---|---|---|---|---|---|---|---|---|---|---|---|---|---|---|
| | | | | | | | | EUR GWAS3 | 0.859 | 0.883 | 0.798 | 0.132 | 8.74E-02 | | |
| #34 | rs3734266 | 6 | 34823187 | T | C | UHRF1BP1 | intronic | Trans meta | - | - | 1.289 | 0.025 | 9.40E-24 | 0.083 | (1)23273568 |
| | | | | | | | | EAS meta | - | - | 1.339 | 0.033 | 1.96E-18 | | (2)26502338 |
| | | | | | | | | HK GWAS | 0.166 | 0.125 | 1.378 | 0.060 | 1.02E-07 | | |
| | | | | | | | | CC GWAS | 0.178 | 0.127 | 1.481 | 0.067 | 4.00E-09 | | |
| | | | | | | | | GZ GWAS | 0.134 | 0.107 | 1.296 | 0.089 | 3.54E-03 | | |
| | | | | | | | | KR IC | 0.100 | 0.087 | 1.160 | 0.074 | 3.81E-02 | | |
| | | | | | | | | BJ IC | 0.164 | 0.120 | 1.450 | 0.130 | 4.60E-03 | | |
| | | | | | | | | MC IC | 0.133 | 0.117 | 1.160 | 0.180 | 4.08E-01 | | |
| | | | | | | | | EUR meta | - | - | 1.225 | 0.039 | 1.80E-07 | | |
| | | | | | | | | EUR GWAS1 | 0.206 | 0.174 | 1.230 | 0.106 | 4.96E-02 | | |
| | | | | | | | | EUR GWAS2 | 0.136 | 0.112 | 1.267 | 0.055 | 1.63E-05 | | |
| | | | | | | | | EUR GWAS3 | 0.174 | 0.157 | 1.127 | 0.119 | 3.16E-01 | | |
| | | | | | | | | SP GWAS | - | - | 1.184 | 0.077 | 2.67E-02 | | |
| #35 | rs597325 | 6 | 91002494 | A | G | BACH2 | intronic | Trans meta | - | - | 1.087 | 0.018 | 3.26E-06 | 0.653 | 27399966 |
| | | | | | | | | EAS meta | - | - | 1.094 | 0.023 | 8.56E-05 | | |
| | | | | | | | | HK GWAS | 0.502 | 0.484 | 1.078 | 0.043 | 8.41E-02 | | |
| | | | | | | | | CC GWAS | 0.508 | 0.486 | 1.099 | 0.050 | 5.95E-02 | | |
| | | | | | | | | GZ GWAS | 0.522 | 0.514 | 1.031 | 0.061 | 6.17E-01 | | |
| | | | | | | | | KR IC | 0.453 | 0.425 | 1.120 | 0.044 | 8.26E-03 | | |
| | | | | | | | | BJ IC | 0.514 | 0.505 | 1.040 | 0.051 | 6.83E-01 | | |
| | | | | | | | | MC IC | 0.530 | 0.453 | 1.360 | 0.120 | 9.14E-03 | | |
| | | | | | | | | EUR meta | - | - | 1.076 | 0.029 | 1.13E-02 | | |
| | | | | | | | | EUR GWAS1 | 0.655 | 0.665 | 0.955 | 0.088 | 5.97E-01 | | |
| | | | | | | | | EUR GWAS2 | 0.629 | 0.606 | 1.121 | 0.038 | 2.39E-03 | | |

| | | | | | | | | | | | | | | | |
|---|---|---|---|---|---|---|---|---|---|---|---|---|---|---|---|
| | | | | | | | | EUR GWAS3 | 0.675 | 0.644 | 1.141 | 0.093 | 1.57E-01 | | |
| | | | | | | | | SP GWAS | - | - | 0.991 | 0.063 | 6.04E-01 | | |
| **#36** | **rs548234** | **6** | **106568034** | **C** | **T** | **PRDM1;ATG5** | intergenic | **Trans meta** | **-** | **-** | **0.814** | **0.019** | **2.39E-28** | 0.275 | 19838193 |
| | | | | | | | | EAS meta | - | - | 0.801 | 0.024 | 2.39E-20 | | |
| | | | | | | | | *HK GWAS* | *0.676* | *0.713* | *0.835* | *0.047* | *1.26E-04* | | |
| | | | | | | | | *CC GWAS* | *0.681* | *0.739* | *0.749* | *0.055* | *1.40E-07* | | |
| | | | | | | | | *GZ GWAS* | *0.680* | *0.718* | *0.834* | *0.062* | *3.72E-03* | | |
| | | | | | | | | *KR IC* | *0.671* | *0.718* | *0.800* | *0.043* | *1.59E-06* | | |
| | | | | | | | | *BJ IC* | *0.690* | *0.752* | *0.730* | *0.101* | *1.92E-03* | | |
| | | | | | | | | *MC IC* | *0.671* | *0.710* | *0.833* | *0.127* | *1.54E-01* | | |
| | | | | | | | | EUR meta | - | - | 0.835 | 0.029 | 8.29E-10 | | |
| | | | | | | | | *EUR GWAS1* | *0.645* | *0.704* | *0.756* | *0.091* | *2.04E-03* | | |
| | | | | | | | | *EUR GWAS2* | *0.629* | *0.668* | *0.833* | *0.038* | *1.67E-06* | | |
| | | | | | | | | *EUR GWAS3* | *0.693* | *0.730* | *0.823* | *0.099* | *4.85E-02* | | |
| | | | | | | | | *SP GWAS* | *-* | *-* | *0.888* | *0.064* | *4.15E-02* | | |
| **#37** | **rs2327832** | **6** | **137973068** | **A** | **G** | **OLIG3** | **intergenic** | **EUR meta** | **-** | **-** | **1.177** | **0.034** | **1.14E-06** | QC fail in EAS | 28714469 |
| | | | | | | | | *EUR GWAS1* | *0.211* | *0.214* | *0.979* | *0.102* | *8.36E-01* | | |
| | | | | | | | | *EUR GWAS2* | *0.247* | *0.216* | *1.196* | *0.043* | *3.48E-05* | | |
| | | | | | | | | *EUR GWAS3* | *0.204* | *0.165* | *1.320* | *0.115* | *1.63E-02* | | |
| | | | | | | | | *SP GWAS* | *-* | *-* | *1.181* | *0.074* | *1.47E-02* | | |
| **#38** | **rs2230926** | **6** | **138196066** | **T** | **G** | **TNFAIP3** | missense | **Trans meta** | **-** | **-** | **1.881** | **0.039** | **7.34E-59** | 0.033 | 19838193 |
| | | | | | | | | EAS meta | - | - | 2.001 | 0.049 | 5.76E-46 | | |
| | | | | | | | | *HK GWAS* | *0.055* | *0.024* | *2.371* | *0.112* | *1.57E-14* | | |
| | | | | | | | | *CC GWAS* | *0.074* | *0.045* | *1.695* | *0.101* | *1.55E-07* | | |
| | | | | | | | | *GZ GWAS* | *0.057* | *0.029* | *2.025* | *0.156* | *6.24E-06* | | |

| | | | | | | | | | | | | | | | |
|---|---|---|---|---|---|---|---|---|---|---|---|---|---|---|---|
| | | | | | | | | *KR IC* | *0.100* | *0.055* | *1.930* | *0.078* | *5.22E-17* | | |
| | | | | | | | | *BJ IC* | *0.106* | *0.043* | *2.670* | *0.188* | *7.82E-08* | | |
| | | | | | | | | *MC IC* | *0.060* | *0.030* | *2.080* | *0.303* | *1.39E-02* | | |
| | | | | | | | | EUR meta | - | - | 1.683 | 0.065 | 1.48E-15 | | |
| | | | | | | | | *EUR GWAS1* | *0.068* | *0.047* | *1.518* | *0.188* | *2.66E-02* | | |
| | | | | | | | | *EUR GWAS2* | *0.054* | *0.034* | *1.737* | *0.089* | *4.50E-10* | | |
| | | | | | | | | *EUR GWAS3* | *0.071* | *0.031* | *2.331* | *0.205* | *3.62E-05* | | |
| | | | | | | | | *SP GWAS* | - | - | *1.433* | *0.134* | *5.99E-02* | | |
| **#39** | **rs849142** | **7** | **28185891** | **T** | **C** | **JAZF1** | **intronic** | **Trans meta** | **-** | **-** | **0.923** | **0.027** | **3.40E-03** | 0.578 | (1)19838195, |
| | | | | | | | | EAS meta | - | - | 0.854 | 0.143 | 2.68E-01 | | (2)26502338 |
| | | | | | | | | *HK GWAS* | *0.006* | *0.007* | *0.862* | *0.279* | *5.95E-01* | | |
| | | | | | | | | *CC GWAS* | *0.011* | *0.013* | *0.893* | *0.237* | *6.34E-01* | | |
| | | | | | | | | *GZ GWAS* | *0.006* | *0.007* | *0.887* | *0.363* | *7.41E-01* | | |
| | | | | | | | | EUR meta | - | - | 0.926 | 0.028 | 5.63E-03 | | |
| | | | | | | | | *EUR GWAS1* | *0.510* | *0.537* | *0.903* | *0.084* | *2.24E-01* | | |
| | | | | | | | | *EUR GWAS2* | *0.486* | *0.503* | *0.953* | *0.037* | *1.94E-01* | | |
| | | | | | | | | *EUR GWAS3* | *0.445* | *0.501* | *0.790* | *0.090* | *9.04E-03* | | |
| | | | | | | | | *SP GWAS* | - | - | *0.929* | *0.060* | *3.85E-01* | | |
| **#40** | **rs2366293** | **7** | **50227828** | **G** | **C** | **C7orf72;IKZF1** | **intergenic** | **Trans meta** | **-** | **-** | **0.873** | **0.034** | **5.45E-05** | 0.190 | 22046141 |
| | | | | | | | | EAS meta | - | - | 0.970 | 0.087 | 7.23E-01 | | |
| | | | | | | | | *HK GWAS* | *0.968* | *0.970* | *0.907* | *0.128* | *4.48E-01* | | |
| | | | | | | | | *CC GWAS* | *0.973* | *0.976* | *0.861* | *0.160* | *3.50E-01* | | |
| | | | | | | | | *GZ GWAS* | *0.971* | *0.965* | *1.274* | *0.178* | *1.73E-01* | | |
| | | | | | | | | EUR meta | - | - | 0.857 | 0.037 | 2.36E-05 | | |
| | | | | | | | | *EUR GWAS1* | *0.797* | *0.801* | *0.963* | *0.104* | *7.19E-01* | | |

| | | | | | | | | | | | | | | | |
|---|---|---|---|---|---|---|---|---|---|---|---|---|---|---|---|
| | | | | | | | | EUR GWAS2 | 0.834 | 0.855 | 0.819 | 0.051 | 8.67E-05 | | |
| | | | | | | | | EUR GWAS3 | 0.790 | 0.783 | 1.049 | 0.110 | 6.65E-01 | | |
| | | | | | | | | SP GWAS | - | - | 0.809 | 0.074 | 1.87E-02 | | |
| **#41** | **rs4917014** | **7** | **50305863** | **T** | **G** | **C7orf72;IKZF1** | **intergenic** | **Trans meta** | **-** | **-** | **0.798** | **0.019** | **2.17E-31** | 4E-04 | (1)19838193, |
| | | | | | | | | EAS meta | - | - | 0.753 | 0.025 | 5.18E-29 | | (2)26502338 |
| | | | | | | | | HK GWAS | 0.239 | 0.291 | 0.757 | 0.050 | 2.23E-08 | | |
| | | | | | | | | CC GWAS | 0.256 | 0.318 | 0.729 | 0.057 | 2.77E-08 | | |
| | | | | | | | | GZ GWAS | 0.238 | 0.270 | 0.829 | 0.067 | 4.83E-03 | | |
| | | | | | | | | KR IC | 0.325 | 0.398 | 0.730 | 0.047 | 2.22E-12 | | |
| | | | | | | | | BJ IC | 0.263 | 0.325 | 0.740 | 0.100 | 2.87E-03 | | |
| | | | | | | | | MC IC | 0.239 | 0.284 | 0.790 | 0.135 | 8.07E-02 | | |
| | | | | | | | | EUR meta | - | - | 0.865 | 0.030 | 1.34E-06 | | |
| | | | | | | | | EUR GWAS1 | 0.299 | 0.367 | 0.741 | 0.088 | 6.44E-04 | | |
| | | | | | | | | EUR GWAS2 | 0.295 | 0.321 | 0.884 | 0.040 | 1.85E-03 | | |
| | | | | | | | | EUR GWAS3 | 0.357 | 0.351 | 1.023 | 0.093 | 8.05E-01 | | |
| | | | | | | | | SP GWAS | - | - | 0.821 | 0.065 | 8.40E-03 | | |
| **#42** | **rs13238909** | **7** | **67076373** | **G** | **A** | **ST3AGL4** | **intergenic** | **Trans meta** | **-** | **-** | **0.912** | **0.033** | **5.08E-03** | 0.956 | 29625966 |
| | | | | | | | | EAS meta | - | - | 0.915 | 0.069 | 1.97E-01 | | |
| | | | | | | | | HK GWAS | 0.034 | 0.043 | 0.808 | 0.115 | 6.27E-02 | | |
| | | | | | | | | CC GWAS | 0.063 | 0.066 | 0.972 | 0.101 | 7.81E-01 | | |
| | | | | | | | | GZ GWAS | 0.036 | 0.034 | 1.006 | 0.162 | 9.72E-01 | | |
| | | | | | | | | EUR meta | - | - | 0.911 | 0.037 | 1.29E-02 | | |
| | | | | | | | | EUR GWAS1 | 0.134 | 0.170 | 0.774 | 0.113 | 2.30E-02 | | |
| | | | | | | | | EUR GWAS2 | 0.161 | 0.178 | 0.885 | 0.049 | 1.25E-02 | | |
| | | | | | | | | EUR GWAS3 | 0.167 | 0.162 | 1.039 | 0.117 | 7.41E-01 | | |

| | | | | | | | | | | | | | | | |
|---|---|---|---|---|---|---|---|---|---|---|---|---|---|---|---|
| | | | | | | | | *SP GWAS* | - | - | *1.013* | *0.083* | *6.55E-01* | | |
| **#43** | **rs73135369** | **7** | **73940978** | **T** | **C** | **GTF2IRD1** | **intronic** | **Trans meta** | **-** | **-** | **1.350** | **0.045** | **2.46E-11** | 0.104 | 27399966 |
| | | | | | | | | EAS meta | - | - | 1.400 | 0.050 | 2.15E-11 | | |
| | | | | | | | | *HK GWAS* | *0.095* | *0.068* | *1.472* | *0.080* | *1.18E-06* | | |
| | | | | | | | | *CC GWAS* | *0.115* | *0.089* | *1.365* | *0.082* | *1.36E-04* | | |
| | | | | | | | | *GZ GWAS* | *0.095* | *0.072* | *1.336* | *0.107* | *6.71E-03* | | |
| | | | | | | | | EUR meta | - | - | 1.167 | 0.100 | 1.24E-01 | | |
| | | | | | | | | *EUR GWAS1* | *0.021* | *0.017* | *1.250* | *0.305* | *4.65E-01* | | |
| | | | | | | | | *EUR GWAS2* | *0.028* | *0.021* | *1.200* | *0.116* | *1.15E-01* | | |
| | | | | | | | | *EUR GWAS3* | *0.027* | *0.028* | *0.952* | *0.269* | *8.53E-01* | | |
| **#44** | **rs117026326** | **7** | **74126034** | **C** | **T** | **GTF2I** | **intronic** | **EAS meta** | **-** | **-** | **2.994** | **0.102** | **7.35E-27** | QC fail in EUR | 26808113; |
| | | | | | | | | *GZ GWAS* | *0.198* | *0.081* | *2.994* | *0.102* | *7.35E-27* | | 28135245 |
| **#45** | **rs11773745** | **7** | **75171438** | **A** | **G** | **HIP1** | **intronic** | Trans meta | - | - | 1.115 | 0.023 | 1.41E-06 | 3.77E-06 | 19838193 |
| | | | | | | | | **EAS meta** | **-** | **-** | **1.239** | **0.032** | **2.52E-11** | | 26663300 |
| | | | | | | | | *HK GWAS* | *0.716* | *0.676* | *1.209* | *0.048* | *7.60E-05* | | |
| | | | | | | | | *CC GWAS* | *0.743* | *0.696* | *1.277* | *0.058* | *2.38E-05* | | |
| | | | | | | | | *GZ GWAS* | *0.725* | *0.683* | *1.249* | *0.065* | *6.77E-04* | | |
| | | | | | | | | EUR meta | - | - | 1.006 | 0.032 | 8.55E-01 | | |
| | | | | | | | | *EUR GWAS1* | *0.219* | *0.244* | *0.870* | *0.098* | *1.55E-01* | | |
| | | | | | | | | *EUR GWAS2* | *0.274* | *0.266* | *1.036* | *0.041* | *3.91E-01* | | |
| | | | | | | | | *EUR GWAS3* | *0.253* | *0.261* | *0.965* | *0.082* | *6.63E-01* | | |
| | | | | | | | | *Spain GWAS* | - | - | *0.983* | *0.073* | *9.26E-01* | | |
| **#46** | **rs729302** | **7** | **128568960** | **A** | **C** | **KCP;IRF5** | **intergenic** | **Trans meta** | **-** | **-** | **0.774** | **0.019** | **2.80E-41** | 0.385 | (1)23273568 |
| | | | | | | | | EAS meta | - | - | 0.763 | 0.024 | 1.74E-28 | | (2)23053960 |
| | | | | | | | | *HK GWAS* | *0.262* | *0.337* | *0.705* | *0.048* | *3.08E-13* | | |

| | | | | | | | | | | | | | | | |
|---|---|---|---|---|---|---|---|---|---|---|---|---|---|---|---|
| | | | | | | | | *CC GWAS* | *0.294* | *0.344* | *0.792* | *0.054* | *1.81E-05* | | |
| | | | | | | | | *GZ GWAS* | *0.284* | *0.331* | *0.794* | *0.062* | *1.99E-04* | | |
| | | | | | | | | *KR IC* | *0.265* | *0.326* | *0.750* | *0.046* | *6.14E-10* | | |
| | | | | | | | | *BJ IC* | *0.321* | *0.355* | *0.860* | *0.097* | *1.16E-01* | | |
| | | | | | | | | *MC IC* | *0.319* | *0.348* | *0.870* | *0.129* | *2.86E-01* | | |
| | | | | | | | | EUR meta | - | - | 0.790 | 0.031 | 1.36E-14 | | |
| | | | | | | | | *EUR GWAS1* | *0.260* | *0.298* | *0.840* | *0.092* | *5.70E-02* | | |
| | | | | | | | | *EUR GWAS2* | *0.270* | *0.326* | *0.768* | *0.040* | *5.17E-11* | | |
| | | | | | | | | *EUR GWAS3* | *0.280* | *0.328* | *0.797* | *0.097* | *1.90E-02* | | |
| | | | | | | | | *SP GWAS* | *-* | *-* | *0.823* | *0.067* | *8.82E-03* | | |
| **#47** | **rs4728142** | **7** | **128573967** | **G** | **A** | **KCP;IRF5** | **intergenic** | **Trans meta** | **-** | **-** | **1.451** | **0.021** | **4.64E-71** | 7.89E-03 | (1)19838193, |
| | | | | | | | | EAS meta | - | - | 1.562 | 0.032 | 2.55E-45 | | (2)24871463 |
| | | | | | | | | *HK GWAS* | *0.182* | *0.125* | *1.567* | *0.060* | *7.35E-14* | | |
| | | | | | | | | *CC GWAS* | *0.178* | *0.134* | *1.423* | *0.068* | *2.35E-07* | | |
| | | | | | | | | *GZ GWAS* | *0.176* | *0.117* | *1.638* | *0.086* | *9.13E-09* | | |
| | | | | | | | | *KR IC* | *0.180* | *0.117* | *1.660* | *0.061* | *5.60E-18* | | |
| | | | | | | | | *BJ IC* | *0.196* | *0.121* | *1.770* | *0.127* | *4.87E-06* | | |
| | | | | | | | | *MC IC* | *0.167* | *0.148* | *1.150* | *0.162* | *3.88E-01* | | |
| | | | | | | | | EUR meta | - | - | 1.370 | 0.028 | 1.27E-29 | | |
| | | | | | | | | *EUR GWAS1* | *0.558* | *0.502* | *1.241* | *0.085* | *1.08E-02* | | |
| | | | | | | | | *EUR GWAS2* | *0.524* | *0.440* | *1.405* | *0.037* | *1.81E-20* | | |
| | | | | | | | | *EUR GWAS3* | *0.506* | *0.457* | *1.213* | *0.088* | *2.80E-02* | | |
| | | | | | | | | *SP GWAS* | *-* | *-* | *1.429* | *0.061* | *4.03E-09* | | |
| **#48** | **rs12531711** | **7** | **128617466** | **A** | **G** | **IRF5;TNPO3** | **intron** | **EUR meta** | **-** | **-** | **1.718** | **0.040** | **7.38E-41** | Mononorphic | 23053960 |
| | | | | | | | | *EUR GWAS1* | *0.147* | *0.111* | *1.375* | *0.133* | *1.66E-02* | SNV in EAS | |

| | | | | | | | | | | | | | | | |
|---|---|---|---|---|---|---|---|---|---|---|---|---|---|---|---|
| | | | | | | | | *EUR GWAS2* | *0.185* | *0.109* | *1.775* | *0.051* | *3.12E-29* | | |
| | | | | | | | | *EUR GWAS3* | *0.149* | *0.094* | *1.692* | *0.136* | *1.04E-04* | | |
| | | | | | | | | *SP GWAS* | - | - | *1.735* | *0.092* | *1.19E-08* | | |
| **#49** | **rs2428** | **8** | **8641145** | **C** | **T** | **MFHAS1** | **UTR3** | **Trans meta** | **-** | **-** | ***1.073*** | ***0.026*** | ***7.04E-03*** | 0.103 | 29625966 |
| | | | | | | | | EAS meta | - | - | *1.017* | *0.042* | *6.92E-01* | | |
| | | | | | | | | *HK GWAS* | *0.871* | *0.862* | *1.070* | *0.063* | *2.84E-01* | | |
| | | | | | | | | *CC GWAS* | *0.860* | *0.871* | *0.942* | *0.075* | *4.25E-01* | | |
| | | | | | | | | *GZ GWAS* | *0.875* | *0.872* | *1.024* | *0.088* | *7.87E-01* | | |
| | | | | | | | | EUR meta | - | - | *1.110* | *0.034* | *1.80E-03* | | |
| | | | | | | | | *EUR GWAS1* | *0.494* | *0.459* | *1.137* | *0.082* | *1.18E-01* | | |
| | | | | | | | | *EUR GWAS2* | *0.512* | *0.476* | *1.025* | *0.054* | *6.44E-01* | | |
| | | | | | | | | *EUR GWAS3* | *0.539* | *0.502* | *1.164* | *0.089* | *8.82E-02* | | |
| | | | | | | | | *SP GWAS* | - | - | *1.185* | *0.061* | *1.09E-02* | | |
| **#50** | **rs12680762** | **8** | **11332026** | **C** | **T** | **FAM167A** | **intron** | **EUR meta** | **-** | **-** | **1.216** | **0.035** | **3.50E-08** | QC fail in EAS | 23053960 |
| | | | | | | | | *EUR GWAS1* | *0.263* | *0.244* | *1.110* | *0.099* | *2.94E-01* | | |
| | | | | | | | | *EUR GWAS2* | *0.320* | *0.272* | *1.193* | *0.050* | *3.80E-04* | | |
| | | | | | | | | *EUR GWAS3* | *0.290* | *0.240* | *1.344* | *0.106* | *5.14E-03* | | |
| | | | | | | | | *SP GWAS* | - | - | *1.264* | *0.071* | *2.90E-03* | | |
| **#51** | **rs2736340** | **8** | **11343973** | **C** | **T** | **FAM167A;BLK** | **intergenic** | **Trans meta** | **-** | **-** | **1.363** | **0.021** | **2.01E-50** | 0.065 | 19838195 |
| | | | | | | | | EAS meta | - | - | 1.403 | 0.026 | 1.29E-38 | | |
| | | | | | | | | *HK GWAS* | *0.780* | *0.712* | *1.431* | *0.051* | *2.82E-12* | | |
| | | | | | | | | *CC GWAS* | *0.797* | *0.733* | *1.427* | *0.060* | *3.79E-09* | | |
| | | | | | | | | *GZ GWAS* | *0.774* | *0.715* | *1.359* | *0.066* | *2.98E-06* | | |
| | | | | | | | | *KR IC* | *0.787* | *0.726* | *1.389* | *0.047* | *3.93E-11* | | |
| | | | | | | | | *BJ IC* | *0.784* | *0.725* | *1.370* | *0.107* | *2.58E-03* | | |

| | | | | | | | | | | | | | | | |
|---|---|---|---|---|---|---|---|---|---|---|---|---|---|---|---|
| | | | | | | | | *MC IC* | *0.774* | *0.704* | *1.449* | *0.141* | *7.14E-03* | | |
| | | | | | | | | EUR meta | - | - | 1.296 | 0.034 | 3.36E-14 | | |
| | | | | | | | | *EUR GWAS1* | *0.272* | *0.258* | *1.064* | *0.093* | *5.01E-01* | | |
| | | | | | | | | *EUR GWAS2* | *0.314* | *0.258* | *1.275* | *0.047* | *3.02E-07* | | |
| | | | | | | | | *EUR GWAS3* | *0.310* | *0.233* | *1.514* | *0.102* | *4.72E-05* | | |
| | | | | | | | | *SP GWAS* | - | - | *1.397* | *0.071* | *2.04E-05* | | |
| **#52** | **rs7097397** | **10** | **50025396** | **G** | **A** | **WDFY4** | **missense** | **Trans meta** | **-** | **-** | **0.804** | **0.018** | **1.82E-32** | 0.069 | 20169177 |
| | | | | | | | | EAS meta | - | - | 0.783 | 0.024 | 4.79E-25 | | |
| | | | | | | | | *HK GWAS* | *0.649* | *0.706* | *0.750* | *0.048* | *1.55E-09* | | |
| | | | | | | | | *CC GWAS* | *0.617* | *0.667* | *0.791* | *0.054* | *1.31E-05* | | |
| | | | | | | | | *GZ GWAS* | *0.640* | *0.676* | *0.852* | *0.062* | *1.04E-02* | | |
| | | | | | | | | *KR IC* | *0.579* | *0.638* | *0.781* | *0.042* | *1.25E-08* | | |
| | | | | | | | | *BJ IC* | *0.599* | *0.681* | *0.699* | *0.094* | *1.37E-04* | | |
| | | | | | | | | *MC IC* | *0.650* | *0.677* | *0.885* | *0.124* | *3.36E-01* | | |
| | | | | | | | | EUR meta | - | - | 0.838 | 0.029 | 1.07E-09 | | |
| | | | | | | | | *EUR GWAS1* | *0.391* | *0.456* | *0.759* | *0.087* | *1.55E-03* | | |
| | | | | | | | | *EUR GWAS2* | *0.320* | *0.363* | *0.841* | *0.039* | *6.95E-06* | | |
| | | | | | | | | *EUR GWAS3* | *0.385* | *0.465* | *0.707* | *0.092* | *1.72E-04* | | |
| | | | | | | | | *SP GWAS* | - | - | *0.940* | *0.061* | *1.74E-01* | | |
| **#53** | **rs4948496** | **10** | **63805617** | **T** | **C** | **ARID5B** | **intronic** | **Trans meta** | **-** | **-** | **1.139** | **0.020** | **1.29E-10** | 0.871 | (1)23273568, |
| | | | | | | | | EAS meta | - | - | 1.143 | 0.030 | 6.00E-06 | | (2)26502338 |
| | | | | | | | | *HK GWAS* | *0.663* | *0.623* | *1.183* | *0.045* | *1.94E-04* | | |
| | | | | | | | | *CC GWAS* | *0.656* | *0.625* | *1.133* | *0.052* | *1.56E-02* | | |
| | | | | | | | | *GZ GWAS* | *0.652* | *0.633* | *1.089* | *0.060* | *1.56E-01* | | |
| | | | | | | | | EUR meta | - | - | 1.136 | 0.028 | 4.92E-06 | | |

| | | | | | | | | | | | | | | | |
|---|---|---|---|---|---|---|---|---|---|---|---|---|---|---|---|
| | | | | | | | | *EUR GWAS1* | *0.536* | *0.499* | *1.176* | *0.084* | *5.42E-02* | | |
| | | | | | | | | *EUR GWAS2* | *0.481* | *0.449* | *1.132* | *0.037* | *6.81E-04* | | |
| | | | | | | | | *EUR GWAS3* | *0.569* | *0.546* | *1.088* | *0.089* | *3.46E-01* | | |
| | | | | | | | | *SP GWAS* | *-* | *-* | *1.147* | *0.060* | *7.81E-02* | | |
| **#54** | **rs12802200** | **11** | **566936** | **C** | **A** | **MIR210HG** | **ncRNA_exonic** | **Trans meta** | **-** | **-** | **0.820** | **0.033** | **1.59E-09** | 0.966 | 26502338 |
| | | | | | | | | EAS meta | - | - | 0.818 | 0.080 | 1.23E-02 | | |
| | | | | | | | | *HK GWAS* | *0.024* | *0.028* | *0.841* | *0.140* | *2.16E-01* | | |
| | | | | | | | | *CC GWAS* | *0.018* | *0.019* | *0.888* | *0.202* | *5.55E-01* | | |
| | | | | | | | | *GZ GWAS* | *0.017* | *0.024* | *0.667* | *0.202* | *4.53E-02* | | |
| | | | | | | | | *KR IC* | *0.017* | *0.018* | *0.930* | *0.167* | *6.71E-01* | | |
| | | | | | | | | *BJ IC* | *0.019* | *0.027* | *0.700* | *0.304* | *2.41E-01* | | |
| | | | | | | | | *MC IC* | *0.023* | *0.033* | *0.680* | *0.365* | *2.91E-01* | | |
| | | | | | | | | EUR meta | - | - | 0.821 | 0.036 | 3.97E-08 | | |
| | | | | | | | | *EUR GWAS1* | *0.172* | *0.202* | *0.815* | *0.106* | *5.41E-02* | | |
| | | | | | | | | *EUR GWAS2* | *0.165* | *0.196* | *0.819* | *0.048* | *2.93E-05* | | |
| | | | | | | | | *EUR GWAS3* | *0.166* | *0.208* | *0.758* | *0.116* | *1.67E-02* | | |
| | | | | | | | | *SP GWAS* | *-* | *-* | *0.857* | *0.077* | *6.86E-02* | | |
| **#55** | **rs2732552** | **11** | **35084592** | **T** | **C** | **PDHX** | **intergenic** | **Trans meta** | **-** | **-** | **1.174** | **0.019** | **1.13E-16** | 0.565 | (1)26502338 |
| | | | | | | | | EAS meta | - | - | 1.162 | 0.027 | 1.74E-08 | | (2)21194677 |
| | | | | | | | | *HK GWAS* | *0.788* | *0.771* | *1.100* | *0.052* | *6.82E-02* | | |
| | | | | | | | | *CC GWAS* | *0.782* | *0.754* | *1.172* | *0.061* | *8.83E-03* | | |
| | | | | | | | | *GZ GWAS* | *0.789* | *0.760* | *1.192* | *0.069* | *1.12E-02* | | |
| | | | | | | | | *KR IC* | *0.731* | *0.690* | *1.220* | *0.047* | *2.52E-05* | | |
| | | | | | | | | *BJ IC* | *0.775* | *0.749* | *1.149* | *0.106* | *1.75E-01* | | |
| | | | | | | | | *MC IC* | *0.784* | *0.790* | *0.962* | *0.144* | *8.04E-01* | | |

| | | | | | | | | | | | | | | | |
|---|---|---|---|---|---|---|---|---|---|---|---|---|---|---|---|
| | | | | | | | | EUR meta | - | - | 1.188 | 0.028 | 1.02E-09 | | |
| | | | | | | | | *EUR GWAS1* | *0.597* | *0.548* | *1.233* | *0.085* | *1.34E-02* | | |
| | | | | | | | | *EUR GWAS2* | *0.588* | *0.554* | *1.161* | *0.037* | *5.19E-05* | | |
| | | | | | | | | *EUR GWAS3* | *0.618* | *0.542* | *1.378* | *0.092* | *4.73E-04* | | |
| | | | | | | | | *SP GWAS* | - | - | *1.160* | *0.061* | *3.31E-02* | | |
| **#56** | **rs1061502** | **11** | **614318** | **T** | **C** | **IRF7** | **missense** | ***Trans meta*** | **-** | **-** | ***0.835*** | ***0.030*** | **1.11E-09** | 0.866 | 21360504 |
| | | | | | | | | *EAS meta* | - | - | *0.795* | *0.101* | *2.25E-02* | | |
| | | | | | | | | *HK GWAS* | *0.023* | *0.027* | *0.837* | *0.142* | *2.09E-01* | | |
| | | | | | | | | *CC GWAS* | *0.016* | *0.018* | *0.883* | *0.200* | *5.34E-01* | | |
| | | | | | | | | *GZ GWAS* | *0.016* | *0.024* | *0.637* | *0.206* | *2.82E-02* | | |
| | | | | | | | | EUR meta | - | - | *0.834* | *0.032* | *1.10E-08* | | |
| | | | | | | | | *EUR GWAS1* | *0.232* | *0.277* | *0.799* | *0.094* | *1.74E-02* | | |
| | | | | | | | | *EUR GWAS2* | *0.229* | *0.269* | *0.815* | *0.042* | *1.31E-06* | | |
| | | | | | | | | *EUR GWAS3* | *0.258* | *0.282* | *0.885* | *0.099* | *2.18E-01* | | |
| | | | | | | | | *SP GWAS* | - | - | *0.879* | *0.069* | *9.48E-02* | | |
| **#57** | **rs2009453** | **11** | **65399528** | **C** | **T** | **PCNX3;RELA** | **intronic** | **Trans meta** | **-** | **-** | **0.885** | **0.018** | **4.22E-12** | 0.034 | 26808113 |
| | | | | | | | | EAS meta | - | - | 0.859 | 0.023 | 1.96E-11 | | |
| | | | | | | | | *HK GWAS* | *0.469* | *0.485* | *0.944* | *0.044* | *1.84E-01* | | |
| | | | | | | | | *CC GWAS* | *0.409* | *0.452* | *0.828* | *0.051* | *2.53E-04* | | |
| | | | | | | | | *GZ GWAS* | *0.455* | *0.503* | *0.829* | *0.058* | *1.19E-03* | | |
| | | | | | | | | *KR IC* | *0.400* | *0.448* | *0.820* | *0.042* | *5.52E-06* | | |
| | | | | | | | | *BJ IC* | *0.416* | *0.448* | *0.880* | *0.090* | *1.53E-01* | | |
| | | | | | | | | *MC IC* | *0.430* | *0.470* | *0.850* | *0.117* | *1.66E-01* | | |
| | | | | | | | | EUR meta | - | - | 0.927 | 0.028 | 6.23E-03 | | |
| | | | | | | | | *EUR GWAS1* | *0.506* | *0.527* | *0.932* | *0.083* | *3.95E-01* | | |

| | | | | | | | | | | | | | | | |
|---|---|---|---|---|---|---|---|---|---|---|---|---|---|---|---|
| | | | | | | | | *EUR GWAS2* | *0.535* | *0.551* | *0.921* | *0.037* | *2.37E-02* | | |
| | | | | | | | | *EUR GWAS3* | *0.592* | *0.571* | *1.094* | *0.090* | *3.16E-01* | | |
| | | | | | | | | *SP GWAS* | *-* | *-* | *0.873* | *0.060* | *3.42E-02* | | |
| #58 | rs1308020 | 11 | 65497558 | G | A | RNASEH2C | intergenic | **Trans meta** | **-** | **-** | **0.874** | **0.022** | **1.28E-09** | 7.87E-03 | 28108556 |
| | | | | | | | | EAS meta | - | - | 0.818 | 0.033 | 1.83E-09 | | |
| | | | | | | | | *HK GWAS* | *0.227* | *0.260* | *0.843* | *0.051* | *7.29E-04* | | |
| | | | | | | | | *CC GWAS* | *0.213* | *0.250* | *0.814* | *0.060* | *5.98E-04* | | |
| | | | | | | | | *GZ GWAS* | *0.224* | *0.271* | *0.783* | *0.066* | *2.09E-04* | | |
| | | | | | | | | EUR meta | - | - | 0.921 | 0.030 | 5.33E-03 | | |
| | | | | | | | | *EUR GWAS1* | *0.356* | *0.373* | *0.935* | *0.087* | *4.45E-01* | | |
| | | | | | | | | *EUR GWAS2* | *0.319* | *0.343* | *0.908* | *0.039* | *1.41E-02* | | |
| | | | | | | | | *EUR GWAS3* | *0.342* | *0.353* | *0.954* | *0.093* | *6.17E-01* | | |
| | | | | | | | | *SP GWAS* | *-* | *-* | *0.932* | *0.063* | *2.02E-01* | | |
| #59 | rs494003 | 11 | 65542298 | G | A | AP5B1 | UTR3 | **Trans meta** | **-** | **-** | **1.172** | **0.027** | **8.02E-09** | 0.177 | 27399966 |
| | | | | | | | | EAS meta | - | - | 1.113 | 0.047 | 2.08E-02 | | |
| | | | | | | | | *HK GWAS* | *0.109* | *0.105* | *1.040* | *0.070* | *5.76E-01* | | |
| | | | | | | | | *CC GWAS* | *0.106* | *0.093* | *1.167* | *0.082* | *5.80E-02* | | |
| | | | | | | | | *GZ GWAS* | *0.114* | *0.098* | *1.185* | *0.096* | *7.69E-02* | | |
| | | | | | | | | EUR meta | - | - | 1.204 | 0.034 | 4.92E-08 | | |
| | | | | | | | | *EUR GWAS1* | *0.218* | *0.226* | *0.943* | *0.100* | *5.57E-01* | | |
| | | | | | | | | *EUR GWAS2* | *0.225* | *0.195* | *1.192* | *0.044* | *6.94E-05* | | |
| | | | | | | | | *EUR GWAS3* | *0.171* | *0.153* | *1.145* | *0.121* | *2.63E-01* | | |
| | | | | | | | | *SP GWAS* | *-* | *-* | *1.437* | *0.074* | *5.27E-07* | | |
| #60 | rs10750836 | 11 | 68815523 | C | T | TPCN2 | upstream | Trans meta | - | - | 0.907 | 0.028 | 5.46E-04 | 0.066 | 29233832 |
| | | | | | | | | **EAS meta** | **-** | **-** | **0.897** | **0.029** | **1.67E-04** | | |

| | | | | | | | | | | | | | | | |
|---|---|---|---|---|---|---|---|---|---|---|---|---|---|---|---|
| | | | | | | | | *HK GWAS* | *0.503* | *0.516* | *0.952* | *0.044* | *2.65E-01* | | |
| | | | | | | | | *CC GWAS* | *0.554* | *0.589* | *0.863* | *0.053* | *5.25E-03* | | |
| | | | | | | | | *GZ GWAS* | *0.494* | *0.534* | *0.847* | *0.057* | *3.49E-03* | | |
| | | | | | | | | EUR meta | - | - | 1.157 | 0.136 | 2.83E-01 | | |
| | | | | | | | | *EUR GWAS1* | *0.977* | *0.971* | *1.324* | *0.253* | *2.68E-01* | | |
| | | | | | | | | *EUR GWAS2* | *0.990* | *0.990* | *1.179* | *0.183* | *3.68E-01* | | |
| | | | | | | | | *EUR GWAS3* | *0.982* | *0.984* | *0.849* | *0.341* | *6.32E-01* | | |
| **#61** | **rs11235604** | **11** | **72533536** | **C** | **T** | **ATG16L2** | **missense** | **EAS meta** | **-** | **-** | **0.778** | **0.052** | **1.26E-06** | Mononorphic | 28108556 |
| | | | | | | | | HK GWAS | 0.076 | 0.097 | 0.774 | 0.079 | 1.24E-03 | SNV in EUR | |
| | | | | | | | | CC GWAS | 0.073 | 0.096 | 0.741 | 0.094 | 1.42E-03 | | |
| | | | | | | | | GZ GWAS | 0.087 | 0.102 | 0.830 | 0.099 | 5.84E-02 | | |
| **#62** | **rs11603023** | **11** | **118486067** | **T** | **C** | **PHLDB1** | **intronic** | **Trans meta** | **-** | **-** | **0.910** | **0.019** | **4.28E-07** | 0.107 | 24001599 |
| | | | | | | | | EAS meta | - | - | 0.886 | 0.025 | 1.27E-06 | | |
| | | | | | | | | *HK GWAS* | *0.649* | *0.674* | *0.899* | *0.046* | *2.02E-02* | | |
| | | | | | | | | *CC GWAS* | *0.744* | *0.772* | *0.870* | *0.057* | *1.52E-02* | | |
| | | | | | | | | *GZ GWAS* | *0.662* | *0.688* | *0.864* | *0.062* | *1.92E-02* | | |
| | | | | | | | | *KR IC* | *0.758* | *0.774* | *0.917* | *0.049* | *8.15E-02* | | |
| | | | | | | | | *BJ IC* | *0.760* | *0.800* | *0.794* | *0.109* | *3.22E-02* | | |
| | | | | | | | | *MC IC* | *0.675* | *0.702* | *0.885* | *0.131* | *3.30E-01* | | |
| | | | | | | | | EUR meta | - | - | 0.941 | 0.028 | 3.02E-02 | | |
| | | | | | | | | *EUR GWAS1* | *0.534* | *0.584* | *0.812* | *0.085* | *1.45E-02* | | |
| | | | | | | | | *EUR GWAS2* | *0.577* | *0.567* | *1.016* | *0.037* | *6.72E-01* | | |
| | | | | | | | | *EUR GWAS3* | *0.542* | *0.580* | *0.855* | *0.088* | *7.57E-02* | | |
| | | | | | | | | *SP GWAS* | - | - | *0.862* | *0.061* | *2.70E-02* | | |
| **#63** | **rs4639966** | **11** | **118573519** | **T** | **C** | **TREH;DDX6** | **intergenic** | Trans meta | - | - | 1.106 | 0.019 | 1.20E-07 | 7.76E-03 | 19838193 |

| | | | | | | | | | | | | | | | |
|---|---|---|---|---|---|---|---|---|---|---|---|---|---|---|---|
| | | | | | | | | **EAS meta** | **-** | **-** | **1.147** | **0.024** | **5.20E-09** | | |
| | | | | | | | | *HK GWAS* | *0.450* | *0.421* | *1.116* | *0.044* | *1.21E-02* | | |
| | | | | | | | | *CC GWAS* | *0.340* | *0.303* | *1.162* | *0.053* | *4.69E-03* | | |
| | | | | | | | | *GZ GWAS* | *0.440* | *0.401* | *1.200* | *0.058* | *1.71E-03* | | |
| | | | | | | | | *KR IC* | *0.298* | *0.272* | *1.140* | *0.047* | *5.58E-03* | | |
| | | | | | | | | *BJ IC* | *0.304* | *0.282* | *1.110* | *0.100* | *2.81E-01* | | |
| | | | | | | | | *MC IC* | *0.439* | *0.395* | *1.200* | *0.121* | *1.33E-01* | | |
| | | | | | | | | EUR meta | - | - | 1.032 | 0.032 | 3.23E-01 | | |
| | | | | | | | | *EUR GWAS1* | *0.293* | *0.258* | *1.198* | *0.094* | *5.57E-02* | | |
| | | | | | | | | *EUR GWAS2* | *0.244* | *0.248* | *0.983* | *0.043* | *6.84E-01* | | |
| | | | | | | | | *EUR GWAS3* | *0.278* | *0.249* | *1.170* | *0.100* | *1.18E-01* | | |
| | | | | | | | | *SP GWAS* | *-* | *-* | *1.022* | *0.069* | *8.52E-01* | | |
| **#64** | **rs1128334** | **11** | **128328959** | **C** | **T** | **ETS1** | **UTR3** | **Trans meta** | **-** | **-** | **1.355** | **0.021** | **6.41E-49** | 0.204 | 20169177 |
| | | | | | | | | EAS meta | - | - | 1.373 | 0.023 | 1.70E-42 | | |
| | | | | | | | | *HK GWAS* | *0.411* | *0.351* | *1.293* | *0.045* | *8.58E-09* | | |
| | | | | | | | | *CC GWAS* | *0.399* | *0.341* | *1.274* | *0.051* | *2.20E-06* | | |
| | | | | | | | | *GZ GWAS* | *0.435* | *0.353* | *1.414* | *0.060* | *8.54E-09* | | |
| | | | | | | | | *KR IC* | *0.454* | *0.366* | *1.440* | *0.044* | *3.38E-17* | | |
| | | | | | | | | *BJ IC* | *0.423* | *0.300* | *1.710* | *0.095* | *1.29E-08* | | |
| | | | | | | | | *MC IC* | *0.425* | *0.343* | *1.420* | *0.124* | *4.39E-03* | | |
| | | | | | | | | EUR meta | - | - | 1.287 | 0.045 | 2.52E-08 | | |
| | | | | | | | | *EUR GWAS1* | *0.084* | *0.080* | *1.041* | *0.151* | *7.89E-01* | | |
| | | | | | | | | *EUR GWAS2* | *0.126* | *0.096* | *1.370* | *0.057* | *4.22E-08* | | |
| | | | | | | | | *EUR GWAS3* | *0.084* | *0.087* | *0.959* | *0.161* | *7.96E-01* | | |
| | | | | | | | | *SP GWAS* | *-* | *-* | *1.311* | *0.100* | *2.98E-03* | | |

| | | | | | | | | | | | | | | | |
|---|---|---|---|---|---|---|---|---|---|---|---|---|---|---|---|
| #65 | rs7941765 | 11 | 128499000 | T | C | ETS1 | intergenic | Trans meta | - | - | 1.134 | 0.022 | 6.37E-09 | 0.908 | 26502338 |
| | | | | | | LOC101929538 | | EAS meta | - | - | 1.130 | 0.035 | 4.20E-04 | | |
| | | | | | | | | HK GWAS | 0.787 | 0.759 | 1.178 | 0.052 | 1.76E-03 | | |
| | | | | | | | | CC GWAS | 0.795 | 0.778 | 1.122 | 0.063 | 6.58E-02 | | |
| | | | | | | | | GZ GWAS | 0.786 | 0.776 | 1.061 | 0.069 | 3.86E-01 | | |
| | | | | | | | | EUR meta | - | - | 1.136 | 0.028 | 3.95E-06 | | |
| | | | | | | | | EUR GWAS1 | 0.492 | 0.438 | 1.221 | 0.082 | 1.46E-02 | | |
| | | | | | | | | EUR GWAS2 | 0.552 | 0.515 | 1.145 | 0.036 | 1.90E-04 | | |
| | | | | | | | | EUR GWAS3 | 0.507 | 0.491 | 1.067 | 0.089 | 4.66E-01 | | |
| | | | | | | | | SP GWAS | - | - | 1.099 | 0.060 | 1.80E-01 | | |
| #66 | rs10845606 | 12 | 12834894 | C | A | GPR19 | intronic | Trans meta | - | - | 0.878 | 0.023 | 1.90E-08 | 8.73E-03 | 23273568 |
| | | | | | | | | EAS meta | - | - | 0.829 | 0.032 | 3.80E-09 | | |
| | | | | | | | | HK GWAS | 0.285 | 0.328 | 0.811 | 0.048 | 1.27E-05 | | |
| | | | | | | | | CC GWAS | 0.242 | 0.282 | 0.811 | 0.058 | 2.83E-04 | | |
| | | | | | | | | GZ GWAS | 0.277 | 0.303 | 0.884 | 0.063 | 4.98E-02 | | |
| | | | | | | | | EUR meta | - | - | 0.937 | 0.034 | 5.27E-02 | | |
| | | | | | | | | EUR GWAS1 | 0.212 | 0.228 | 0.919 | 0.101 | 4.04E-01 | | |
| | | | | | | | | EUR GWAS2 | 0.212 | 0.224 | 0.927 | 0.044 | 8.96E-02 | | |
| | | | | | | | | EUR GWAS3 | 0.177 | 0.212 | 0.796 | 0.115 | 4.71E-02 | | |
| | | | | | | | | SP GWAS | - | - | 1.035 | 0.072 | 8.90E-01 | | |
| #67 | rs10774625 | 12 | 111910219 | A | G | ATXN2;SH2B3 | intron | EUR meta | - | - | 0.827 | 0.028 | 7.28E-12 | Rare in EAS | 26502338 |
| | | | | | | | | EUR GWAS1 | 0.496 | 0.545 | 0.824 | 0.084 | 2.17E-02 | | |
| | | | | | | | | EUR GWAS2 | 0.451 | 0.503 | 0.798 | 0.036 | 6.35E-10 | | |
| | | | | | | | | EUR GWAS3 | 0.446 | 0.475 | 0.892 | 0.089 | 1.98E-01 | | |
| | | | | | | | | SP GWAS | - | - | 0.879 | 0.060 | 7.40E-02 | | |

| | | | | | | | | | | | | | | | |
|---|---|---|---|---|---|---|---|---|---|---|---|---|---|---|---|
| **#68** | **rs1385374** | **12** | **129300694** | **C** | **T** | **SLC15A4** | **intronic** | **Trans meta** | **-** | **-** | **1.286** | **0.023** | **2.62E-27** | 0.812 | 19838193 |
| | | | | | | | | EAS meta | - | - | 1.290 | 0.027 | 1.35E-20 | | |
| | | | | | | | | *HK GWAS* | *0.207* | *0.165* | *1.339* | *0.056* | *1.46E-07* | | |
| | | | | | | | | *CC GWAS* | *0.264* | *0.203* | *1.435* | *0.058* | *4.67E-10* | | |
| | | | | | | | | *GZ GWAS* | *0.212* | *0.184* | *1.179* | *0.073* | *2.43E-02* | | |
| | | | | | | | | *KR IC* | *0.257* | *0.216* | *1.250* | *0.050* | *5.94E-06* | | |
| | | | | | | | | *BJ IC* | *0.255* | *0.221* | *1.210* | *0.103* | *7.67E-02* | | |
| | | | | | | | | *MC IC* | *0.205* | *0.196* | *1.060* | *0.149* | *6.90E-01* | | |
| | | | | | | | | EUR meta | - | - | 1.275 | 0.044 | 3.05E-08 | | |
| | | | | | | | | *EUR GWAS1* | *0.139* | *0.092* | *1.652* | *0.140* | *3.36E-04* | | |
| | | | | | | | | *EUR GWAS2* | *0.122* | *0.107* | *1.235* | *0.057* | *2.12E-04* | | |
| | | | | | | | | *EUR GWAS3* | *0.127* | *0.099* | *1.327* | *0.142* | *4.56E-02* | | |
| | | | | | | | | *SP GWAS* | *-* | *-* | *1.214* | *0.095* | *1.25E-01* | | |
| **#69** | **rs7329174** | **13** | **41558110** | **A** | **G** | **ELF1** | **intronic** | **Trans meta** | **-** | **-** | **1.189** | **0.033** | **1.40E-07** | 0.554 | 21044949 |
| | | | | | | | | EAS meta | - | - | 1.192 | 0.033 | 1.17E-07 | | |
| | | | | | | | | *HK GWAS* | *0.265* | *0.217* | *1.296* | *0.050* | *2.23E-07* | | |
| | | | | | | | | *CC GWAS* | *0.229* | *0.205* | *1.147* | *0.060* | *2.13E-02* | | |
| | | | | | | | | *GZ GWAS* | *0.267* | *0.252* | *1.081* | *0.065* | *2.36E-01* | | |
| | | | | | | | | EUR meta | - | - | 1.010 | 0.278 | 9.72E-01 | | |
| | | | | | | | | *EUR GWAS1* | *0.006* | *0.006* | *1.096* | *0.570* | *8.72E-01* | | |
| | | | | | | | | *EUR GWAS3* | *0.004* | *0.006* | *0.642* | *0.673* | *5.10E-01* | | |
| | | | | | | | | *SP GWAS* | *-* | *-* | *1.114* | *0.362* | *9.63E-01* | | |
| **#70** | **rs12874404** | **13** | **108993494** | **A** | **G** | **TNFSF13B** | **intergenic** | **EUR meta** | **-** | **-** | **1.293** | **0.061** | **2.24E-05** | Mononorphic | 28445677 |
| | | | | | | | | *EUR GWAS1* | *0.092* | *0.073* | *1.327* | *0.165* | *8.58E-02* | SNV in EAS | |
| | | | | | | | | *EUR GWAS2* | *0.079* | *0.065* | *1.296* | *0.073* | *3.63E-04* | | |

| | | | | | | | | | | | | | | | |
|---|---|---|---|---|---|---|---|---|---|---|---|---|---|---|---|
| | | | | | | | | *EUR GWAS3* | *0.125* | *0.104* | *1.252* | *0.146* | *1.22E-01* | | |
| **#71** | **rs4902562** | **14** | **68731458** | **A** | **G** | **RAD51B** | **intronic** | Trans meta | - | - | 0.924 | 0.021 | 1.52E-04 | 0.045 | 26502338 |
| | | | | | | | | EAS meta | - | - | 0.967 | 0.031 | 2.70E-01 | | |
| | | | | | | | | *HK GWAS* | *0.313* | *0.323* | *0.961* | *0.047* | *3.98E-01* | | |
| | | | | | | | | *CC GWAS* | *0.392* | *0.400* | *0.973* | *0.051* | *5.88E-01* | | |
| | | | | | | | | *GZ GWAS* | *0.341* | *0.346* | *0.968* | *0.067* | *6.25E-01* | | |
| | | | | | | | | **EUR meta** | **-** | **-** | **0.889** | **0.028** | **3.50E-05** | | |
| | | | | | | | | *EUR GWAS1* | *0.630* | *0.665* | *0.856* | *0.088* | *7.66E-02* | | |
| | | | | | | | | *EUR GWAS2* | *0.542* | *0.569* | *0.926* | *0.037* | *3.70E-02* | | |
| | | | | | | | | *EUR GWAS3* | *0.590* | *0.631* | *0.836* | *0.092* | *5.15E-02* | | |
| | | | | | | | | *SP GWAS* | - | - | *0.830* | *0.062* | *4.24E-03* | | |
| **#72** | **rs2841280** | **14** | **105393556** | **G** | **C** | **PLD4** | **nonsynonymous** | Trans meta | - | - | 1.125 | 0.020 | 5.56E-09 | 5.148E-04 | 29233832 |
| | | | | | | | | **EAS meta** | **-** | **-** | 1.210 | **0.029** | **6.12E-11** | | |
| | | | | | | | | *HK GWAS* | 0.620 | 0.575 | 1.208 | 0.044 | 1.84E-05 | | |
| | | | | | | | | *CC GWAS* | 0.644 | 0.600 | 1.211 | 0.052 | 2.55E-04 | | |
| | | | | | | | | *GZ GWAS* | 0.621 | 0.573 | 1.215 | 0.058 | 8.80E-04 | | |
| | | | | | | | | EUR meta | - | - | 1.052 | 0.028 | 7.10E-02 | | |
| | | | | | | | | *EUR GWAS1* | 0.438 | 0.438 | 1.000 | 0.083 | 9.97E-01 | | |
| | | | | | | | | *EUR GWAS2* | 0.485 | 0.468 | 1.039 | 0.037 | 2.99E-01 | | |
| | | | | | | | | *EUR GWAS3* | 0.393 | 0.361 | 1.163 | 0.093 | 1.05E-01 | | |
| | | | | | | | | *Spain GWAS* | - | - | 1.073 | 0.061 | 2.21E-01 | | |
| **#73** | **rs117518546** | **14** | **106204113** | **C** | **T** | **IGHG1** | **nonsynonymous** | *CC GWAS* | 0.433 | 0.407 | 1.065 | 0.059 | 2.86E-01 | Monomorphic in EUR | 30287618 |
| **#74** | **rs8035957** | **15** | **38838264** | **T** | **C** | **RASGRP1** | **intronic** | **Trans meta** | **-** | **-** | **1.123** | **0.018** | **3.23E-10** | 0.379 | 29233832 |
| | | | | | | | | EAS meta | - | - | 1.136 | 0.023 | 2.43E-08 | | |

| | | | | | | | | | | | | | | | |
|---|---|---|---|---|---|---|---|---|---|---|---|---|---|---|---|
| | | | | | | | | *HK GWAS* | *0.600* | *0.579* | *1.089* | *0.044* | *5.33E-02* | | |
| | | | | | | | | *CC GWAS* | *0.598* | *0.562* | *1.153* | *0.050* | *4.60E-03* | | |
| | | | | | | | | *GZ GWAS* | *0.619* | *0.582* | *1.172* | *0.058* | *6.63E-03* | | |
| | | | | | | | | *KR IC* | *0.608* | *0.581* | *1.124* | *0.044* | *9.75E-03* | | |
| | | | | | | | | *BJ IC* | *0.567* | *0.527* | *1.176* | *0.093* | *7.51E-02* | | |
| | | | | | | | | *MC IC* | *0.607* | *0.550* | *1.266* | *0.120* | *5.30E-02* | | |
| | | | | | | | | EUR meta | - | - | 1.098 | 0.031 | 2.44E-03 | | |
| | | | | | | | | *EUR GWAS1* | *0.297* | *0.252* | *1.225* | *0.093* | *2.96E-02* | | |
| | | | | | | | | *EUR GWAS2* | *0.280* | *0.260* | *1.096* | *0.040* | *2.29E-02* | | |
| | | | | | | | | *EUR GWAS3* | *0.262* | *0.275* | *0.939* | *0.099* | *5.25E-01* | | |
| | | | | | | | | *SP GWAS* | - | - | *1.122* | *0.068* | *1.93E-01* | | |
| **#75** | **rs2289583** | **15** | **75311036** | **C** | **A** | **SCAMP5** | **intronic** | **Trans meta** | **-** | **-** | **1.113** | **0.023** | **2.36E-06** | 0.119 | 26502338 |
| | | | | | | | | EAS meta | - | - | 1.068 | 0.035 | 6.10E-02 | | |
| | | | | | | | | *HK GWAS* | *0.200* | *0.193* | *1.059* | *0.055* | *2.96E-01* | | |
| | | | | | | | | *CC GWAS* | *0.234* | *0.222* | *1.067* | *0.059* | *2.75E-01* | | |
| | | | | | | | | *GZ GWAS* | *0.215* | *0.201* | *1.086* | *0.072* | *2.54E-01* | | |
| | | | | | | | | EUR meta | - | - | 1.148 | 0.030 | 4.16E-06 | | |
| | | | | | | | | *EUR GWAS1* | *0.320* | *0.297* | *1.115* | *0.090* | *2.23E-01* | | |
| | | | | | | | | *EUR GWAS2* | *0.326* | *0.294* | *1.136* | *0.039* | *1.11E-03* | | |
| | | | | | | | | *EUR GWAS3* | *0.308* | *0.281* | *1.139* | *0.098* | *1.81E-01* | | |
| | | | | | | | | *SP GWAS* | - | - | *1.202* | *0.065* | *1.43E-03* | | |
| **#76** | **rs12599402** | **16** | **11189888** | **T** | **C** | **CLEC16A** | **intronic** | **Trans meta** | **-** | **-** | **0.841** | **0.018** | **2.74E-22** | 0.850 | 20805369 |
| | | | | | | | | EAS meta | - | - | 0.843 | 0.023 | 1.63E-13 | | |
| | | | | | | | | *HK GWAS* | *0.391* | *0.435* | *0.829* | *0.044* | *2.04E-05* | | |
| | | | | | | | | *CC GWAS* | *0.362* | *0.403* | *0.832* | *0.052* | *3.94E-04* | | |

| | | | | | | | | | | | | | | |
|---|---|---|---|---|---|---|---|---|---|---|---|---|---|---|
| | | | | | | | | GZ GWAS | 0.403 | 0.460 | 0.789 | 0.059 | 5.79E-05 | | |
| | | | | | | | | KR IC | 0.325 | 0.355 | 0.870 | 0.045 | 2.85E-03 | | |
| | | | | | | | | BJ IC | 0.371 | 0.391 | 0.920 | 0.096 | 3.85E-01 | | |
| | | | | | | | | MC IC | 0.396 | 0.416 | 0.920 | 0.118 | 4.86E-01 | | |
| | | | | | | | | EUR meta | - | - | 0.837 | 0.028 | 2.66E-10 | | |
| | | | | | | | | EUR GWAS1 | 0.433 | 0.469 | 0.863 | 0.086 | 8.60E-02 | | |
| | | | | | | | | EUR GWAS2 | 0.392 | 0.428 | 0.866 | 0.037 | 1.04E-04 | | |
| | | | | | | | | EUR GWAS3 | 0.474 | 0.493 | 0.929 | 0.087 | 3.96E-01 | | |
| | | | | | | | | SP GWAS | - | - | 0.714 | 0.061 | 8.33E-09 | | |
| #77 | rs4592664 | 16 | 23890735 | T | C | PRKCB | intronic | Trans meta | - | - | 0.943 | 0.024 | 1.44E-02 | 2.23E-05 | 21134959 |
| | | | | | | | | EAS meta | - | - | 0.852 | 0.034 | 2.37E-06 | | |
| | | | | | | | | HK GWAS | 0.219 | 0.234 | 0.922 | 0.051 | 1.13E-01 | | |
| | | | | | | | | AH GWAS | 0.229 | 0.278 | 0.771 | 0.059 | 1.07E-05 | | |
| | | | | | | | | GZ GWAS | 0.228 | 0.254 | 0.846 | 0.071 | 1.77E-02 | | |
| | | | | | | | | EUR meta | - | - | 1.046 | 0.035 | 0.190 | | |
| | | | | | | | | EUR GWAS1 | 0.209 | 0.208 | 0.993 | 0.101 | 9.41E-01 | | |
| | | | | | | | | EUR GWAS2 | 0.201 | 0.196 | 1.052 | 0.045 | 2.68E-01 | | |
| | | | | | | | | EUR GWAS3 | 0.204 | 0.176 | 1.209 | 0.111 | 8.72E-02 | | |
| | | | | | | | | Spain GWAS | - | - | 0.994 | 0.075 | 8.79E-01 | | |
| #78 | rs1143679 | 16 | 31276811 | G | A | ITGAM | missense | Trans meta | - | - | 1.797 | 0.038 | 0.00E+00 | 0.488 | 18204448 |
| | | | | | | | | EAS meta | - | - | 2.276 | 0.174 | 2.36E-06 | | |
| | | | | | | | | HK GWAS | 0.014 | 0.007 | 2.021 | 0.215 | 1.05E-03 | | |
| | | | | | | | | CC GWAS | 0.005 | 0.002 | 3.064 | 0.428 | 8.96E-03 | | |
| | | | | | | | | GZ GWAS | 0.010 | 0.004 | 2.684 | 0.415 | 1.74E-02 | | |
| | | | | | | | | EUR meta | - | - | 1.776 | 0.038 | 1.49E-50 | | |

| | | | | | | | | | | | | | | | |
|---|---|---|---|---|---|---|---|---|---|---|---|---|---|---|---|
| | | | | | | | | *EUR GWAS1* | *0.234* | *0.153* | *1.684* | *0.113* | *4.17E-06* | | |
| | | | | | | | | *EUR GWAS2* | *0.167* | *0.104* | *1.770* | *0.054* | *2.79E-26* | | |
| | | | | | | | | *EUR GWAS3* | *0.248* | *0.155* | *1.788* | *0.111* | *1.71E-07* | | |
| | | | | | | | | *SP GWAS* | *-* | *-* | *1.825* | *0.076* | *2.11E-13* | | |
| **#79** | **rs223881** | **16** | **57386566** | **T** | **C** | **PLLP;CCL22** | **intergenic** | **Trans meta** | **-** | **-** | **0.865** | **0.019** | **5.86E-15** | 0.710 | 28108556 |
| | | | | | | | | EAS meta | - | - | 0.869 | 0.023 | 6.97E-10 | | |
| | | | | | | | | *HK GWAS* | *0.532* | *0.575* | *0.833* | *0.043* | *2.50E-05* | | |
| | | | | | | | | *CC GWAS* | *0.536* | *0.539* | *0.973* | *0.050* | *5.82E-01* | | |
| | | | | | | | | *GZ GWAS* | *0.539* | *0.576* | *0.869* | *0.058* | *1.52E-02* | | |
| | | | | | | | | *KR IC* | *0.451* | *0.483* | *0.880* | *0.044* | *2.87E-03* | | |
| | | | | | | | | *BJ IC* | *0.457* | *0.528* | *0.750* | *0.093* | *1.58E-03* | | |
| | | | | | | | | *MC IC* | *0.509* | *0.584* | *0.740* | *0.161* | *1.06E-02* | | |
| | | | | | | | | EUR meta | - | - | 0.856 | 0.032 | 1.57E-06 | | |
| | | | | | | | | *EUR GWAS1* | *0.762* | *0.772* | *0.944* | *0.102* | *5.71E-01* | | |
| | | | | | | | | *EUR GWAS2* | *0.733* | *0.767* | *0.831* | *0.042* | *9.29E-06* | | |
| | | | | | | | | *EUR GWAS3* | *0.753* | *0.764* | *0.938* | *0.105* | *5.41E-01* | | |
| | | | | | | | | *SP GWAS* | *-* | *-* | *0.854* | *0.071* | *1.83E-01* | | |
| **#80** | **rs2731783** | **16** | **58253460** | **A** | **G** | **CSNK2A2** | **intergenic** | **Trans meta** | **-** | **-** | **0.890** | **0.023** | **6.68E-07** | 0.564 | 29625966 |
| | | | | | | | | EAS meta | - | - | 0.881 | 0.030 | 2.42E-05 | | |
| | | | | | | | | *HK GWAS* | *0.674* | *0.696* | *0.900* | *0.046* | *2.35E-02* | | |
| | | | | | | | | *CC GWAS* | *0.619* | *0.658* | *0.842* | *0.052* | *8.35E-04* | | |
| | | | | | | | | *GZ GWAS* | *0.675* | *0.699* | *0.903* | *0.062* | *1.01E-01* | | |
| | | | | | | | | EUR meta | - | - | 0.905 | 0.037 | 7.23E-03 | | |
| | | | | | | | | *EUR GWAS1* | *0.819* | *0.816* | *1.015* | *0.106* | *8.89E-01* | | |
| | | | | | | | | *EUR GWAS2* | *0.823* | *0.848* | *0.817* | *0.049* | *3.25E-05* | | |

| | | | | | | | | | | | | | | |
|---|---|---|---|---|---|---|---|---|---|---|---|---|---|---|
| | | | | | | | | *EUR GWAS3* | *0.861* | *0.858* | *1.022* | *0.127* | *8.61E-01* | | |
| | | | | | | | | *SP GWAS* | *-* | *-* | *1.067* | *0.080* | *3.01E-01* | | |
| **#81** | **rs1170426** | **16** | **68603798** | **C** | **T** | **ZFP90** | **intronic** | **Trans meta** | **-** | **-** | **0.900** | **0.024** | **8.30E-06** | 0.767 | 27399966 |
| | | | | | | | | EAS meta | - | - | 0.893 | 0.036 | 1.44E-03 | | |
| | | | | | | | | *HK GWAS* | *0.794* | *0.812* | *0.893* | *0.054* | *3.60E-02* | | |
| | | | | | | | | *CC GWAS* | *0.810* | *0.832* | *0.863* | *0.063* | *2.04E-02* | | |
| | | | | | | | | *GZ GWAS* | *0.788* | *0.798* | *0.931* | *0.072* | *3.18E-01* | | |
| | | | | | | | | EUR meta | - | - | 0.905 | 0.032 | 1.75E-03 | | |
| | | | | | | | | *EUR GWAS1* | *0.760* | *0.748* | *1.064* | *0.094* | *5.08E-01* | | |
| | | | | | | | | *EUR GWAS2* | *0.746* | *0.767* | *0.889* | *0.042* | *5.26E-03* | | |
| | | | | | | | | *EUR GWAS3* | *0.725* | *0.749* | *0.893* | *0.097* | *2.45E-01* | | |
| | | | | | | | | *SP GWAS* | *-* | *-* | *0.877* | *0.069* | *1.62E-02* | | |
| **#82** | **rs2934498** | **16** | **85968282** | **A** | **G** | **IRF8** | **intergenic** | **Trans meta** | **-** | **-** | **1.155** | **0.022** | **2.56E-11** | 0.857 | 25890262 |
| | | | | | | | | EAS meta | - | - | 1.151 | 0.031 | 4.55E-06 | | |
| | | | | | | | | *HK GWAS* | *0.310* | *0.294* | *1.091* | *0.047* | *6.61E-02* | | |
| | | | | | | | | *CC GWAS* | *0.377* | *0.334* | *1.223* | *0.052* | *1.06E-04* | | |
| | | | | | | | | *GZ GWAS* | *0.332* | *0.301* | *1.158* | *0.064* | *2.13E-02* | | |
| | | | | | | | | EUR meta | - | - | 1.160 | 0.031 | 1.25E-06 | | |
| | | | | | | | | *EUR GWAS1* | *0.302* | *0.300* | *1.000* | *0.091* | *9.98E-01* | | |
| | | | | | | | | *EUR GWAS2* | *0.296* | *0.266* | *1.146* | *0.040* | *7.27E-04* | | |
| | | | | | | | | *EUR GWAS3* | *0.313* | *0.269* | *1.245* | *0.099* | *2.63E-02* | | |
| | | | | | | | | *SP GWAS* | *-* | *-* | *1.254* | *0.066* | *8.35E-04* | | |
| **#83** | **rs11644034** | **16** | **85972612** | **G** | **A** | **IRF8;LINC01082** | **intergenic** | **Trans meta** | **-** | **-** | **0.777** | **0.030** | **5.30E-17** | 0.194 | (1)26502338 |
| | | | | | | | | EAS meta | - | - | 0.822 | 0.053 | 2.33E-04 | | (2)22464253 |
| | | | | | | | | *HK GWAS* | *0.075* | *0.093* | *0.782* | *0.080* | *2.30E-03* | | |

| | | | | | | | | | | | | | | | |
|---|---|---|---|---|---|---|---|---|---|---|---|---|---|---|---|
| | | | | | | | | CC GWAS | 0.077 | 0.088 | 0.868 | 0.093 | 1.29E-01 | | |
| | | | | | | | | GZ GWAS | 0.068 | 0.080 | 0.836 | 0.109 | 1.01E-01 | | |
| | | | | | | | | EUR meta | - | - | 0.756 | 0.037 | 2.17E-14 | | |
| | | | | | | | | EUR GWAS1 | 0.138 | 0.163 | 0.842 | 0.116 | 1.36E-01 | | |
| | | | | | | | | EUR GWAS2 | 0.194 | 0.239 | 0.752 | 0.045 | 2.08E-10 | | |
| | | | | | | | | EUR GWAS3 | 0.091 | 0.135 | 0.652 | 0.145 | 3.23E-03 | | |
| | | | | | | | | SP GWAS | - | - | 0.767 | 0.088 | 2.50E-03 | | |
| **#84** | **rs2280381** | **16** | **86018633** | **C** | **T** | **IRF8;LINC01082** | **intergenic** | **Trans meta** | **-** | **-** | **1.231** | **0.023** | **3.94E-19** | 8.39E-03 | 22046141 |
| | | | | | | | | EAS meta | - | - | 1.323 | 0.036 | 6.75E-15 | | |
| | | | | | | | | HK GWAS | 0.902 | 0.874 | 1.338 | 0.070 | 3.25E-05 | | |
| | | | | | | | | CC GWAS | 0.885 | 0.862 | 1.216 | 0.077 | 1.14E-02 | | |
| | | | | | | | | GZ GWAS | 0.901 | 0.881 | 1.233 | 0.092 | 2.29E-02 | | |
| | | | | | | | | KR IC | 0.893 | 0.854 | 1.429 | 0.068 | 4.83E-08 | | |
| | | | | | | | | BJ IC | 0.894 | 0.866 | 1.299 | 0.138 | 5.84E-02 | | |
| | | | | | | | | MC IC | 0.921 | 0.880 | 1.587 | 0.199 | 1.98E-02 | | |
| | | | | | | | | EUR meta | - | - | 1.168 | 0.030 | 3.12E-07 | | |
| | | | | | | | | EUR GWAS1 | 0.661 | 0.650 | 1.037 | 0.091 | 6.87E-01 | | |
| | | | | | | | | EUR GWAS2 | 0.643 | 0.610 | 1.185 | 0.040 | 2.27E-05 | | |
| | | | | | | | | EUR GWAS3 | 0.738 | 0.701 | 1.224 | 0.105 | 5.38E-02 | | |
| | | | | | | | | SP GWAS | - | - | 1.176 | 0.064 | 1.34E-02 | | |
| #85 | **rs34562254** | **17** | **16842991** | **G** | **A** | **TNFRSF13B** | **missense** | Trans meta | - | - | 1.124 | 0.024 | 1.53E-06 | 3.58E-04 | 29233832 |
| | | | | | | | | **EAS meta** | **-** | **-** | **1.176** | **0.029** | **2.88E-08** | | |
| | | | | | | | | HK GWAS | 0.501 | 0.455 | 1.198 | 0.044 | 4.18E-05 | | |
| | | | | | | | | CC GWAS | 0.383 | 0.358 | 1.106 | 0.052 | 5.43E-02 | | |
| | | | | | | | | GZ GWAS | 0.489 | 0.443 | 1.229 | 0.058 | 4.22E-04 | | |

| | | | | | | | | | | | | | | | |
|---|---|---|---|---|---|---|---|---|---|---|---|---|---|---|---|
| | | | | | | | | EUR meta | - | - | 1.014 | 0.044 | 7.57E-01 | | |
| | | | | | | | | *EUR GWAS1* | *0.108* | *0.111* | *0.984* | *0.135* | *9.05E-01* | | |
| | | | | | | | | *EUR GWAS2* | *0.111* | *0.107* | *1.038* | *0.059* | *5.25E-01* | | |
| | | | | | | | | *EUR GWAS3* | *0.154* | *0.148* | *1.052* | *0.124* | *6.86E-01* | | |
| | | | | | | | | SP GWAS | - | - | 0.939 | 0.101 | 5.75E-01 | | |
| **#86** | **rs8079075** | **17** | **38010815** | **A** | **G** | **IKZF3** | **intron** | **EUR meta** | **-** | **-** | **1.486** | **0.071** | **2.15E-08** | Mononorphic | 22464253 |
| | | | | | | | | *EUR GWAS1* | *0.037* | *0.036* | *1.049* | *0.221* | *8.30E-01* | SNV in EAS | |
| | | | | | | | | *EUR GWAS2* | *0.049* | *0.033* | *1.494* | *0.090* | *8.14E-06* | | |
| | | | | | | | | *EUR GWAS3* | *0.027* | *0.019* | *1.467* | *0.295* | *1.94E-01* | | |
| | | | | | | | | *SP GWAS* | *-* | *-* | *1.733* | *0.151* | *3.40E-04* | | |
| **#87** | **rs930297** | **17** | **73404537** | **C** | **T** | **GRB2** | **intergenic** | **Trans meta** | **-** | **-** | **1.151** | **0.033** | **1.60E-05** | 0.766 | 28714469 |
| | | | | | | | | EAS meta | - | - | 1.120 | 0.051 | 2.67E-02 | | |
| | | | | | | | | *HK GWAS* | *0.889* | *0.887* | *1.033* | *0.073* | *6.56E-01* | | |
| | | | | | | | | *CC GWAS* | *0.053* | *0.052* | *1.003* | *0.109* | *9.81E-01* | | |
| | | | | | | | | *GZ GWAS* | *0.893* | *0.861* | *1.327* | *0.108* | *8.79E-03* | | |
| | | | | | | | | *KR IC* | *0.974* | *0.968* | *1.235* | *0.128* | *9.60E-02* | | |
| | | | | | | | | *BJ IC* | *0.958* | *0.951* | *1.176* | *0.221* | *4.65E-01* | | |
| | | | | | | | | *MC IC* | *0.904* | *0.916* | *0.855* | *0.205* | *4.58E-01* | | |
| | | | | | | | | EUR meta | - | - | 1.171 | 0.042 | 1.69E-04 | | |
| | | | | | | | | *EUR GWAS1* | *0.857* | *0.843* | *1.119* | *0.118* | *3.38E-01* | | |
| | | | | | | | | *EUR GWAS2* | *0.900* | *0.882* | *1.185* | *0.059* | *4.33E-03* | | |
| | | | | | | | | *EUR GWAS3* | *0.835* | *0.814* | *1.161* | *0.114* | *1.91E-01* | | |
| | | | | | | | | *SP GWAS* | *-* | *-* | *1.177* | *0.086* | *4.50E-02* | | |
| **#88** | **rs763361** | **18** | **67531642** | **T** | **C** | **CD226** | **missense** | **Trans meta** | **-** | **-** | **0.883** | **0.018** | **3.03E-12** | 0.477 | 29625966 |
| | | | | | | | | EAS meta | - | - | 0.893 | 0.023 | 9.76E-07 | | |

| | | | | | | | | | | | | | | | |
|---|---|---|---|---|---|---|---|---|---|---|---|---|---|---|---|
| | | | | | | | | HK GWAS | 0.649 | 0.662 | 0.943 | 0.045 | 1.98E-01 | | |
| | | | | | | | | CC GWAS | 0.635 | 0.661 | 0.892 | 0.052 | 2.75E-02 | | |
| | | | | | | | | GZ GWAS | 0.649 | 0.668 | 0.924 | 0.061 | 1.93E-01 | | |
| | | | | | | | | KR IC | 0.571 | 0.605 | 0.870 | 0.043 | 1.00E-03 | | |
| | | | | | | | | BJ IC | 0.631 | 0.684 | 0.787 | 0.092 | 1.34E-02 | | |
| | | | | | | | | MC IC | 0.616 | 0.671 | 0.787 | 0.124 | 5.19E-02 | | |
| | | | | | | | | EUR meta | - | - | 0.870 | 0.028 | 5.17E-07 | | |
| | | | | | | | | EUR GWAS1 | 0.525 | 0.584 | 0.798 | 0.084 | 6.97E-03 | | |
| | | | | | | | | EUR GWAS2 | 0.503 | 0.532 | 0.872 | 0.036 | 1.57E-04 | | |
| | | | | | | | | EUR GWAS3 | 0.441 | 0.462 | 0.923 | 0.088 | 3.68E-01 | | |
| | | | | | | | | SP GWAS | - | - | 0.881 | 0.060 | 4.68E-02 | | |
| **#89** | **rs2304256** | **19** | **10475652** | **C** | **A** | **TYK2** | **missense** | Trans meta | - | - | 0.931 | 0.018 | 8.17E-05 | 2.25E-07 | 26502338 |
| | | | | | | | | EAS meta | - | - | 0.993 | 0.022 | 7.58E-01 | | |
| | | | | | | | | HK GWAS | 0.570 | 0.573 | 0.978 | 0.044 | 6.07E-01 | | |
| | | | | | | | | CC GWAS | 0.465 | 0.464 | 0.990 | 0.050 | 8.41E-01 | | |
| | | | | | | | | GZ GWAS | 0.563 | 0.563 | 1.017 | 0.058 | 7.65E-01 | | |
| | | | | | | | | KR IC | 0.395 | 0.400 | 0.980 | 0.040 | 6.00E-01 | | |
| | | | | | | | | BJ IC | 0.453 | 0.439 | 1.060 | 0.088 | 5.35E-01 | | |
| | | | | | | | | MC IC | 0.576 | 0.570 | 1.030 | 0.091 | 8.23E-01 | | |
| | | | | | | | | **EUR meta** | **-** | **-** | **0.811** | **0.032** | **8.12E-11** | | |
| | | | | | | | | EUR GWAS1 | 0.221 | 0.263 | 0.817 | 0.097 | 3.82E-02 | | |
| | | | | | | | | EUR GWAS2 | 0.240 | 0.287 | 0.785 | 0.042 | 8.79E-09 | | |
| | | | | | | | | EUR GWAS3 | 0.251 | 0.278 | 0.871 | 0.102 | 1.76E-01 | | |
| | | | | | | | | SP GWAS | - | - | 0.857 | 0.071 | 3.05E-02 | | |
| **#90** | **rs2305772** | **19** | **52033742** | **G** | **A** | **SIGLEC6** | **missense** | **Trans meta** | **-** | **-** | **0.914** | **0.018** | **4.50E-07** | 0.514 | 26808113 |

| | | | | | | | | | | | | | | | |
|---|---|---|---|---|---|---|---|---|---|---|---|---|---|---|---|
| | | | | | | | | EAS meta | - | - | 0.905 | 0.023 | 1.73E-05 | | |
| | | | | | | | | *HK GWAS* | *0.404* | *0.418* | *0.943* | *0.044* | *1.82E-01* | | |
| | | | | | | | | *CC GWAS* | *0.432* | *0.446* | *0.948* | *0.050* | *2.88E-01* | | |
| | | | | | | | | *GZ GWAS* | *0.394* | *0.415* | *0.914* | *0.059* | *1.25E-01* | | |
| | | | | | | | | *KR IC* | *0.415* | *0.460* | *0.830* | *0.047* | *2.03E-05* | | |
| | | | | | | | | *BJ IC* | *0.412* | *0.442* | *0.880* | *0.095* | *1.80E-01* | | |
| | | | | | | | | *MC IC* | *0.388* | *0.409* | *0.910* | *0.124* | *4.54E-01* | | |
| | | | | | | | | EUR meta | - | - | 0.927 | 0.028 | 6.42E-03 | | |
| | | | | | | | | *EUR GWAS1* | *0.581* | *0.615* | *0.882* | *0.083* | *1.30E-01* | | |
| | | | | | | | | *EUR GWAS2* | *0.587* | *0.600* | *0.960* | *0.037* | *2.71E-01* | | |
| | | | | | | | | *EUR GWAS3* | *0.548* | *0.567* | *0.928* | *0.088* | *3.95E-01* | | |
| | | | | | | | | *SP GWAS* | - | - | *0.863* | *0.061* | *3.87E-03* | | |
| **#91** | **rs4810485** | **20** | **44747947** | **T** | **G** | **CD40** | **intronic** | **Trans meta** | **-** | **-** | **0.939** | **0.018** | **5.81E-04** | 0.432 | 21914625 |
| | | | | | | | | EAS meta | - | - | 0.929 | 0.023 | 1.20E-03 | | |
| | | | | | | | | *HK GWAS* | *0.512* | *0.527* | *0.948* | *0.044* | *2.23E-01* | | |
| | | | | | | | | *CC GWAS* | *0.606* | *0.627* | *0.929* | *0.051* | *1.52E-01* | | |
| | | | | | | | | *GZ GWAS* | *0.529* | *0.535* | *0.963* | *0.058* | *5.12E-01* | | |
| | | | | | | | | *KR IC* | *0.631* | *0.659* | *0.885* | *0.043* | *5.39E-03* | | |
| | | | | | | | | *BJ IC* | *0.675* | *0.670* | *1.020* | *0.095* | *8.09E-01* | | |
| | | | | | | | | *MC IC* | *0.537* | *0.577* | *0.855* | *0.120* | *1.81E-01* | | |
| | | | | | | | | EUR meta | - | - | 0.957 | 0.031 | 1.61E-01 | | |
| | | | | | | | | *EUR GWAS1* | *0.743* | *0.737* | *1.047* | *0.095* | *6.30E-01* | | |
| | | | | | | | | *EUR GWAS2* | *0.742* | *0.751* | *0.948* | *0.041* | *1.99E-01* | | |
| | | | | | | | | *EUR GWAS3* | *0.675* | *0.686* | *0.952* | *0.095* | *6.07E-01* | | |
| | | | | | | | | *SP GWAS* | - | - | *0.941* | *0.067* | *3.14E-01* | | |

| | | | | | | | | | | | | | | | |
|---|---|---|---|---|---|---|---|---|---|---|---|---|---|---|---|
| **#92** | **rs463426** | **22** | **21809185** | **T** | **C** | **HIC2;TMEM191C** | **intergenic** | **Trans meta** | **-** | **-** | **0.868** | **0.018** | **6.57E-15** | 0.023 | 19838193 |
| | | | | | | | | EAS meta | - | - | 0.841 | 0.023 | 3.60E-14 | | |
| | | | | | | | | *HK GWAS* | *0.411* | *0.448* | *0.856* | *0.044* | *4.88E-04* | | |
| | | | | | | | | *CC GWAS* | *0.464* | *0.522* | *0.797* | *0.050* | *6.72E-06* | | |
| | | | | | | | | *GZ GWAS* | *0.417* | *0.444* | *0.877* | *0.059* | *2.52E-02* | | |
| | | | | | | | | *KR IC* | *0.509* | *0.564* | *0.800* | *0.043* | *2.14E-07* | | |
| | | | | | | | | *BJ IC* | *0.533* | *0.533* | *1.000* | *0.093* | *9.71E-01* | | |
| | | | | | | | | *MC IC* | *0.410* | *0.432* | *0.917* | *0.121* | *4.61E-01* | | |
| | | | | | | | | EUR meta | - | - | 0.916 | 0.030 | 3.56E-03 | | |
| | | | | | | | | *EUR GWAS1* | *0.686* | *0.717* | *0.872* | *0.092* | *1.37E-01* | | |
| | | | | | | | | *EUR GWAS2* | *0.692* | *0.705* | *0.936* | *0.039* | *9.28E-02* | | |
| | | | | | | | | *EUR GWAS3* | *0.675* | *0.703* | *0.887* | *0.093* | *1.99E-01* | | |
| | | | | | | | | *SP GWAS* | - | - | *0.899* | *0.066* | *1.93E-01* | | |
| **#93** | **rs7444** | **22** | **21976934** | **T** | **C** | **UBE2L3** | **UTR3** | **Trans meta** | **-** | **-** | **1.220** | **0.021** | **8.99E-22** | 0.648 | 26502338 |
| | | | | | | | | EAS meta | - | - | 1.215 | 0.023 | 8.56E-18 | | |
| | | | | | | | | *HK GWAS* | *0.602* | *0.555* | *1.213* | *0.044* | *1.27E-05* | | |
| | | | | | | | | *CC GWAS* | *0.525* | *0.462* | *1.271* | *0.050* | *1.50E-06* | | |
| | | | | | | | | *GZ GWAS* | *0.588* | *0.557* | *1.163* | *0.059* | *1.05E-02* | | |
| | | | | | | | | *KR IC* | *0.475* | *0.416* | *1.270* | *0.042* | *2.36E-08* | | |
| | | | | | | | | *BJ IC* | *0.446* | *0.434* | *1.050* | *0.089* | *5.97E-01* | | |
| | | | | | | | | *MC IC* | *0.582* | *0.573* | *1.030* | *0.086* | *7.75E-01* | | |
| | | | | | | | | EUR meta | - | - | 1.246 | 0.051 | 1.87E-05 | | |
| | | | | | | | | *EUR GWAS1* | *0.246* | *0.202* | *1.260* | *0.100* | *2.11E-02* | | |
| | | | | | | | | *EUR GWAS3* | *0.264* | *0.195* | *1.461* | *0.103* | *2.42E-04* | | |
| | | | | | | | | *SP GWAS* | - | - | *1.143* | *0.074* | *1.71E-01* | | |

| | | | | | | | | | | | | | | | |
|---|---|---|---|---|---|---|---|---|---|---|---|---|---|---|---|
| **#94** | **rs61616683** | **22** | **39755773** | **C** | **T** | **SYNGR1** | **intron** | Trans meta | - | - | 1.118 | 0.022 | 3.46E-07 | 0.021 | 26808113 |
| | | | | | | | | **EAS meta** | **-** | **-** | **1.168** | **0.029** | **8.15E-08** | | |
| | | | | | | | | *HK GWAS* | *0.777* | *0.762* | *1.101* | *0.053* | *6.85E-02* | | |
| | | | | | | | | *CC GWAS* | *0.800* | *0.777* | *1.175* | *0.066* | *1.41E-02* | | |
| | | | | | | | | *GZ GWAS* | *0.787* | *0.771* | *1.108* | *0.071* | *1.49E-01* | | |
| | | | | | | | | *KR IC* | *0.869* | *0.843* | *1.235* | *0.059* | *5.75E-04* | | |
| | | | | | | | | *BJ IC* | *0.803* | *0.759* | *1.299* | *0.113* | *1.73E-02* | | |
| | | | | | | | | *MC IC* | *0.814* | *0.772* | *1.299* | *0.148* | *7.78E-02* | | |
| | | | | | | | | EUR meta | - | - | *1.055* | *0.033* | *1.10E-01* | | |
| | | | | | | | | *EUR GWAS1* | *0.224* | *0.221* | *1.031* | *0.101* | *7.64E-01* | | |
| | | | | | | | | *EUR GWAS2* | *0.251* | *0.231* | *1.069* | *0.043* | *1.18E-01* | | |
| | | | | | | | | *EUR GWAS3* | *0.179* | *0.169* | *1.076* | *0.115* | *5.24E-01* | | |
| | | | | | | | | *SP GWAS* | - | - | *1.017* | *0.073* | *6.82E-01* | | |

FA: frequency in affected individuals; FU: frequency in unaffected individuals; Trans meta: association results of transethnic meta-analysis across ten SLE genetic data sets; EAS meta: association results of meta-analysis across genetic data sets from six areas in East Asia (HK: Hong Kong; CC: Central China; GZ: Guangzhou; KR: Korea; BJ: Beijing; MC: Malaysia); EUR meta: association results of meta-analysis across four European GWAS for SLE; IC: Immunochip.

**Extended Data Table 4** Summary association statistics of 38 novel SLE susceptibility loci across different genetic cohorts

| Order | rsid | Chr | Pos (hg19) | AlleleA | AlleleB | Gene context | Annotation | Cohorts | FA_alleleB | FU_alleleB | OR | SE | P | Cochran' s Q-test |
|---|---|---|---|---|---|---|---|---|---|---|---|---|---|---|
| **#1** | rs12093154 | 1 | 1178925 | G | A | C1QTNF12 | nonsynonymous SNV | **Trans meta** | **-** | **-** | 0.835 | **0.030** | **2.51E-09** | 0.648 |
| | | | | | | | | EAS meta | - | - | 0.844 | 0.037 | 4.12E-06 | |
| | | | | | | | | *HK GWAS* | 0.150 | 0.180 | 0.811 | 0.059 | 4.02E-04 | |
| | | | | | | | | *CC GWAS* | 0.218 | 0.250 | 0.845 | 0.060 | 5.32E-03 | |
| | | | | | | | | *GZ GWAS* | 0.165 | 0.178 | 0.897 | 0.076 | 1.55E-01 | |
| | | | | | | | | EUR meta | - | - | 0.819 | 0.052 | 1.38E-04 | |
| | | | | | | | | *EUR GWAS1* | 0.056 | 0.060 | 0.902 | 0.177 | 5.62E-01 | |
| | | | | | | | | *EUR GWAS2* | 0.083 | 0.099 | 0.804 | 0.064 | 6.98E-04 | |
| | | | | | | | | *EUR GWAS3* | 0.064 | 0.069 | 0.916 | 0.182 | 6.29E-01 | |
| | | | | | | | | *SP GWAS* | - | - | 0.794 | 0.127 | 6.11E-02 | |
| **#2** | rs3795310 | 1 | 8431607 | C | T | RERE | intronic | **Trans meta** | **-** | **-** | 0.881 | **0.023** | **3.36E-08** | 0.920 |
| | | | | | | | | EAS meta | - | - | 0.877 | 0.041 | 1.40E-03 | |
| | | | | | | | | *HK GWAS* | 0.120 | 0.145 | 0.794 | 0.065 | 4.71E-04 | |
| | | | | | | | | *CC GWAS* | 0.139 | 0.150 | 0.916 | 0.072 | 2.24E-01 | |
| | | | | | | | | *GZ GWAS* | 0.133 | 0.135 | 0.978 | 0.084 | 7.94E-01 | |
| | | | | | | | | EUR meta | - | - | 0.882 | 0.028 | 8.03E-06 | |
| | | | | | | | | *EUR GWAS1* | 0.512 | 0.579 | 0.747 | 0.087 | 7.36E-04 | |
| | | | | | | | | *EUR GWAS2* | 0.430 | 0.456 | 0.929 | 0.037 | 4.61E-02 | |
| | | | | | | | | *EUR GWAS3* | 0.615 | 0.619 | 0.975 | 0.091 | 7.86E-01 | |
| | | | | | | | | *SP GWAS* | - | - | 0.803 | 0.060 | 1.67E-04 | |
| **#3** | rs28411034 | 1 | 38276997 | G | A | MTF1 | UTR3 | **Trans meta** | **-** | **-** | 0.863 | **0.023** | **1.50E-10** | 0.860 |
| | | | | | | | | EAS meta | - | - | 0.859 | 0.033 | 5.80E-06 | |
| | | | | | | | | *HK GWAS* | 0.218 | 0.259 | 0.797 | 0.052 | 1.27E-05 | |

| | | | | | | | | | | | | | | |
|---|---|---|---|---|---|---|---|---|---|---|---|---|---|---|
| | | | | | | | | *CC GWAS* | 0.263 | 0.276 | 0.937 | 0.059 | 2.66E-01 | |
| | | | | | | | | *GZ GWAS* | 0.243 | 0.269 | 0.871 | 0.065 | 3.32E-02 | |
| | | | | | | | | EUR meta | - | - | 0.866 | 0.032 | 5.93E-06 | |
| | | | | | | | | *EUR GWAS1* | 0.234 | 0.267 | 0.839 | 0.097 | 6.91E-02 | |
| | | | | | | | | *EUR GWAS2* | 0.266 | 0.287 | 0.869 | 0.041 | 5.78E-04 | |
| | | | | | | | | *EUR GWAS3* | 0.224 | 0.254 | 0.850 | 0.105 | 1.22E-01 | |
| | | | | | | | | *SP GWAS* | - | - | 0.881 | 0.071 | 9.52E-02 | |
| **#4** | rs6702599 | 1 | 67825399 | A | C | IL12RB2 | intronic | **Trans meta** | - | - | 0.836 | **0.030** | **3.18E-09** | 0.302 |
| | | | | | | | | EAS meta | - | - | 0.882 | 0.060 | 3.56E-02 | |
| | | | | | | | | *HK GWAS* | 0.935 | 0.943 | 0.875 | 0.089 | 1.31E-01 | |
| | | | | | | | | *CC GWAS* | 0.941 | 0.950 | 0.821 | 0.107 | 6.53E-02 | |
| | | | | | | | | *GZ GWAS* | 0.941 | 0.942 | 0.985 | 0.123 | 9.03E-01 | |
| | | | | | | | | EUR meta | - | - | 0.821 | 0.035 | 1.78E-08 | |
| | | | | | | | | *EUR GWAS1* | 0.784 | 0.830 | 0.768 | 0.105 | 1.16E-02 | |
| | | | | | | | | *EUR GWAS2* | 0.799 | 0.825 | 0.829 | 0.047 | 5.89E-05 | |
| | | | | | | | | *EUR GWAS3* | 0.752 | 0.779 | 0.856 | 0.105 | 1.39E-01 | |
| | | | | | | | | *SP GWAS* | - | - | 0.813 | 0.075 | 2.76E-02 | |
| **#5** | rs1016140 | 1 | 117076547 | G | T | CD58 | intronic | Trans meta | - | - | 0.893 | 0.020 | 1.26E-08 | 0.007 |
| | | | | | | | | **EAS meta** | - | - | 0.870 | **0.022** | **2.95E-10** | |
| | | | | | | | | *HK GWAS* | 0.526 | 0.557 | 0.873 | 0.044 | 2.11E-03 | |
| | | | | | | | | *CC GWAS* | 0.505 | 0.550 | 0.831 | 0.050 | 2.37E-04 | |
| | | | | | | | | *GZ GWAS* | 0.527 | 0.554 | 0.897 | 0.057 | 5.93E-02 | |
| | | | | | | | | *KR IC* | 0.572 | 0.596 | 0.901 | 0.040 | 2.05E-02 | |
| | | | | | | | | *BJ IC* | 0.498 | 0.556 | 0.794 | 0.088 | 1.02E-02 | |
| | | | | | | | | *MC IC* | 0.526 | 0.575 | 0.820 | 0.122 | 9.85E-02 | |

| | | | | | | | | | | | | | |
|---|---|---|---|---|---|---|---|---|---|---|---|---|---|
| | | | | | | | | EUR meta | - | - | 0.998 | 0.046 | 9.62E-01 | |
| | | | | | | | | *EUR GWAS1* | 0.090 | 0.088 | 1.004 | 0.148 | 9.76E-01 | |
| | | | | | | | | *EUR GWAS2* | 0.103 | 0.104 | 1.007 | 0.059 | 9.12E-01 | |
| | | | | | | | | *EUR GWAS3* | 0.100 | 0.119 | 0.811 | 0.144 | 1.46E-01 | |
| | | | | | | | | *SP GWAS* | - | - | 1.072 | 0.099 | 5.84E-01 | |
| **#6** | rs11264750 | 1 | 157497160 | A | G | FCRL5 | intronic | **Trans meta** | - | - | 0.755 | **0.039** | **8.70E-13** | 0.655 |
| | | | | | | | | EAS meta | - | - | 0.751 | 0.041 | 1.95E-12 | |
| | | | | | | | | *HK GWAS* | 0.070 | 0.084 | 0.823 | 0.083 | 1.90E-02 | |
| | | | | | | | | *CC GWAS* | 0.090 | 0.117 | 0.748 | 0.084 | 5.85E-04 | |
| | | | | | | | | *GZ GWAS* | 0.061 | 0.087 | 0.664 | 0.111 | 2.31E-04 | |
| | | | | | | | | *KR IC* | 0.076 | 0.102 | 0.730 | 0.078 | 3.13E-05 | |
| | | | | | | | | *BJ IC* | 0.094 | 0.129 | 0.700 | 0.142 | 1.39E-02 | |
| | | | | | | | | *MC IC* | 0.081 | 0.084 | 0.960 | 0.215 | 8.57E-01 | |
| | | | | | | | | EUR meta | - | - | 0.808 | 0.159 | 1.81E-01 | |
| | | | | | | | | *EUR GWAS1* | 0.011 | 0.009 | 1.290 | 0.448 | 5.69E-01 | |
| | | | | | | | | *EUR GWAS2* | 0.009 | 0.008 | 0.971 | 0.199 | 8.83E-01 | |
| | | | | | | | | *EUR GWAS3* | 0.007 | 0.013 | 0.536 | 0.489 | 2.03E-01 | |
| | | | | | | | | *SP GWAS* | - | - | 0.291 | 0.442 | 4.41E-03 | |
| **#7** | rs12132445 | 1 | 170811799 | G | A | PRRX1;MROH9 | intergenic | Trans meta | - | - | 1.079 | 0.020 | 1.74E-04 | 1.05E-04 |
| | | | | | | | | **EAS meta** | - | - | 1.171 | **0.029** | **6.75E-08** | |
| | | | | | | | | *HK GWAS* | 0.659 | 0.616 | 1.195 | 0.045 | 7.64E-05 | |
| | | | | | | | | *CC GWAS* | 0.646 | 0.605 | 1.178 | 0.052 | 1.50E-03 | |
| | | | | | | | | *GZ GWAS* | 0.647 | 0.625 | 1.123 | 0.059 | 4.93E-02 | |
| | | | | | | | | EUR meta | - | - | 1.001 | 0.028 | 9.67E-01 | |
| | | | | | | | | *EUR GWAS1* | 0.425 | 0.393 | 1.122 | 0.085 | 1.77E-01 | |

| | | | | | | | | | | | | | | |
|---|---|---|---|---|---|---|---|---|---|---|---|---|---|---|
| | | | | | | | | *EUR GWAS2* | 0.455 | 0.458 | 0.992 | 0.036 | 8.33E-01 | |
| | | | | | | | | *EUR GWAS3* | 0.426 | 0.436 | 0.955 | 0.089 | 6.09E-01 | |
| | | | | | | | | *SP GWAS* | - | - | 0.990 | 0.061 | 5.54E-01 | |
| **#8** | rs549669428 | 1 | 174895022 | T | G | RABGAP1L | intronic | **Trans meta** | - | - | 0.839 | **0.032** | **4.53E-08** | 0.062 |
| | | | | | | | | EAS meta | - | - | 0.754 | 0.066 | 1.85E-05 | |
| | | | | | | | | *HK GWAS* | 0.052 | 0.064 | 0.794 | 0.097 | 1.78E-02 | |
| | | | | | | | | *CC GWAS* | 0.050 | 0.071 | 0.659 | 0.120 | 4.87E-04 | |
| | | | | | | | | *GZ GWAS* | 0.047 | 0.056 | 0.811 | 0.135 | 1.21E-01 | |
| | | | | | | | | EUR meta | - | - | 0.869 | 0.037 | 1.67E-04 | |
| | | | | | | | | *EUR GWAS1* | 0.262 | 0.281 | 0.893 | 0.095 | 2.32E-01 | |
| | | | | | | | | *EUR GWAS2* | 0.240 | 0.268 | 0.859 | 0.044 | 5.40E-04 | |
| | | | | | | | | *EUR GWAS3* | 0.250 | 0.268 | 0.898 | 0.107 | 3.15E-01 | |
| **#9** | rs1547624 | 1 | 192543837 | A | T | RGS21;RGS1 | intergenic | Trans meta | - | - | 1.126 | 0.023 | 1.95E-07 | 0.025 |
| | | | | | | | | **EAS meta** | - | - | 1.172 | **0.029** | **4.55E-08** | |
| | | | | | | | | *HK GWAS* | 0.840 | 0.822 | 1.132 | 0.058 | 3.37E-02 | |
| | | | | | | | | *CC GWAS* | 0.814 | 0.798 | 1.112 | 0.064 | 9.74E-02 | |
| | | | | | | | | *GZ GWAS* | 0.850 | 0.818 | 1.267 | 0.077 | 1.99E-03 | |
| | | | | | | | | *KR IC* | 0.780 | 0.747 | 1.205 | 0.053 | 3.11E-04 | |
| | | | | | | | | *BJ IC* | 0.833 | 0.807 | 1.190 | 0.115 | 1.44E-01 | |
| | | | | | | | | *MC IC* | 0.850 | 0.833 | 1.136 | 0.162 | 4.45E-01 | |
| | | | | | | | | EUR meta | - | - | 1.056 | 0.037 | 1.38E-01 | |
| | | | | | | | | *EUR GWAS1* | 0.824 | 0.824 | 1.005 | 0.111 | 9.65E-01 | |
| | | | | | | | | *EUR GWAS2* | 0.829 | 0.821 | 1.041 | 0.048 | 3.95E-01 | |
| | | | | | | | | *EUR GWAS3* | 0.846 | 0.823 | 1.180 | 0.121 | 1.72E-01 | |
| | | | | | | | | *SP GWAS* | - | - | 1.072 | 0.079 | 3.18E-01 | |

| | | | | | | | | | | | | | | |
|---|---|---|---|---|---|---|---|---|---|---|---|---|---|---|
| **#10** | rs2381401 | 2 | 144020974 | C | T | ARHGAP15 | intronic | **Trans meta** | - | - | 1.149 | **0.023** | **1.73E-09** | 0.221 |
| | | | | | | | | EAS meta | - | - | 1.179 | 0.031 | 1.11E-07 | |
| | | | | | | | | *HK GWAS* | 0.331 | 0.299 | 1.154 | 0.047 | 2.05E-03 | |
| | | | | | | | | *CC GWAS* | 0.323 | 0.283 | 1.205 | 0.054 | 4.90E-04 | |
| | | | | | | | | *GZ GWAS* | 0.323 | 0.290 | 1.189 | 0.066 | 8.58E-03 | |
| | | | | | | | | EUR meta | - | - | 1.113 | 0.035 | 1.97E-03 | |
| | | | | | | | | *EUR GWAS1* | 0.228 | 0.215 | 1.069 | 0.102 | 5.12E-01 | |
| | | | | | | | | *EUR GWAS2* | 0.183 | 0.180 | 1.065 | 0.047 | 1.81E-01 | |
| | | | | | | | | *EUR GWAS3* | 0.238 | 0.209 | 1.178 | 0.104 | 1.16E-01 | |
| | | | | | | | | *SP GWAS* | - | - | 1.226 | 0.072 | 2.22E-03 | |
| **#11** | rs9630991 | 2 | 191432139 | G | A | NEMP2;NAB1 | intergenic | **Trans meta** | - | - | 0.851 | **0.022** | **1.08E-13** | 0.971 |
| | | | | | | | | EAS meta | - | - | 0.850 | 0.035 | 3.34E-06 | |
| | | | | | | | | *HK GWAS* | 0.198 | 0.233 | 0.808 | 0.054 | 6.79E-05 | |
| | | | | | | | | *CC GWAS* | 0.198 | 0.231 | 0.824 | 0.062 | 1.71E-03 | |
| | | | | | | | | *GZ GWAS* | 0.221 | 0.228 | 0.963 | 0.069 | 5.85E-01 | |
| | | | | | | | | EUR meta | - | - | 0.851 | 0.028 | 6.73E-09 | |
| | | | | | | | | *EUR GWAS1* | 0.478 | 0.498 | 0.920 | 0.084 | 3.18E-01 | |
| | | | | | | | | *EUR GWAS2* | 0.490 | 0.528 | 0.847 | 0.036 | 4.54E-06 | |
| | | | | | | | | *EUR GWAS3* | 0.468 | 0.512 | 0.835 | 0.090 | 4.46E-02 | |
| | | | | | | | | *SP GWAS* | - | - | 0.838 | 0.060 | 7.52E-03 | |
| **#12** | rs11679484 | 2 | 198921604 | C | A | PLCL1 | intronic | **Trans meta** | - | - | 1.118 | **0.021** | **6.26E-08** | 0.413 |
| | | | | | | | | EAS meta | - | - | 1.147 | 0.034 | 5.59E-05 | |
| | | | | | | | | *HK GWAS* | 0.201 | 0.176 | 1.189 | 0.055 | 1.66E-03 | |
| | | | | | | | | *CC GWAS* | 0.261 | 0.235 | 1.162 | 0.057 | 8.50E-03 | |
| | | | | | | | | *GZ GWAS* | 0.201 | 0.190 | 1.057 | 0.072 | 4.45E-01 | |

| | | | | | | | | | | | | | | |
|---|---|---|---|---|---|---|---|---|---|---|---|---|---|---|
| | | | | | | | | EUR meta | - | - | 1.102 | 0.026 | 1.91E-04 | |
| | | | | | | | | *EUR GWAS1* | 0.419 | 0.402 | 1.077 | 0.086 | 3.90E-01 | |
| | | | | | | | | *EUR GWAS2* | 0.403 | 0.381 | 1.086 | 0.037 | 2.65E-02 | |
| | | | | | | | | *EUR GWAS3* | 0.491 | 0.447 | 1.192 | 0.089 | 4.80E-02 | |
| | | | | | | | | *SP GWAS* | - | - | 1.112 | 0.061 | 9.65E-02 | |
| **#13** | rs3087243 | 2 | 204738919 | G | A | CTLA4 | downstream | **Trans meta** | - | - | 0.895 | **0.020** | **2.29E-08** | 0.396 |
| | | | | | | | | EAS meta | - | - | 0.879 | 0.029 | 6.44E-06 | |
| | | | | | | | | *HK GWAS* | 0.223 | 0.232 | 0.943 | 0.052 | 2.65E-01 | |
| | | | | | | | | *CC GWAS* | 0.156 | 0.180 | 0.830 | 0.068 | 6.19E-03 | |
| | | | | | | | | *GZ GWAS* | 0.219 | 0.239 | 0.895 | 0.069 | 1.10E-01 | |
| | | | | | | | | *KR IC* | 0.153 | 0.173 | 0.870 | 0.056 | 1.34E-02 | |
| | | | | | | | | *BJ IC* | 0.149 | 0.189 | 0.750 | 0.122 | 1.89E-02 | |
| | | | | | | | | *MC IC* | 0.209 | 0.241 | 0.830 | 0.141 | 1.89E-01 | |
| | | | | | | | | EUR meta | - | - | 0.910 | 0.028 | 6.61E-04 | |
| | | | | | | | | *EUR GWAS1* | 0.475 | 0.520 | 0.834 | 0.084 | 2.95E-02 | |
| | | | | | | | | *EUR GWAS2* | 0.430 | 0.442 | 0.968 | 0.037 | 3.79E-01 | |
| | | | | | | | | *EUR GWAS3* | 0.486 | 0.517 | 0.883 | 0.087 | 1.54E-01 | |
| | | | | | | | | *SP GWAS* | - | - | 0.816 | 0.060 | 1.02E-03 | |
| **#14** | rs438613 | 3 | 28072086 | T | C | LINC01980;CMC1 | intergenic | **Trans meta** | - | - | 1.102 | **0.017** | **1.32E-08** | 0.201 |
| | | | | | | | | EAS meta | - | - | 1.125 | 0.022 | 6.46E-08 | |
| | | | | | | | | *HK GWAS* | 0.503 | 0.487 | 1.065 | 0.044 | 1.49E-01 | |
| | | | | | | | | *CC GWAS* | 0.425 | 0.386 | 1.147 | 0.050 | 6.14E-03 | |
| | | | | | | | | *GZ GWAS* | 0.512 | 0.481 | 1.154 | 0.058 | 1.31E-02 | |
| | | | | | | | | *KR IC* | 0.419 | 0.396 | 1.127 | 0.039 | 3.13E-02 | |
| | | | | | | | | *BJ IC* | 0.405 | 0.391 | 1.060 | 0.087 | 5.07E-01 | |

| | | | | | | | | | | | | | | |
|---|---|---|---|---|---|---|---|---|---|---|---|---|---|---|
| | | | | | | | | *MC IC* | 0.542 | 0.451 | 1.440 | 0.110 | 2.11E-03 | |
| | | | | | | | | EUR meta | - | - | 1.066 | 0.028 | 1.98E-02 | |
| | | | | | | | | *EUR GWAS1* | 0.504 | 0.469 | 1.163 | 0.085 | 7.46E-02 | |
| | | | | | | | | *EUR GWAS2* | 0.487 | 0.472 | 1.048 | 0.036 | 1.91E-01 | |
| | | | | | | | | *EUR GWAS3* | 0.548 | 0.513 | 1.149 | 0.089 | 1.17E-01 | |
| | | | | | | | | *SP GWAS* | - | - | 1.035 | 0.060 | 8.87E-01 | |
| **#15** | rs4690055 | 4 | 2748663 | G | A | TNIP2 | intronic | **Trans meta** | - | - | 0.909 | **0.018** | **5.68E-08** | 0.776 |
| | | | | | | | | EAS meta | - | - | 0.911 | 0.023 | 4.14E-05 | |
| | | | | | | | | *HK GWAS* | 0.347 | 0.351 | 0.991 | 0.046 | 8.43E-01 | |
| | | | | | | | | *CC GWAS* | 0.403 | 0.433 | 0.888 | 0.051 | 1.96E-02 | |
| | | | | | | | | *GZ GWAS* | 0.349 | 0.359 | 0.931 | 0.061 | 2.38E-01 | |
| | | | | | | | | *KR IC* | 0.429 | 0.455 | 0.890 | 0.041 | 1.46E-02 | |
| | | | | | | | | *BJ IC* | 0.431 | 0.505 | 0.740 | 0.087 | 9.39E-04 | |
| | | | | | | | | *MC IC* | 0.343 | 0.346 | 0.987 | 0.128 | 9.20E-01 | |
| | | | | | | | | EUR meta | - | - | 0.907 | 0.028 | 4.73E-04 | |
| | | | | | | | | *EUR GWAS1* | 0.437 | 0.455 | 0.937 | 0.082 | 4.29E-01 | |
| | | | | | | | | *EUR GWAS2* | 0.452 | 0.471 | 0.936 | 0.036 | 7.05E-02 | |
| | | | | | | | | *EUR GWAS3* | 0.414 | 0.443 | 0.891 | 0.089 | 1.95E-01 | |
| | | | | | | | | *SP GWAS* | - | - | 0.823 | 0.061 | 3.20E-03 | |
| **#16** | rs4697651 | 4 | 10721433 | C | T | CLNK;MIR572 | intergenic | **Trans meta** | - | - | 0.878 | **0.024** | **6.36E-08** | 0.256 |
| | | | | | | | | EAS meta | - | - | 0.848 | 0.040 | 4.49E-05 | |
| | | | | | | | | *HK GWAS* | 0.139 | 0.161 | 0.830 | 0.061 | 2.62E-03 | |
| | | | | | | | | *CC GWAS* | 0.134 | 0.153 | 0.852 | 0.072 | 2.80E-02 | |
| | | | | | | | | *GZ GWAS* | 0.142 | 0.159 | 0.875 | 0.081 | 1.03E-01 | |
| | | | | | | | | EUR meta | - | - | 0.897 | 0.030 | 3.26E-04 | |

| | | | | | | | | | | | | | | |
|---|---|---|---|---|---|---|---|---|---|---|---|---|---|---|
| | | | | | | | | *EUR GWAS1* | 0.307 | 0.306 | 1.007 | 0.105 | 9.42E-01 | |
| | | | | | | | | *EUR GWAS2* | 0.285 | 0.307 | 0.912 | 0.039 | 2.15E-02 | |
| | | | | | | | | *EUR GWAS3* | 0.242 | 0.276 | 0.843 | 0.100 | 9.08E-02 | |
| | | | | | | | | *SP GWAS* | - | - | 0.839 | 0.065 | 2.07E-02 | |
| **#17** | rs6871748 | 5 | 35885982 | T | C | IL7R;CAPSL | intergenic | **Trans meta** | - | - | 0.886 | **0.022** | **3.96E-08** | 0.579 |
| | | | | | | | | EAS meta | - | - | 0.875 | 0.031 | 1.44E-05 | |
| | | | | | | | | *HK GWAS* | 0.146 | 0.163 | 0.881 | 0.061 | 3.75E-02 | |
| | | | | | | | | *CC GWAS* | 0.133 | 0.150 | 0.867 | 0.073 | 5.11E-02 | |
| | | | | | | | | *GZ GWAS* | 0.159 | 0.175 | 0.898 | 0.077 | 1.64E-01 | |
| | | | | | | | | *KR IC* | 0.169 | 0.187 | 0.880 | 0.055 | 2.44E-02 | |
| | | | | | | | | *BJ IC* | 0.131 | 0.155 | 0.820 | 0.126 | 1.30E-01 | |
| | | | | | | | | *MC IC* | 0.155 | 0.181 | 0.830 | 0.157 | 2.41E-01 | |
| | | | | | | | | EUR meta | - | - | 0.897 | 0.032 | 6.40E-04 | |
| | | | | | | | | *EUR GWAS1* | 0.242 | 0.242 | 1.002 | 0.098 | 9.87E-01 | |
| | | | | | | | | *EUR GWAS2* | 0.256 | 0.278 | 0.892 | 0.041 | 5.21E-03 | |
| | | | | | | | | *EUR GWAS3* | 0.203 | 0.223 | 0.886 | 0.110 | 2.72E-01 | |
| | | | | | | | | *SP GWAS* | - | - | 0.868 | 0.069 | 1.63E-02 | |
| **#18** | rs6927090 | 6 | 252145 | G | T | DUSP22;IRF4 | intergenic | **EAS meta** | - | - | 1.233 | **0.033** | **3.48E-10** | QC Failed in EUR GWAS |
| | | | | | | | | *HK GWAS* | 0.260 | 0.216 | 1.274 | 0.051 | 1.60E-06 | |
| | | | | | | | | *CC GWAS* | 0.246 | 0.228 | 1.108 | 0.058 | 7.76E-02 | |
| | | | | | | | | *GZ GWAS* | 0.263 | 0.209 | 1.348 | 0.069 | 1.51E-05 | |
| **#19** | rs9387400 | 6 | 116694120 | C | A | DSE | intronic | Trans meta | - | - | 0.892 | 0.025 | 5.89E-06 | 1.17E-04 |
| | | | | | | | | **EAS meta** | - | - | 0.743 | **0.054** | **3.14E-08** | |
| | | | | | | | | *HK GWAS* | 0.934 | 0.948 | 0.763 | 0.090 | 2.69E-03 | |
| | | | | | | | | *CC GWAS* | 0.884 | 0.913 | 0.703 | 0.081 | 1.28E-05 | |

| | | | | | | | | | | | | | | |
|---|---|---|---|---|---|---|---|---|---|---|---|---|---|---|
| | | | | | | | | *GZ GWAS* | 0.926 | 0.941 | 0.801 | 0.119 | 6.07E-02 | |
| | | | | | | | | EUR meta | - | - | 0.939 | 0.029 | 2.96E-02 | |
| | | | | | | | | *EUR GWAS1* | 0.378 | 0.389 | 0.958 | 0.084 | 6.11E-01 | |
| | | | | | | | | *EUR GWAS2* | 0.358 | 0.373 | 0.910 | 0.038 | 1.29E-02 | |
| | | | | | | | | *EUR GWAS3* | 0.394 | 0.417 | 0.899 | 0.092 | 2.47E-01 | |
| | | | | | | | | *SP GWAS* | - | - | 1.033 | 0.062 | 5.62E-01 | |
| **#20** | rs13260060 | 8 | 71218360 | G | A | NCOA2 | intronic | Trans meta | - | - | 0.903 | 0.019 | 7.94E-08 | 0.044 |
| | | | | | | | | **EAS meta** | - | - | 0.889 | **0.021** | **1.92E-08** | |
| | | | | | | | | *HK GWAS* | 0.374 | 0.407 | 0.868 | 0.044 | 1.52E-03 | |
| | | | | | | | | *CC GWAS* | 0.358 | 0.368 | 0.944 | 0.051 | 2.58E-01 | |
| | | | | | | | | *GZ GWAS* | 0.369 | 0.405 | 0.859 | 0.059 | 9.92E-03 | |
| | | | | | | | | *KR IC* | 0.394 | 0.423 | 0.890 | 0.036 | 5.32E-03 | |
| | | | | | | | | *BJ IC* | 0.330 | 0.370 | 0.839 | 0.098 | 5.92E-02 | |
| | | | | | | | | *MC IC* | 0.367 | 0.382 | 0.940 | 0.118 | 6.03E-01 | |
| | | | | | | | | EUR meta | - | - | 0.986 | 0.046 | 7.55E-01 | |
| | | | | | | | | *EUR GWAS1* | 0.101 | 0.111 | 0.886 | 0.138 | 3.81E-01 | |
| | | | | | | | | *EUR GWAS2* | 0.101 | 0.097 | 1.075 | 0.061 | 2.36E-01 | |
| | | | | | | | | *EUR GWAS3* | 0.108 | 0.126 | 0.840 | 0.140 | 2.13E-01 | |
| | | | | | | | | *SP GWAS* | - | - | 0.898 | 0.100 | 2.22E-01 | |
| **#21** | rs2445610 | 8 | 128197088 | A | G | PRNCR1;CASC19 | intergenic | Trans meta | - | - | 0.913 | 0.018 | 2.60E-07 | 0.027 |
| | | | | | | | | **EAS meta** | - | - | 0.886 | **0.022** | **5.26E-08** | |
| | | | | | | | | *HK GWAS* | 0.458 | 0.478 | 0.915 | 0.043 | 4.41E-02 | |
| | | | | | | | | *CC GWAS* | 0.381 | 0.416 | 0.862 | 0.050 | 4.02E-03 | |
| | | | | | | | | *GZ GWAS* | 0.457 | 0.492 | 0.867 | 0.056 | 1.07E-02 | |
| | | | | | | | | *KR IC* | 0.404 | 0.434 | 0.880 | 0.041 | 4.43E-03 | |

| | | | | | | | | | | | | | | |
|---|---|---|---|---|---|---|---|---|---|---|---|---|---|---|
| | | | | | | | | *BJ IC* | 0.360 | 0.367 | 0.969 | 0.098 | 7.49E-01 | |
| | | | | | | | | *MC IC* | 0.460 | 0.502 | 0.840 | 0.115 | 1.54E-01 | |
| | | | | | | | | EUR meta | - | - | 0.961 | 0.029 | 1.82E-01 | |
| | | | | | | | | *EUR GWAS1* | 0.319 | 0.312 | 1.036 | 0.091 | 6.97E-01 | |
| | | | | | | | | *EUR GWAS2* | 0.359 | 0.364 | 0.958 | 0.038 | 2.54E-01 | |
| | | | | | | | | *EUR GWAS3* | 0.288 | 0.309 | 0.901 | 0.100 | 2.95E-01 | |
| | | | | | | | | *SP GWAS* | - | - | 0.962 | 0.065 | 5.18E-01 | |
| **#22** | rs7815944 | 8 | 129427518 | A | G | LINC00824 | ncRNA_intronic | Trans meta | - | - | 0.868 | 0.024 | 2.21E-09 | 0.303 |
| | | | | | | | | **EAS meta** | - | - | 0.862 | **0.025** | **1.78E-09** | |
| | | | | | | | | *HK GWAS* | 0.242 | 0.272 | 0.855 | 0.050 | 1.70E-03 | |
| | | | | | | | | *CC GWAS* | 0.283 | 0.316 | 0.865 | 0.055 | 8.41E-03 | |
| | | | | | | | | *GZ GWAS* | 0.233 | 0.266 | 0.832 | 0.066 | 5.19E-03 | |
| | | | | | | | | *KR IC* | 0.343 | 0.369 | 0.890 | 0.044 | 9.34E-03 | |
| | | | | | | | | *BJ IC* | 0.305 | 0.347 | 0.830 | 0.102 | 4.83E-02 | |
| | | | | | | | | *MC IC* | 0.240 | 0.281 | 0.810 | 0.136 | 1.22E-01 | |
| | | | | | | | | EUR meta | - | - | 0.939 | 0.080 | 4.27E-01 | |
| | | | | | | | | *EUR GWAS1* | 0.034 | 0.030 | 1.158 | 0.242 | 5.45E-01 | |
| | | | | | | | | *EUR GWAS2* | 0.031 | 0.034 | 0.940 | 0.103 | 5.50E-01 | |
| | | | | | | | | *EUR GWAS3* | 0.026 | 0.034 | 0.746 | 0.266 | 2.71E-01 | |
| | | | | | | | | *SP GWAS* | - | - | 0.926 | 0.173 | 5.06E-01 | |
| **#23** | rs4978037 | 9 | 21228423 | T | C | IFNB1-IFNE | upstream | **Trans meta** | - | - | 1.121 | **0.021** | **5.68E-08** | 0.902 |
| | | | | | | | | EAS meta | - | - | 1.117 | 0.028 | 7.36E-05 | |
| | | | | | | | | *HK GWAS* | 0.484 | 0.463 | 1.088 | 0.043 | 5.47E-02 | |
| | | | | | | | | *CC GWAS* | 0.526 | 0.489 | 1.162 | 0.050 | 2.74E-03 | |
| | | | | | | | | *GZ GWAS* | 0.511 | 0.484 | 1.111 | 0.058 | 7.01E-02 | |

| | | | | | | | | | | | | | | |
|---|---|---|---|---|---|---|---|---|---|---|---|---|---|---|
| | | | | | | | | EUR meta | - | - | 1.124 | 0.033 | 3.92E-04 | |
| | | | | | | | | *EUR GWAS1* | 0.805 | 0.809 | 0.998 | 0.105 | 9.85E-01 | |
| | | | | | | | | *EUR GWAS2* | 0.799 | 0.784 | 1.126 | 0.043 | 5.65E-03 | |
| | | | | | | | | *EUR GWAS3* | 0.816 | 0.779 | 1.249 | 0.108 | 4.33E-02 | |
| | | | | | | | | *SP GWAS* | - | - | 1.126 | 0.076 | 8.65E-02 | |
| **#24** | rs1405209 | 9 | 102585545 | T | C | NR4A3 | intronic | **Trans meta** | - | - | 1.112 | **0.019** | **2.42E-08** | 0.816 |
| | | | | | | | | EAS meta | - | - | 1.106 | 0.028 | 3.10E-04 | |
| | | | | | | | | *HK GWAS* | 0.158 | 0.150 | 1.071 | 0.060 | 2.50E-01 | |
| | | | | | | | | *CC GWAS* | 0.202 | 0.193 | 1.064 | 0.062 | 3.18E-01 | |
| | | | | | | | | *GZ GWAS* | 0.157 | 0.140 | 1.128 | 0.081 | 1.39E-01 | |
| | | | | | | | | *KR IC* | 0.250 | 0.237 | 1.080 | 0.049 | 1.40E-01 | |
| | | | | | | | | *BJ IC* | 0.248 | 0.191 | 1.400 | 0.109 | 2.32E-03 | |
| | | | | | | | | *MC IC* | 0.166 | 0.132 | 1.300 | 0.165 | 1.16E-01 | |
| | | | | | | | | EUR meta | - | - | 1.116 | 0.028 | 8.55E-05 | |
| | | | | | | | | *EUR GWAS1* | 0.375 | 0.365 | 1.042 | 0.085 | 6.27E-01 | |
| | | | | | | | | *EUR GWAS2* | 0.388 | 0.362 | 1.112 | 0.037 | 4.38E-03 | |
| | | | | | | | | *EUR GWAS3* | 0.334 | 0.340 | 0.970 | 0.094 | 7.48E-01 | |
| | | | | | | | | *SP GWAS* | - | - | 1.245 | 0.062 | 2.34E-04 | |
| **#25** | rs4745876 | 10 | 64425126 | G | A | ZNF365 | intronic | **Trans meta** | - | - | 0.882 | **0.023** | **3.66E-08** | 0.569 |
| | | | | | | | | EAS meta | - | - | 0.876 | 0.027 | 1.36E-06 | |
| | | | | | | | | *HK GWAS* | 0.213 | 0.231 | 0.894 | 0.053 | 3.28E-02 | |
| | | | | | | | | *CC GWAS* | 0.191 | 0.222 | 0.820 | 0.063 | 1.53E-03 | |
| | | | | | | | | *GZ GWAS* | 0.208 | 0.226 | 0.907 | 0.070 | 1.64E-01 | |
| | | | | | | | | *KR IC* | 0.202 | 0.232 | 0.840 | 0.052 | 6.64E-04 | |
| | | | | | | | | *BJ IC* | 0.211 | 0.211 | 0.998 | 0.108 | 9.88E-01 | |

| | | | | | | | | | | | | | | |
|---|---|---|---|---|---|---|---|---|---|---|---|---|---|---|
| | | | | | | | | *MC IC* | 0.202 | 0.201 | 1.005 | 0.147 | 9.76E-01 | |
| | | | | | | | | EUR meta | - | - | 0.901 | 0.042 | 1.40E-02 | |
| | | | | | | | | *EUR GWAS1* | 0.100 | 0.114 | 0.867 | 0.135 | 2.89E-01 | |
| | | | | | | | | *EUR GWAS2* | 0.132 | 0.145 | 0.896 | 0.054 | 4.00E-02 | |
| | | | | | | | | *EUR GWAS3* | 0.109 | 0.106 | 1.042 | 0.145 | 7.77E-01 | |
| | | | | | | | | *SP GWAS* | - | - | 0.878 | 0.098 | 1.21E-01 | |
| **#26** | rs10999979 | 10 | 73501066 | C | A | CDH23 | intronic | **Trans meta** | - | - | 1.165 | **0.028** | **4.65E-08** | 0.726 |
| | | | | | | | | EAS meta | - | - | 1.163 | 0.028 | 1.01E-07 | |
| | | | | | | | | *HK GWAS* | 0.210 | 0.183 | 1.183 | 0.054 | 1.92E-03 | |
| | | | | | | | | *CC GWAS* | 0.214 | 0.181 | 1.220 | 0.062 | 1.37E-03 | |
| | | | | | | | | *GZ GWAS* | 0.214 | 0.201 | 1.092 | 0.072 | 2.21E-01 | |
| | | | | | | | | *KR IC* | 0.176 | 0.158 | 1.140 | 0.055 | 2.00E-02 | |
| | | | | | | | | *BJ IC* | 0.187 | 0.165 | 1.160 | 0.118 | 2.12E-01 | |
| | | | | | | | | *MC IC* | 0.207 | 0.179 | 1.190 | 0.148 | 2.43E-01 | |
| | | | | | | | | EUR meta | - | - | 1.280 | 0.270 | 3.61E-01 | |
| | | | | | | | | *EUR GWAS1* | 0.003 | 0.006 | 0.465 | 0.638 | 2.31E-01 | |
| | | | | | | | | *EUR GWAS3* | 0.015 | 0.008 | 1.733 | 0.398 | 1.67E-01 | |
| | | | | | | | | *SP GWAS* | - | - | 1.435 | 0.448 | 4.84E-01 | |
| **#27** | rs7975703 | 12 | 121099302 | C | T | CABP1 | intronic | **Trans meta** | - | - | 0.884 | **0.022** | **2.99E-08** | 0.367 |
| | | | | | | | | EAS meta | - | - | 0.869 | 0.029 | 1.47E-06 | |
| | | | | | | | | *HK GWAS* | 0.429 | 0.448 | 0.920 | 0.043 | 5.60E-02 | |
| | | | | | | | | *CC GWAS* | 0.362 | 0.413 | 0.798 | 0.053 | 1.78E-05 | |
| | | | | | | | | *GZ GWAS* | 0.433 | 0.469 | 0.873 | 0.058 | 1.90E-02 | |
| | | | | | | | | EUR meta | - | - | 0.905 | 0.034 | 3.89E-03 | |
| | | | | | | | | *EUR GWAS1* | 0.202 | 0.203 | 1.005 | 0.103 | 9.58E-01 | |

| | | | | | | | | | | | | | | |
|---|---|---|---|---|---|---|---|---|---|---|---|---|---|---|
| | | | | | | | | *EUR GWAS2* | 0.196 | 0.213 | 0.881 | 0.046 | 5.66E-03 | |
| | | | | | | | | *EUR GWAS3* | 0.221 | 0.229 | 0.950 | 0.107 | 6.29E-01 | |
| | | | | | | | | *SP GWAS* | - | - | 0.900 | 0.074 | 9.41E-02 | |
| **#28** | rs76725306 | 13 | 50177453 | G | A | RCBTB1;ARL11 | intergenic | **Trans meta** | **-** | **-** | 1.157 | **0.027** | **3.60E-08** | 0.617 |
| | | | | | | | | EAS meta | - | - | 1.150 | 0.029 | 1.38E-06 | |
| | | | | | | | | *HK GWAS* | 0.349 | 0.323 | 1.136 | 0.046 | 5.88E-03 | |
| | | | | | | | | *CC GWAS* | 0.393 | 0.352 | 1.191 | 0.051 | 6.13E-04 | |
| | | | | | | | | *GZ GWAS* | 0.352 | 0.326 | 1.118 | 0.061 | 6.86E-02 | |
| | | | | | | | | EUR meta | - | - | 1.193 | 0.067 | 8.22E-03 | |
| | | | | | | | | *EUR GWAS1* | 0.041 | 0.029 | 1.432 | 0.241 | 1.36E-01 | |
| | | | | | | | | *EUR GWAS2* | 0.052 | 0.045 | 1.170 | 0.084 | 6.16E-02 | |
| | | | | | | | | *EUR GWAS3* | 0.065 | 0.047 | 1.387 | 0.189 | 8.33E-02 | |
| | | | | | | | | *SP GWAS* | - | - | 1.054 | 0.164 | 8.23E-01 | |
| **#29** | rs1885889 | 13 | 100091300 | A | G | UBAC2;LINC01232 | intergenic | **Trans meta** | **-** | **-** | 0.868 | **0.019** | **2.05E-13** | 0.870 |
| | | | | | | | | EAS meta | - | - | 0.870 | 0.023 | 7.93E-10 | |
| | | | | | | | | *HK GWAS* | 0.591 | 0.613 | 0.912 | 0.044 | 3.53E-02 | |
| | | | | | | | | *CC GWAS* | 0.627 | 0.642 | 0.931 | 0.052 | 1.73E-01 | |
| | | | | | | | | *GZ GWAS* | 0.585 | 0.622 | 0.844 | 0.061 | 5.03E-03 | |
| | | | | | | | | *KR IC* | 0.595 | 0.639 | 0.826 | 0.041 | 2.02E-05 | |
| | | | | | | | | *BJ IC* | 0.596 | 0.628 | 0.877 | 0.093 | 1.47E-01 | |
| | | | | | | | | *MC IC* | 0.569 | 0.639 | 0.741 | 0.123 | 1.46E-02 | |
| | | | | | | | | EUR meta | - | - | 0.864 | 0.036 | 5.69E-05 | |
| | | | | | | | | *EUR GWAS1* | 0.821 | 0.828 | 0.956 | 0.110 | 6.85E-01 | |
| | | | | | | | | *EUR GWAS2* | 0.831 | 0.843 | 0.898 | 0.049 | 2.66E-02 | |
| | | | | | | | | *EUR GWAS3* | 0.773 | 0.802 | 0.838 | 0.107 | 1.00E-01 | |

| | | | | | | | | | | | | | | |
|---|---|---|---|---|---|---|---|---|---|---|---|---|---|---|
| | | | | | | | | *SP GWAS* | - | - | 0.758 | 0.077 | 8.31E-04 | |
| **#30** | rs12148050 | 14 | 103263788 | A | G | TRAF3 | intronic | **Trans meta** | **-** | **-** | 0.907 | **0.018** | **2.57E-08** | 0.062 |
| | | | | | | | | EAS meta | - | - | 0.930 | 0.022 | 1.04E-03 | |
| | | | | | | | | *HK GWAS* | 0.556 | 0.572 | 0.937 | 0.043 | 1.33E-01 | |
| | | | | | | | | *CC GWAS* | 0.555 | 0.576 | 0.921 | 0.050 | 1.04E-01 | |
| | | | | | | | | *GZ GWAS* | 0.545 | 0.555 | 0.956 | 0.059 | 4.45E-01 | |
| | | | | | | | | *KR IC* | 0.499 | 0.522 | 0.909 | 0.040 | 2.95E-02 | |
| | | | | | | | | *BJ IC* | 0.575 | 0.572 | 1.010 | 0.090 | 9.11E-01 | |
| | | | | | | | | *MC IC* | 0.532 | 0.566 | 0.870 | 0.119 | 2.41E-01 | |
| | | | | | | | | EUR meta | - | - | 0.869 | 0.029 | 1.11E-06 | |
| | | | | | | | | *EUR GWAS1* | 0.617 | 0.671 | 0.793 | 0.087 | 7.61E-03 | |
| | | | | | | | | *EUR GWAS2* | 0.619 | 0.646 | 0.883 | 0.038 | 1.05E-03 | |
| | | | | | | | | *EUR GWAS3* | 0.640 | 0.664 | 0.902 | 0.092 | 2.63E-01 | |
| | | | | | | | | *SP GWAS* | - | - | 0.857 | 0.062 | 1.32E-02 | |
| **#31** | rs869310 | 15 | 77830306 | T | G | LOC101929457;LINGO1 | intergenic | **Trans meta** | **-** | **-** | 0.881 | **0.023** | **1.90E-08** | 0.294 |
| | | | | | | | | EAS meta | - | - | 0.856 | 0.035 | 1.11E-05 | |
| | | | | | | | | *HK GWAS* | 0.204 | 0.229 | 0.864 | 0.053 | 6.10E-03 | |
| | | | | | | | | *CC GWAS* | 0.186 | 0.206 | 0.881 | 0.064 | 4.76E-02 | |
| | | | | | | | | *GZ GWAS* | 0.205 | 0.240 | 0.814 | 0.070 | 3.33E-03 | |
| | | | | | | | | EUR meta | - | - | 0.898 | 0.029 | 2.54E-04 | |
| | | | | | | | | *EUR GWAS1* | 0.308 | 0.335 | 0.887 | 0.088 | 1.75E-01 | |
| | | | | | | | | *EUR GWAS2* | 0.341 | 0.357 | 0.929 | 0.038 | 5.31E-02 | |
| | | | | | | | | *EUR GWAS3* | 0.320 | 0.347 | 0.882 | 0.095 | 1.85E-01 | |
| | | | | | | | | *SP GWAS* | - | - | 0.830 | 0.064 | 3.39E-03 | |
| **#32** | rs9899849 | 17 | 7234983 | G | A | NEURL4;ACAP1 | intergenic | Trans meta | - | - | 1.078 | 0.023 | 1.08E-03 | 0.000 |

| | | | | | | | | | | | | | | |
|---|---|---|---|---|---|---|---|---|---|---|---|---|---|---|
| | | | | | | | | EAS meta | - | - | 0.969 | 0.032 | 3.32E-01 | |
| | | | | | | | | *HK GWAS* | 0.291 | 0.293 | 0.990 | 0.049 | 8.36E-01 | |
| | | | | | | | | *CC GWAS* | 0.282 | 0.296 | 0.922 | 0.057 | 1.50E-01 | |
| | | | | | | | | *GZ GWAS* | 0.302 | 0.301 | 0.997 | 0.064 | 9.65E-01 | |
| | | | | | | | | **EUR meta** | **-** | **-** | 1.209 | **0.033** | **1.10E-08** | |
| | | | | | | | | *EUR GWAS1* | 0.217 | 0.188 | 1.179 | 0.105 | 1.18E-01 | |
| | | | | | | | | *EUR GWAS2* | 0.265 | 0.225 | 1.237 | 0.042 | 4.24E-07 | |
| | | | | | | | | *EUR GWAS3* | 0.145 | 0.128 | 1.180 | 0.132 | 2.08E-01 | |
| | | | | | | | | *SP GWAS* | - | - | 1.152 | 0.072 | 6.35E-02 | |
| **#33** | rs2384991 | 19 | 18386634 | A | C | KIAA1683;JUND | intergenic | **Trans meta** | **-** | **-** | 1.108 | **0.019** | **4.95E-08** | 0.657 |
| | | | | | | | | EAS meta | - | - | 1.101 | 0.023 | 2.88E-05 | |
| | | | | | | | | *HK GWAS* | 0.352 | 0.335 | 1.087 | 0.045 | 6.50E-02 | |
| | | | | | | | | *CC GWAS* | 0.381 | 0.364 | 1.081 | 0.051 | 1.28E-01 | |
| | | | | | | | | *GZ GWAS* | 0.350 | 0.335 | 1.067 | 0.062 | 2.92E-01 | |
| | | | | | | | | *KR IC* | 0.382 | 0.348 | 1.160 | 0.042 | 9.24E-04 | |
| | | | | | | | | *BJ IC* | 0.404 | 0.384 | 1.090 | 0.092 | 3.72E-01 | |
| | | | | | | | | *MC IC* | 0.328 | 0.328 | 1.002 | 0.127 | 9.84E-01 | |
| | | | | | | | | EUR meta | - | - | 1.121 | 0.032 | 4.22E-04 | |
| | | | | | | | | *EUR GWAS1* | 0.207 | 0.192 | 1.092 | 0.107 | 4.10E-01 | |
| | | | | | | | | *EUR GWAS2* | 0.280 | 0.251 | 1.141 | 0.041 | 1.47E-03 | |
| | | | | | | | | *EUR GWAS3* | 0.212 | 0.206 | 1.040 | 0.109 | 7.23E-01 | |
| | | | | | | | | *SP GWAS* | - | - | 1.113 | 0.071 | 4.38E-02 | |
| **#34** | rs13344313 | 19 | 18517767 | G | A | LRRC25;SSBP4 | intergenic | **Trans meta** | **-** | **-** | 0.861 | **0.023** | **1.34E-10** | 0.959 |
| | | | | | | | | EAS meta | - | - | 0.860 | 0.035 | 1.35E-05 | |
| | | | | | | | | *HK GWAS* | 0.088 | 0.100 | 0.878 | 0.076 | 8.59E-02 | |

| | | | | | | | | | | | | | | |
|---|---|---|---|---|---|---|---|---|---|---|---|---|---|---|
| | | | | | | | | *CC GWAS* | 0.126 | 0.145 | 0.868 | 0.075 | 5.85E-02 | |
| | | | | | | | | *GZ GWAS* | 0.118 | 0.124 | 0.909 | 0.101 | 3.45E-01 | |
| | | | | | | | | *KR IC* | 0.148 | 0.175 | 0.820 | 0.059 | 6.01E-04 | |
| | | | | | | | | *BJ IC* | 0.146 | 0.169 | 0.840 | 0.123 | 1.54E-01 | |
| | | | | | | | | *MC IC* | 0.099 | 0.096 | 1.030 | 0.197 | 8.74E-01 | |
| | | | | | | | | EUR meta | - | - | 0.862 | 0.031 | 2.33E-06 | |
| | | | | | | | | *EUR GWAS1* | 0.235 | 0.288 | 0.748 | 0.096 | 2.43E-03 | |
| | | | | | | | | *EUR GWAS2* | 0.264 | 0.282 | 0.915 | 0.041 | 3.04E-02 | |
| | | | | | | | | *EUR GWAS3* | 0.220 | 0.239 | 0.897 | 0.106 | 3.05E-01 | |
| | | | | | | | | *SP GWAS* | - | - | 0.771 | 0.068 | 3.67E-04 | |
| **#35** | rs405858 | 19 | 33106621 | C | T | ANKRD27 | synonymous SNV | **Trans meta** | **-** | **-** | 0.894 | **0.019** | **2.38E-09** | 0.108 |
| | | | | | | | | EAS meta | - | - | 0.872 | 0.024 | 1.70E-08 | |
| | | | | | | | | *HK GWAS* | 0.285 | 0.309 | 0.893 | 0.048 | 1.78E-02 | |
| | | | | | | | | *CC GWAS* | 0.296 | 0.333 | 0.843 | 0.055 | 2.00E-03 | |
| | | | | | | | | *GZ GWAS* | 0.284 | 0.313 | 0.871 | 0.063 | 2.75E-02 | |
| | | | | | | | | *KR IC* | 0.357 | 0.384 | 0.890 | 0.044 | 1.02E-02 | |
| | | | | | | | | *BJ IC* | 0.311 | 0.355 | 0.820 | 0.096 | 3.96E-02 | |
| | | | | | | | | *MC IC* | 0.289 | 0.328 | 0.840 | 0.131 | 1.64E-01 | |
| | | | | | | | | EUR meta | - | - | 0.928 | 0.030 | 1.14E-02 | |
| | | | | | | | | *EUR GWAS1* | 0.665 | 0.694 | 0.877 | 0.089 | 1.43E-01 | |
| | | | | | | | | *EUR GWAS2* | 0.675 | 0.691 | 0.928 | 0.039 | 5.50E-02 | |
| | | | | | | | | *EUR GWAS3* | 0.676 | 0.691 | 0.936 | 0.093 | 4.77E-01 | |
| | | | | | | | | *SP GWAS* | - | - | 0.950 | 0.065 | 4.51E-01 | |
| **#36** | rs3760667 | 19 | 49814832 | C | T | SLC6A16 | intronic | **Trans meta** | **-** | **-** | 0.872 | **0.023** | **2.31E-09** | 0.595 |
| | | | | | | | | EAS meta | - | - | 0.863 | 0.031 | 1.38E-06 | |

| | | | | | | | | | | | | | | |
|---|---|---|---|---|---|---|---|---|---|---|---|---|---|---|
| | | | | | | | | *HK GWAS* | 0.321 | 0.355 | 0.855 | 0.046 | 6.47E-04 | |
| | | | | | | | | *CC GWAS* | 0.266 | 0.288 | 0.888 | 0.056 | 3.27E-02 | |
| | | | | | | | | *GZ GWAS* | 0.331 | 0.370 | 0.847 | 0.060 | 6.16E-03 | |
| | | | | | | | | EUR meta | - | - | 0.884 | 0.035 | 3.73E-04 | |
| | | | | | | | | *EUR GWAS1* | 0.183 | 0.221 | 0.794 | 0.101 | 2.31E-02 | |
| | | | | | | | | *EUR GWAS2* | 0.199 | 0.214 | 0.894 | 0.045 | 1.30E-02 | |
| | | | | | | | | *EUR GWAS3* | 0.228 | 0.233 | 0.972 | 0.103 | 7.80E-01 | |
| | | | | | | | | *SP GWAS* | - | - | 0.865 | 0.079 | 9.03E-02 | |
| **#37** | rs10419308 | 19 | 55739813 | G | A | TMEM86B | intronic | **Trans meta** | **-** | **-** | 0.843 | **0.031** | **3.60E-08** | 0.326 |
| | | | | | | | | EAS meta | - | - | 0.885 | 0.058 | 3.60E-02 | |
| | | | | | | | | *HK GWAS* | 0.055 | 0.059 | 0.945 | 0.095 | 5.52E-01 | |
| | | | | | | | | *CC GWAS* | 0.073 | 0.081 | 0.887 | 0.096 | 2.14E-01 | |
| | | | | | | | | *GZ GWAS* | 0.062 | 0.075 | 0.801 | 0.115 | 5.29E-02 | |
| | | | | | | | | EUR meta | - | - | 0.827 | 0.037 | 2.12E-07 | |
| | | | | | | | | *EUR GWAS1* | 0.189 | 0.209 | 0.878 | 0.103 | 2.05E-01 | |
| | | | | | | | | *EUR GWAS2* | 0.159 | 0.187 | 0.823 | 0.049 | 8.06E-05 | |
| | | | | | | | | *EUR GWAS3* | 0.202 | 0.225 | 0.863 | 0.110 | 1.81E-01 | |
| | | | | | | | | *SP GWAS* | - | - | 0.792 | 0.079 | 1.12E-02 | |
| **#38** | rs6074813 | 20 | 1541752 | G | T | SIRPD;SIRPB1 | intergenic | **Trans meta** | **-** | **-** | 1.120 | **0.021** | **3.23E-08** | 0.930 |
| | | | | | | | | EAS meta | - | - | 1.118 | 0.028 | 8.49E-05 | |
| | | | | | | | | *HK GWAS* | 0.836 | 0.803 | 1.254 | 0.058 | 8.89E-05 | |
| | | | | | | | | *CC GWAS* | 0.807 | 0.789 | 1.108 | 0.064 | 1.08E-01 | |
| | | | | | | | | *GZ GWAS* | 0.819 | 0.808 | 1.085 | 0.075 | 2.75E-01 | |
| | | | | | | | | *KR IC* | 0.768 | 0.761 | 1.042 | 0.051 | 4.57E-01 | |
| | | | | | | | | *BJ IC* | 0.782 | 0.761 | 1.124 | 0.108 | 2.68E-01 | |

| | | | | | |
|---|---|---|---|---|---|
| *MC IC* | 0.825 | 0.807 | 1.124 | 0.153 | 4.34E-01 |
| EUR meta | - | - | 1.122 | 0.030 | 1.01E-04 |
| *EUR GWAS1* | 0.350 | 0.322 | 1.123 | 0.087 | 1.83E-01 |
| *EUR GWAS2* | 0.306 | 0.287 | 1.123 | 0.039 | 3.15E-03 |
| *EUR GWAS3* | 0.388 | 0.382 | 1.022 | 0.092 | 8.12E-01 |
| *SP GWAS* | - | - | 1.170 | 0.064 | 4.68E-02 |

FA: frequency in affected individuals; FU: frequency in unaffected individuals; Trans meta: association results of transethnic meta-analysis across ten SLE genetic data sets; EAS meta: association results of meta-analysis across genetic data sets from six areas in East Asia (HK: Hong Kong; CC: Central China; GZ: Guangzhou; KR: Korea; BJ: Beijing; MC: Malaysia); EUR meta: association results of meta-analysis across four European GWAS for SLE; IC: Immunochip.

**Extended Data Table 5** Putative disease genes identified at each associated locus

| Tag SNP | Region | Gene symbol | Nominal *P* value | Gene closest to the given SNP | eQTL(Westra et al. 2014) | eQTL(Zhernakova et al., 2016) |
|---|---|---|---|---|---|---|
| rs2476601 | chr1:113933371-114456708 | PTPN22 | 4.87E-19 | TRUE | 1 | 1 |
| | | BCL2L15 | 2.59E-04 | FALSE | 0 | 0 |
| rs12093154 | chr1:1152288-1209265 | SDF4 | 4.48E-07 | FALSE | 1 | 1 |
| | | B3GALT6 | 7.04E-04 | FALSE | 1 | 1 |
| rs1016140 | chr1:117035645-117113661 | CD58 | 8.12E-16 | TRUE | 0 | 0 |
| | | AL390066.1 | 8.55E-07 | FALSE | 0 | 0 |
| rs11264750 | chr1:157483167-157567870 | FCRL4 | 7.85E-06 | FALSE | 0 | 0 |
| | | FCRL5 | 0.003112 | TRUE | 1 | 0 |
| rs1801274, | chr1:161551101- | FCGR2A | 2.45E-09 | TRUE | 0 | 0 |
| rs35426045 | 161648444 | FCGR2B | 1.68E-15 | TRUE | 1 | 1 |
| rs12132445 | chr1:170631869-171033906 | MROH9 | 3.12E-09 | TRUE | 0 | 0 |
| rs2205960 | chr1:173152873-173176452 | TNFSF4 | 2.13E-20 | TRUE | 1 | 0 |
| rs1418190 | chr1:173386932-173430501 | LOC101928673 | 0.118541 | TRUE | 0 | 0 |
| rs549669428 | chr1:174128548-175712906 | KIAA0040 | 5.92E-09 | FALSE | 0 | 0 |
| | | RABGAP1L | 1.73E-08 | TRUE | 0 | 0 |
| rs17849501 | chr1:183236953-183706506 | NCF2 | 0.015752 | TRUE | 0 | 0 |
| rs1547624 | chr1:192544857-192549161 | RGS1 | 3.59E-24 | TRUE | 1 | 1 |
| rs34889541 | chr1:198492352-198726545 | PTPRC | 2.74E-18 | TRUE | 1 | 0 |
| rs2297550 | chr1:206643791-206907628 | IKBKE | 4.63E-18 | TRUE | 1 | 1 |
| | | MAPKAPK2 | 1.38E-17 | FALSE | 1 | 0 |
| rs3024505 | chr1:206940947-206945839 | IL10 | 1.82E-15 | TRUE | 0 | 0 |
| rs9782955 | chr1:235824341-236046940 | LYST | 9.28E-15 | TRUE | 1 | 0 |
| rs28411034 | chr1:38259474-38412729 | INPP5B | 6.24E-21 | FALSE | 1 | 1 |
| | | MTF1 | 1.54E-09 | FALSE | 1 | 0 |
| rs6702599 | chr1:67773047-67862583 | IL12RB2 | 1.12E-14 | TRUE | 1 | 1 |
| rs3795310 | chr1:8377886-8877702 | RERE | 0.689557 | TRUE | 1 | 1 |
| rs2322659 | chr2:135722061-136743670 | LOC100507600 | 3.08E-04 | FALSE | 0 | 0 |
| | | RAB3GAP1 | 5.86E-04 | FALSE | 1 | 0 |
| | | MCM6 | 0.010068 | FALSE | 1 | 1 |

| | | | | | | |
|---|---|---|---|---|---|---|
| rs2381401 | chr2:143848931-144525921 | ARHGAP15 | 3.83E-14 | TRUE | 1 | 1 |
| rs1990760 | chr2:163027194-163695240 | IFIH1 | 1.14E-15 | TRUE | 0 | 0 |
| | | GCA | 5.83E-11 | FALSE | 1 | 0 |
| rs9630991 | chr2:191054461-191557492 | NAB1 | 8.13E-12 | FALSE | 1 | 0 |
| | | TMEM194B | 7.93E-07 | TRUE | 1 | 1 |
| | | MFSD6 | 0.047859 | FALSE | 0 | 1 |
| rs11889341 | chr2:191894302-192016322 | STAT4 | 1.35E-17 | TRUE | 1 | 0 |
| rs11679484 | chr2:198256698-199437305 | COQ10B | 4.18E-05 | FALSE | 1 | 0 |
| rs3087243 | chr2:204732509-204738683 | CTLA4 | 5.77E-17 | TRUE | 1 | 0 |
| rs3768792 | chr2:213864429-214017151 | IKZF2 | 3.10E-14 | TRUE | 0 | 1 |
| rs7579944 | chr2:30369807-30383399 | YPEL5 | 4.32E-04 | TRUE | 1 | 0 |
| rs17321999 | chr2:30569525-30575297 | LINC01936 | 0.759929 | TRUE | 0 | 0 |
| rs13385731 | chr2:33661391-33789817 | RASGRP3 | 9.31E-05 | TRUE | 1 | 1 |
| rs6740462 | chr2:65537985-66126649 | SPRED2 | 0.813881 | FALSE | 1 | 1 |
| rs6705628 | chr2:74229840-74329698 | TET3 | 4.53E-11 | TRUE | 1 | 0 |
| rs1131265 | chr3:119013220-119278449 | CD80 | 1.75E-15 | FALSE | 0 | 0 |
| | | ARHGAP31 | 5.73E-08 | FALSE | 1 | 0 |
| | | POGLUT1 | 2.87E-06 | FALSE | 1 | 1 |
| rs564799 | chr3:159631189-159943086 | IL12A-AS1 | 0.990421 | TRUE | 0 | 0 |
| rs10936599 | chr3:169484713-169530774 | ACTRT3 | 6.52E-06 | FALSE | 0 | 0 |
| rs6762714 | chr3:187871072-188608460 | LPP | 0.846121 | TRUE | 1 | 0 |
| rs438613 | chr3:27872379-27875627 | LINC01980 | 8.41E-07 | TRUE | 0 | 0 |
| rs6445972 rs6445975, rs9311676 | chr3:58223233-58419584 | PXK | 6.18E-10 | TRUE | 1 | 1 |
| rs10028805 | chr4:102332443-102995969 | BANK1 | 4.03E-21 | TRUE | 1 | 1 |
| rs4697651 | chr4:10488019-10686489 | CLNK | 1.03E-10 | TRUE | 0 | 0 |

| | | | | | | |
|---|---|---|---|---|---|---|
| rs4690055 | chr4:2463947-2758103 | TNIP2 | 8.86E-20 | TRUE | 0 | 0 |
| | | RNF4 | 0.03771 | FALSE | 1 | 1 |
| rs4690229 | chr4:843064-1020685 | GAK | 2.33E-06 | FALSE | 1 | 0 |
| | | IDUA | 0.001158 | FALSE | 0 | 1 |
| rs7726159 | chr5:1201710-1445545 | TERT | 0.692092 | TRUE | 1 | 0 |
| rs7726414 | chr5:133450402-133747589 | TCF7 | 7.28E-09 | TRUE | 1 | 0 |
| | | UBE2B | 5.70E-04 | FALSE | 1 | 0 |
| | | CDKN2AIPNL | 0.028221 | FALSE | 0 | 0 |
| rs10036748 | chr5:150409506-150473138 | TNIP1 | 5.31E-19 | TRUE | 1 | 0 |
| rs2431697 | chr5:159895275-159914433 | MIR146A | 9.63E-17 | TRUE | 0 | 0 |
| rs6871748 | chr5:35617946-36001130 | IL7R | 1.48E-10 | TRUE | 1 | 0 |
| | | CAPSL | 3.49E-10 | FALSE | 1 | 0 |
| | | RP11-79C6.3 | 0.006426 | FALSE | 0 | 0 |
| | | SPEF2 | 0.019758 | FALSE | 0 | 0 |
| rs548234 | chr6:106534195-106557814 | PRDM1 | 1.92E-08 | TRUE | 1 | 0 |
| rs9387400 | chr6:116575336-116759442 | DSE | 0.084064 | TRUE | 0 | 0 |
| rs2327832 | chr6:137813336-137815531 | OLIG3 | 0.967378 | TRUE | 1 | 0 |
| rs2230926 | chr6:138144810-138204449 | TNFAIP3 | 7.13E-17 | TRUE | 1 | 0 |
| | | ENSG00000235842 | 8.31E-06 | FALSE | 0 | 0 |
| rs6927090 | chr6:292097-351355 | DUSP22 | 4.08E-11 | TRUE | 1 | 0 |
| rs3734266 | chr6:34549107-34897174 | UHRF1BP1 | 0.004 | TRUE | 1 | 1 |
| rs597325 | chr6:90636248-91006627 | BACH2 | 1.16E-12 | TRUE | 1 | 1 |
| rs12531711, rs4728142, rs729302 | chr7:128577666-128695198 | IRF5 | 3.71E-17 | TRUE | 1 | 1 |
| rs849142 | chr7:27870192-28281515 | JAZF1 | 4.24E-04 | TRUE | 1 | 1 |
| rs2366293 | chr7:49890017-50160925 | ZPBP | 0.998704 | TRUE | 0 | 0 |
| rs4917014 | chr7:50343720-50472799 | IKZF1 | 3.34E-14 | TRUE | 1 | 0 |
| rs13238909 | chr7:66767608-66786513 | STAG3L4 | 0.959175 | TRUE | 0 | 0 |
| rs73135369, rs117026326 | chr7:73901807-74016931 | NCF1 | 0.00135 | FALSE | 1 | 0 |

| | | | | | | |
|---|---|---|---|---|---|---|
| rs11773745 | chr7:75162621-75368280 | HIP1 | 0.608016 | TRUE | 1 | 1 |
| rs12680762, rs2736340 | chr8:11225911-11438851 | BLK | 7.56E-19 | TRUE | 1 | 1 |
| | | LINC00208 | 8.70E-10 | FALSE | 0 | 0 |
| | | C8orf12 | 0.027281 | FALSE | 0 | 0 |
| rs2445610 | chr8:128698588-128746213 | CASC11 | 1.66E-05 | TRUE | 0 | 0 |
| rs7815944 | chr8:129417515-129440162 | LINC00824 | 0.002272 | TRUE | 0 | 0 |
| rs13260060 | chr8:71021997-71316040 | NCOA2 | 2.29E-08 | TRUE | 0 | 0 |
| rs2428 | chr8:8640864-9025646 | MFHAS1 | 4.38E-13 | TRUE | 1 | 1 |
| | | ERI1 | 7.24E-04 | FALSE | 1 | 0 |
| rs1405209 | chr9:102584137-102629173 | NR4A3 | 1.85E-08 | TRUE | 0 | 0 |
| rs4978037 | chr9:21077104-21385396 | IFNA21 | 9.61E-13 | FALSE | 0 | 0 |
| | | IFNA7 | 1.74E-12 | FALSE | 0 | 0 |
| | | IFNB1 | 4.11E-07 | FALSE | 1 | 0 |
| | | IFNA2 | 5.61E-04 | FALSE | 0 | 0 |
| | | IFNA5 | 0.001184 | FALSE | 0 | 0 |
| rs7097397 | chr10:49892921-50191001 | WDFY4 | 2.60E-20 | TRUE | 1 | 0 |
| rs4948496 | chr10:63661059-63856703 | ARID5B | 7.93E-07 | TRUE | 1 | 1 |
| rs4745876 | chr10:64133951-64431771 | ZNF365 | 0.999889 | TRUE | 1 | 0 |
| rs10999979 | chr10:73156691-73575702 | CDH23 | 1.68E-11 | TRUE | 1 | 0 |
| rs11603023 | chr11:118477155-118550399 | TREH | 0.999992 | TRUE | 0 | 0 |
| rs4639966 | chr11:118620034-118661858 | DDX6 | 0.701517 | FALSE | 0 | 1 |
| rs1128334, rs7941765 | chr11:128328656-128457437 | ETS1 | 2.18E-12 | TRUE | 1 | 0 |
| rs2732552 | chr11:34937376-35042138 | PDHX | 0.914892 | TRUE | 1 | 0 |
| rs12802200, rs1061502 | chr11:565660-626078 | IRF7 | 1.34E-10 | FALSE | 1 | 1 |
| rs1308020, rs2009453, rs494003 | chr11:65365226-65564690 | SIPA1 | 2.33E-23 | FALSE | 1 | 1 |
| | | RELA | 2.22E-17 | FALSE | 1 | 0 |
| | | MAP3K11 | 3.52E-17 | FALSE | 1 | 1 |
| | | PCNXL3 | 3.12E-05 | TRUE | 0 | 1 |

| | | | | | | |
|---|---|---|---|---|---|---|
| rs10750836 | chr11:68816350-69490184 | TPCN2 | 0.109298 | TRUE | 0 | 0 |
| rs11235604 | chr11:72335731-72922527 | ATG16L2 | 0.038465 | TRUE | 0 | 0 |
| rs10774625 | chr12:111843752-112947717 | SH2B3 | 2.58E-19 | FALSE | 1 | 1 |
| | | TRAFD1 | 8.40E-16 | FALSE | 1 | 0 |
| | | PTPN11 | 0.006526 | FALSE | 0 | 0 |
| rs7975703 | chr12:120884241-121442296 | UNC119B | 0.39747 | FALSE | 1 | 1 |
| | | COQ5 | 0.901841 | FALSE | 1 | 1 |
| | | MLEC | 0.987851 | FALSE | 0 | 1 |
| | | CABP1 | 0.999574 | TRUE | 1 | 1 |
| rs10845606 | chr12:12813825-12849141 | GPR19 | 0.975997 | TRUE | 1 | 0 |
| rs1385374 | chr12:129277739-129308528 | SLC15A4 | 6.91E-16 | TRUE | 1 | 0 |
| rs1885889 | chr13:100153671-100215645 | TM9SF2 | 0.974693 | FALSE | 1 | 0 |
| rs12874404 | chr13:108903588-108959385 | TNFSF13B | 5.26E-14 | TRUE | 1 | 0 |
| rs7329174 | chr13:41506056-42045018 | ELF1 | 3.23E-17 | TRUE | 1 | 0 |
| | | WBP4 | 7.15E-06 | FALSE | 0 | 0 |
| | | C13orf15 | 5.72E-04 | FALSE | 0 | 0 |
| rs76725306 | chr13:50018429-50265623 | SETDB2 | 3.26E-10 | FALSE | 1 | 0 |
| rs76725306 | chr13:50018429-50265623 | ARL11 | 3.24E-05 | FALSE | 1 | 0 |
| rs12148050 | chr14:103243813-103377837 | TRAF3 | 1.51E-20 | TRUE | 1 | 1 |
| rs2841280 | chr14:105391153-105444694 | PLD4 | 7.73E-12 | TRUE | 0 | 1 |
| rs4902562 | chr14:68286496-69196935 | RAD51B | 3.83E-08 | TRUE | 1 | 0 |
| rs117518546 | chr14:105968621-106289176 | IGHG1 | 6.15E-10 | TRUE | 0 | 0 |
| rs8035957, rs954431 | chr15:38780304-38857776 | RASGRP1 | 5.59E-07 | TRUE | 1 | 0 |
| rs2289583 | chr15:75105057-75343067 | SCAMP2 | 4.83E-15 | FALSE | 1 | 1 |
| | | ULK3 | 2.93E-05 | FALSE | 1 | 1 |
| | | PPCDC | 0.001472 | FALSE | 0 | 0 |
| rs869310 | chr15:77334178-78370066 | TBC1D2B | 1.10E-10 | FALSE | 1 | 0 |
| | | HMG20A | 0.019458 | FALSE | 1 | 0 |
| rs12599402 | chr16:11038345-11276046 | CLEC16A | 6.20E-13 | TRUE | 1 | 0 |

| | | | | | | |
|---|---|---|---|---|---|---|
| rs4592664 | chr16:23847322-24231932 | PRKCB | 8.38E-10 | TRUE | 1 | 1 |
| rs1143679 | chr16:31157884-31737100 | ITGAM | 4.21E-05 | TRUE | 0 | 0 |
| | | ITGAX | 6.41E-06 | FALSE | 0 | 1 |
| rs223881 | chr16:57392684-57449974 | CCL22 | 2.70E-13 | TRUE | 0 | 0 |
| | | CCL17 | 2.65E-10 | FALSE | 1 | 0 |
| rs2731783 | chr16:58191811-58328951 | CSNK2A2 | 0.167448 | TRUE | 1 | 0 |
| rs1170426 | chr16:68573661-68732971 | ZFP90 | 0.988934 | TRUE | 1 | 1 |
| rs11644034, rs2934498, rs2280381 | chr16:85932409-85956197 | IRF8 | 3.90E-17 | TRUE | 1 | 0 |
| rs34562254 | chr17:16826229-16875402 | TNFRSF13B | 3.44E-16 | TRUE | 0 | 0 |
| rs8079075 | chr17:37885409-38083854 | IKZF3 | 4.55E-19 | TRUE | 1 | 0 |
| | | GSDMB | 1.22E-13 | FALSE | 1 | 0 |
| | | ORMDL3 | 1.07E-10 | FALSE | 1 | 0 |
| rs9899849 | chr17:7138350-7464925 | ACAP1 | 8.62E-21 | FALSE | 1 | 1 |
| rs930297 | chr17:73232695-73401790 | GRB2 | 1.04E-19 | TRUE | 1 | 1 |
| | | MIF4GD | 9.19E-17 | FALSE | 1 | 1 |
| | | GGA3 | 1.07E-12 | FALSE | 1 | 1 |
| | | SLC25A19 | 1.50E-12 | FALSE | 1 | 0 |
| rs763361 | chr18:67068284-67624160 | CD226 | 2.96E-16 | TRUE | 1 | 1 |
| rs2304256 | chr19:10461209-10514271 | TYK2 | 4.31E-18 | TRUE | 1 | 1 |
| rs2384991 | chr19:18367909-18385319 | KIAA1683 | 0.789195 | TRUE | 0 | 1 |
| rs13344313 | chr19:18501955-18508415 | LRRC25 | 1.06E-11 | TRUE | 1 | 1 |
| rs405858 | chr19:33072096-33169206 | ANKRD27 | 0.601002 | TRUE | 0 | 1 |
| rs3760667 | chr19:49792897-50192286 | CD37 | 2.07E-20 | FALSE | 1 | 1 |
| | | FLT3LG | 9.69E-17 | FALSE | 0 | 0 |
| | | IRF3 | 3.14E-09 | FALSE | 1 | 0 |
| | | RRAS | 4.68E-05 | FALSE | 1 | 0 |
| | | FCGRT | 0.001201 | FALSE | 1 | 0 |
| | | ALDH16A1 | 0.001642 | FALSE | 1 | 0 |
| rs2305772 | chr19:52022779-52035110 | SIGLEC6 | 2.11E-05 | TRUE | 1 | 1 |
| rs10419308 | | PPP6R1 | 6.18E-16 | FALSE | 1 | 1 |

| | | | | | | |
|---|---|---|---|---|---|---|
| | chr19:55738002-55770363 | TMEM86B | 0.003115 | TRUE | 1 | 0 |
| rs6074813 | chr20:1451386-1638425 | SIRPB2 | 8.45E-15 | FALSE | 0 | 0 |
| | | SIRPG | 8.97E-15 | FALSE | 1 | 0 |
| | | SIRPB1 | 1.75E-06 | FALSE | 0 | 0 |
| | | SIRPD | 0.005634 | FALSE | 1 | 0 |
| rs4810485 | chr20:44650329-44758502 | CD40 | 2.15E-17 | TRUE | 1 | 1 |
| | | NCOA5 | 0.005242 | FALSE | 1 | 0 |
| rs463426 | chr22:21771693-21805752 | HIC2 | 0.928278 | TRUE | 0 | 0 |
| rs7444 | chr22:21903736-21984353 | UBE2L3 | 0.185039 | TRUE | 0 | 1 |
| rs61616683 | chr22:39619364-39781593 | PDGFB | 2.05E-04 | FALSE | 1 | 0 |

**Extended Data Table 6** Pathway enrichment analysis based on putative SLE genes

| Name | Source | *P*-value | FDR |
|---|---|---|---|
| Cytokine Signaling in Immune system | REACTOME | 3.05E-12 | 2.18E-08 |
| Interferon alpha/beta signaling | REACTOME | 2.23E-11 | 7.98E-08 |
| Toll-like receptor signaling pathway | KEGG | 1.34E-10 | 2.91E-07 |
| Measles | KEGG | 1.98E-10 | 2.91E-07 |
| RIG-I/MDA5 mediated induction of IFN-alpha/beta pathways | REACTOME | 2.03E-10 | 2.91E-07 |
| RIG-I-like receptor signaling pathway | KEGG | 5.83E-10 | 6.54E-07 |
| TRAF6 mediated IRF7 activation | REACTOME | 6.41E-10 | 6.54E-07 |
| Regulation of IFNA signaling | REACTOME | 2.25E-09 | 1.80E-06 |
| Cytokine-cytokine receptor interaction | KEGG | 2.27E-09 | 1.80E-06 |
| Hepatitis B | KEGG | 5.93E-09 | 4.24E-06 |
| Herpes simplex infection | KEGG | 1.06E-08 | 6.91E-06 |
| Jak-STAT signaling pathway | KEGG | 1.47E-08 | 8.76E-06 |
| Autoimmune thyroid disease | KEGG | 2.11E-08 | 1.16E-05 |
| Hepatitis C | KEGG | 2.44E-08 | 1.25E-05 |
| Cytosolic DNA-sensing pathway | KEGG | 9.71E-08 | 4.63E-05 |
| TRAF3-dependent IRF activation pathway | REACTOME | 2.17E-07 | 9.71E-05 |
| Innate Immune System | REACTOME | 3.08E-07 | 1.30E-04 |
| NOD-like receptor signaling pathway | KEGG | 3.55E-07 | 1.41E-04 |
| Influenza A | KEGG | 4.23E-07 | 1.59E-04 |
| Interferon Signaling | REACTOME | 1.96E-06 | 6.68E-04 |
| Signal regulatory protein (SIRP) family interactions | REACTOME | 4.17E-06 | 1.35E-03 |
| Factors involved in megakaryocyte development and platelet production | REACTOME | 1.16E-05 | 3.33E-03 |
| Adaptive Immune System | REACTOME | 1.25E-05 | 3.45E-03 |
| Activation of IRF3/IRF7 mediated by TBK1/IKK epsilon | REACTOME | 1.72E-05 | 4.40E-03 |
| Ovarian tumor domain proteases | REACTOME | 1.87E-05 | 4.60E-03 |
| Signaling by Interleukins | REACTOME | 2.05E-05 | 4.88E-03 |
| Osteoclast differentiation | KEGG | 2.18E-05 | 5.03E-03 |
| NF-kappa B signaling pathway | KEGG | 2.27E-05 | 5.08E-03 |
| Natural killer cell mediated cytotoxicity | KEGG | 2.57E-05 | 5.57E-03 |
| Toll Like Receptor 3 (TLR3) Cascade | REACTOME | 2.98E-05 | 5.91E-03 |
| MyD88-independent TLR3/TLR4 cascade | REACTOME | 2.98E-05 | 5.91E-03 |
| TRIF-mediated TLR3/TLR4 signaling | REACTOME | 2.98E-05 | 5.91E-03 |
| Intestinal immune network for IgA production | KEGG | 7.49E-05 | 1.28E-02 |
| Activated TLR4 signalling | REACTOME | 7.78E-05 | 1.29E-02 |
| Epstein-Barr virus infection | KEGG | 8.80E-05 | 1.43E-02 |
| Toll Like Receptor 4 (TLR4) Cascade | REACTOME | 1.38E-04 | 2.10E-02 |
| Hemostasis | REACTOME | 1.87E-04 | 2.57E-02 |
| Ras signaling pathway | KEGG | 2.05E-04 | 2.75E-02 |
| Tuberculosis | KEGG | 2.08E-04 | 2.75E-02 |
| Signaling by the B Cell Receptor (BCR) | REACTOME | 2.74E-04 | 3.56E-02 |

| | | | |
|---|---|---|---|
| Negative regulators of RIG-I/MDA5 signaling | REACTOME | 2.93E-04 | 3.74E-02 |
| Primary immunodeficiency | KEGG | 3.26E-04 | 3.89E-02 |
| T cell receptor signaling pathway | KEGG | 3.26E-04 | 3.89E-02 |
| Downstream signaling events of B Cell Receptor (BCR) | REACTOME | 4.26E-04 | 4.99E-02 |
| B cell receptor signaling pathway | KEGG | 4.37E-04 | 4.99E-02 |

**Extended Data Table 7** Summary of fine-mapping results based on ancestry-dependent and trans-ancestral group methods

| Tag SNP | Locus | Region | Number of variants in 95% credible set | | | Putative casual variant[1] | Posterior Probability[2] | r2 with tag SNP in EAS | r2 with tag SNP in EUR | Consequence[3] |
|---|---|---|---|---|---|---|---|---|---|---|
| | | | Chinese GWAS | European GWAS | Trans-ancestry | | | | | |
| **rs1801274** | FCGR2A | chr1:161457416-161483666 | 14 | 3 | 3 | rs6671847 | 0.595 | 0.93 | 0.89 | intron_variant,NMD_transcript_variant |
| | | | | | | rs7551957 | 0.208 | 0.67 | 0.86 | intergenic_variant |
| | | | | | | rs12139150 | 0.184 | 0.67 | 0.86 | upstream_gene_variant |
| **rs2205960** | TNFSF4 | chr1:173097989-173297704 | 21 | 9 | 1 | rs2205960 | 0.957 | 1.00 | 1.00 | intergenic_variant |
| **rs2297550** | IKBKE | chr1:206620600-206707382 | 10 | 1 | 1 | rs2297550 | 0.997 | 1.00 | 1.00 | regulatory_region_variant TF_binding_site_variant |
| **rs7579944** | LBH | chr2:30426413-30456342 | 7 | 7 | 5 | rs7579944 | 0.285 | 1.00 | 1.00 | intergenic_variant |
| | | | | | | rs906868 | 0.193 | 0.94 | 0.97 | regulatory_region_variant |
| | | | | | | rs12473138 | 0.186 | 0.94 | 0.93 | intergenic_variant |
| | | | | | | rs10173253 | 0.178 | 0.99 | 0.97 | intergenic_variant |
| | | | | | | rs10175798 | 0.152 | 0.92 | 0.92 | upstream_gene_variant |
| **rs13385731** | RASGRP3 | chr2:33668178-33703389 | 2 | 73 | 2 | rs13385731 | 0.631 | 1.00 | 1.00 | regulatory_region_variant TF_binding_site_variant |
| | | | | | | rs13425999 | 0.369 | 0.99 | 0.98 | regulatory_region_variant |
| **rs1990760** | IFIH1 | chr2:162997703-163414643 | 47 | 3 | 3 | rs2111485 | 0.538 | 0.81 | 0.89 | regulatory_region_variant |
| | | | | | | rs1990760 | 0.281 | 1.00 | 1.00 | missense_variant |
| | | | | | | rs2075302 | 0.152 | 0.61 | 0.40 | intron_variant |
| **rs11889341 (Primary signal)** | STAT4 | chr2:191874667-191971565 | 5 | 8 | 4 | rs11889341 | 0.804 | 1.00 | 1.00 | intron_variant |
| | | | | | | rs4853458 | 0.062 | 0.85 | 0.99 | intron_variant |
| | | | | | | rs7574865 | 0.054 | 0.85 | 0.98 | intron_variant |
| | | | | | | rs4274624 | 0.045 | 0.84 | 0.98 | intron_variant |

| | | | | | | | | | | |
|---|---|---|---|---|---|---|---|---|---|---|
| **rs16833239 (Secondary)** | STAT4 | condition on rs11889341 | 5 | 9 | 3 | rs16833239 | 0.594 | 1.00 | 1.00 | intron_variant |
| | | | | | | rs7594501 | 0.309 | 0.94 | 1.00 | intron_variant |
| | | | | | | rs140341964 | 0.061 | 1.00 | 0.37 | intron_variant |
| **rs6762714** | LPP | chr3:188450118-188499238 | 26 | 15 | 4 | rs12486509 | 0.342 | 0.98 | 0.99 | intron_variant |
| | | | | | | rs6762714 | 0.271 | 1.00 | 1.00 | intron_variant |
| | | | | | | rs1040029 | 0.227 | 0.96 | 0.84 | intron_variant |
| | | | | | | rs3830584 | 0.144 | 0.97 | 0.85 | regulatory_region_variant TF_binding_site_variant |
| **rs7726159** | TERT | chr5:1232491-1356450 | 1 | 71 | 1 | rs2736100 | 1.000 | 0.82 | 0.52 | intron_variant,NMD_transcript_variant |
| **rs10036748** | TNIP1 | chr5:150430000-150462638 | 5 | 4 | 3 | rs7708392 | 0.403 | 0.99 | 0.99 | intron_variant,regulatory_region_variant |
| | | | | | | rs6889239 | 0.340 | 0.99 | 0.99 | intron_variant,regulatory_region_variant |
| | | | | | | rs10036748 | 0.231 | 1.00 | 1.00 | intron_variant,regulatory_region_variant |
| **rs2431697 (Primary)** | PTTG1-MIR146A | chr5:159868247-159966924 | 5 | 7 | 1 | rs2431697 | 0.999 | 1.00 | 1.00 | intergenic_variant |
| **rs57095329 (Secondary)** | PTTG1-MIR146A | condition on rs2431697 | 7 | 33 | 4 | rs57095329 | 0.355 | 1.00 | 1.00 | regulatory_region_variant TF_binding_site_variant |
| | | | | | | rs58585875 | 0.304 | 0.99 | 0.97 | upstream_gene_variant regulatory_region_variant |
| | | | | | | rs73318382 | 0.268 | 0.99 | 0.97 | upstream_gene_variant regulatory_region_variant |
| | | | | | | rs2277920 | 0.064 | 0.90 | 0.97 | intron_variant regulatory_region_variant |
| **rs548234** | PRDM1-ATG5 | chr6:106564400-106599710 | 13 | 9 | 5 | rs7768653 | 0.279 | 0.85 | 0.61 | intergenic_variant |
| | | | | | | rs4134466 | 0.265 | 0.86 | 0.61 | intergenic_variant,regulatory_region_variant |

| | | | | | | | | | | |
|---|---|---|---|---|---|---|---|---|---|---|
| | | | | | | rs2005969 | 0.223 | 0.86 | 0.61 | intergenic_variant |
| | | | | | | rs1883231 | 0.170 | 0.85 | 0.61 | intergenic_variant |
| | | | | | | rs6937876 | 0.021 | 0.71 | 0.56 | intergenic_variant |
| **rs2736340** | FAM167A-BLK | chr8:11285384-11400944 | 17 | 11 | 5 | rs2618473 | 0.451 | 0.99 | 0.91 | intergenic_variant |
| | | | | | | rs2061831 | 0.226 | 0.95 | 1.00 | intergenic_variant,regulatory_region_variant |
| | | | | | | rs2618444 | 0.126 | 0.94 | 1.00 | intergenic_variant |
| | | | | | | rs2409780 | 0.124 | 0.95 | 1.00 | intergenic_variant,regulatory_region_variant |
| | | | | | | rs2736338 | 0.033 | 1.00 | 1.00 | intergenic_variant |
| **rs7097397** | WDFY4 | chr10:49984302-50138755 | 67 | 13 | 1 | rs7097397 | 0.993 | 1.00 | 1.00 | missense_variant |
| **rs12599402** | CLEC16A | chr16:11017058-11464639 | 3 | 81 | 3 | rs12599402 | 0.441 | 1.00 | 1.00 | intron_variant |
| | | | | | | rs12917656 | 0.379 | 1.00 | 1.00 | intron_variant |
| | | | | | | rs12917716 | 0.180 | 0.99 | 1.00 | intron_variant |

[1]Putative casual variants in each locus refer to a list of variants that are included in the 95% credible set for the trans-ethnic fine mapping. [2]Posterior probability for the trans-ethnic fine mapping is estimated by PAINTOR under the assumption of single causal variant per locus. If multiple signals were present within a locus, a preliminary round of conditional analysis was performed. [3]The functional consequence of each variant is annotated by using VEP (https://www.ensembl.org/vep).

**Extended Data Table 8** SLE-associated loci with heterogenity between East Asian and European populations

| SNP | Chr | Pos | Risk allele | GENE | Function | East Asian population | | | | European population | | | | Cochran's Q-test |
|---|---|---|---|---|---|---|---|---|---|---|---|---|---|---|
| | | | | | | Risk allele freq | OR | SE | P | Risk allele freq | OR | SE | P | |
| rs12132445 | 1 | 170811799 | A | PRRX1-MROH9 | intergenic | 0.616 | **1.17** | 0.03 | 6.75E-08 | 0.458 | **1.00** | 0.03 | 9.67E-01 | 1.05E-04 |
| rs9387400 | 6 | 116694120 | C | DSE | intronic | 0.052 | **1.35** | 0.05 | 3.14E-08 | 0.627 | **1.06** | 0.03 | 2.96E-02 | 1.17E-04 |
| rs4917014 | 7 | 50305863 | T | IKZF1 | intergenic | 0.709 | **1.33** | 0.03 | 5.18E-29 | 0.679 | **1.16** | 0.03 | 1.34E-06 | 4.02E-04 |
| rs11773745 | 7 | 75171438 | A | HIP1 | intronic | 0.676 | **1.24** | 0.03 | 2.52E-11 | 0.266 | **1.00** | 0.03 | 8.55E-01 | 3.77E-06 |
| rs2841280 | 14 | 105393556 | C | PLD4 | missense | 0.575 | **1.21** | 0.03 | 6.12E-11 | 0.468 | **1.05** | 0.03 | 7.10E-02 | 5.15E-04 |
| rs4592664 | 16 | 23890735 | T | PRKCB | intronic | 0.766 | **1.17** | 0.03 | 2.37E-06 | 0.804 | **0.96** | 0.03 | 1.90E-01 | 2.23E-05 |
| rs9899849 | 17 | 7234983 | A | NEURL4-ACAP1 | intergenic | 0.293 | **0.97** | 0.03 | 3.32E-01 | 0.225 | **1.21** | 0.03 | 1.10E-08 | 1.72E-06 |
| rs34562254 | 17 | 16842991 | A | TNFRSF13B | missense | 0.455 | **1.18** | 0.03 | 2.88E-08 | *0.107* | **1.01** | 0.04 | 7.57E-01 | 3.58E-04 |
| rs2304256 | 19 | 10475652 | C | TYK2 | missense | 0.427 | **0.99** | 0.02 | 7.58E-01 | 0.713 | **1.23** | 0.03 | 8.12E-11 | 2.25E-07 |

Risk allele frequency in East Asians is estimated by using the 3,324 HK controls, and the risk allele frequency in Europeans is estimated by using the 5,379 controls in the EUR GWAS 2 cohort.

**Extended Data Table 9** Twenty-seven immune-unrelated phenotypes studied in European populations used to compare with East Asian SLE association signals in colocalization analyses

| Index | Traits | Consortium |
|---|---|---|
| 1 | Coronary Artery Disease (CAD) | CARDIoGRAM |
| 2 | Forearm bone mineral density (FABMD) | GEFOS |
| 3 | Femoral neck bone mineral density (FNBMD) | GEFOS |
| 4 | Lumbar spine bone mineral density (LSBMD) | GEFOS |
| 5 | Extreme BMI | GIANT |
| 6 | Extreme Height | GIANT |
| 7 | Extreme waist hip ratio (WHR) | GIANT |
| 8 | HDL | GLGC |
| 9 | LDL | GLGC |
| 10 | Total Cholesterol (TC) | GLGC |
| 11 | Triglycerides (TG) | GLGC |
| 12 | Fasting Glucose (FG) | MAGIC |
| 13 | Acetoacetate | MAGNETIC |
| 14 | Acetate | MAGNETIC |
| 15 | Alanine | MAGNETIC |
| 16 | Albumin | MAGNETIC |
| 17 | Anorexia | PGC |
| 18 | Autism Spectrum Disorder (ASD) | PGC |
| 19 | Bipolar Disorder (BIP) | PGC |
| 20 | Major depressive disorder (MDD) | PGC |
| 21 | Schizophrenia | PGC |
| 22 | Educational attainment | SSGAC |
| 23 | Neuroticism | SSGAC |
| 24 | Insomnia | SSGAC |
| 25 | Smoking/cigarettes per day | TAG |
| 26 | Aging | Aging |
| 27 | Childhood obesity | EGG |

**Extended Data Table 10** Groups of putative SLE genes based on heterogeneity in effect-size estimates

| Category | #Loci | #Genes | Genes identified at the disease loci |
|---|---|---|---|
| Ancestry-shared loci with CQ-test *P* > 0.05 (**Category 1**) | 79 | 120 | SDF4,B3GALT6,FCGR2A,LOC101928673,KIAA0040,RABGAP1L,PTPRC,IKBKE,MAPKAPK2,LYST,INPP5B,MTF1,IL12RB2,RERE,WDFY4,ARID5B,ZNF365,TREH,ETS1,PDHX,IRF7,UNC119B,COQ5,MLEC,CABP1,SLC15A4,TM9SF2,SETDB2,ARL11,TRAF3,RASGRP1,SCAMP2,ULK3,PPCDC,TBC1D2B,HMG20A,CLEC16A,CCL22,CCL17,PRSS54,CSNK2A2,ZFP90,IRF8,GRB2,MIF4GD,GGA3,SLC25A19,CD226,KIAA1683,LRRC25,ANKRD27,CD37,FLT3LG,IRF3,RRAS,FCGRT,ALDH16A1,SIGLEC6,PPP6R1,TMEM86B,ARHGAP15,IFIH1,GCA,NAB1,TMEM194B,MFSD6,COQ10B,CTLA4,YPEL5,LINC01936,TET3,SIRPB2,SIRPG,SIRPB1,SIRPD,CD40,NCOA5,UBE2L3,CD80,ARHGAP31,POGLUT1,IL12A-AS1,ACTRT3,LPP,AC098973.1 ,PXK,BANK1,CLNK,TNIP2,RNF4,GAK,IDUA,TERT,TCF7,UBE2B,CDKN2AIPNL,TNIP1,MIR3142HG,IL7R,CAPSL,RP11-79C6.3,SPEF2,PRDM1,UHRF1BP1,BACH2,ZPBP,STAG3L4,NCF1,BLK,LINC00208,C8orf12,RP11-89M16.1,MFHAS1,ERI1,NR4A3,IFNA21,IFNA7,IFNB1,IFNA2,IFNA5 |
| Putative ancestry-heterogenous loci with CQ-test *P* < 0.05 and FDR adjusted CQ-test *P* > 0.05 (**Category 2**) | 20 | 22 | CD58,AL390066.1,TNFSF4,RGS1,IL10,DDX6,SIPA1,RELA,MAP3K11,PCNXL3,GPR19,RAD51B,STAT4,IKZF2,RASGRP3,SPRED2,HIC2,PDGFB,TNFAIP3,IRF5,CASC11 ,NCOA2 |
| Ancestry-heterogenous loci with adjusted CQ-test *P* < 0.05 (**Category 3**) | 9 | 9 | PRKCB,TYK2,PLD4,TNFRSF13B,IKZF1,HIP1,DSE,ACAP1,PRRX1 |
| Disease loci with risk allele monomorphic in one of the two ancestries (**Category 4**) | 7 | 12 | PTPN22,BCL2L15,NCF2,SH2B3,TRAFD1,PTPN11,TNFSF13B,IKZF3,GSDMB,ORMDL3,ATG16L2, IGHG1 |
